# Rapid and efficient analysis of 20,000 RNA-seq samples with Toil

**DOI:** 10.1101/062497

**Authors:** John Vivian, Arjun Rao, Frank Austin Nothaft, Christopher Ketchum, Joel Armstrong, Adam Novak, Jacob Pfeil, Jake Narkizian, Alden D. Deran, Audrey Musselman-Brown, Hannes Schmidt, Peter Amstutz, Brian Craft, Mary Goldman, Kate Rosenbloom, Melissa Cline, Brian O’Connor, Megan Hanna, Chet Birger, W. James Kent, David A. Patterson, Anthony D. Joseph, Jingchun Zhu, Sasha Zaranek, Gad Getz, David Haussler, Benedict Paten

**Affiliations:** Computational Genomics Lab, Genomics Institute, University of California Santa Cruz, Santa Cruz, CA, USA.; AMP Lab, University of California Berkeley, Berkeley, CA, USA.; UC Berkeley ASPIRE Lab, Berkeley, CA, USA.; Curoverse, Somerville, MA, USA.; Broad Institute of Harvard and Massachusetts Institute of Technology (MIT), Cambridge, Massachusetts, USA.

## Abstract

Toil is portable, open-source workflow software that supports contemporary workflow definition languages and can be used to securely and reproducibly run scientific workflows efficiently at large-scale. To demonstrate Toil, we processed over 20,000 RNA-seq samples to create a consistent meta-analysis of five datasets free of computational batch effects that we make freely available. Nearly all the samples were analysed in under four days using a commercial cloud cluster of 32,000 preemptable cores.

---

Contemporary genomic datasets contain tens of thousands of samples and petabytes of sequencing data^1,2,3^. Genomic processing pipelines can consist of dozens of individual steps, each with their own set of parameters^4,5^. As a result of this size and complexity, computational resource limitations and reproducibility are becoming a major concern within genomics. In response to these interrelated issues, we have created Toil.

## Reproducible Workflows

To support the sharing of scientific workflows, Toil is the first software to execute Common Workflow Language (CWL, **Supplementary Note 7**) and provide draft support for Workflow Description Language (WDL), both burgeoning standards for scientific workflows^6,7^. A workflow is composed of a set of tasks, or jobs, that are orchestrated by specification of a set of dependencies that map the inputs and outputs between jobs. In addition to CWL and draft WDL support, Toil provides a Python API that allows workflows to be declared statically, or generated dynamically, so that jobs can define further jobs as needed (**Supplementary Note 1**). The jobs defined in either CWL or Python can consist of Docker containers, which permit sharing of a program without requiring individual tool installation or configuration within a specific environment. Open-source workflows that invoke containers can therefore be run precisely and reproducibly, regardless of environment. We provide a repository of workflows as examples^8^. Toil also integrates with Apache Spark^9^ (**Supplementary Note 6, Supplementary Fig. 4**), and can be used to rapidly create containerized Spark clusters within the context of a larger workflow^10^.

**Figure 1.**
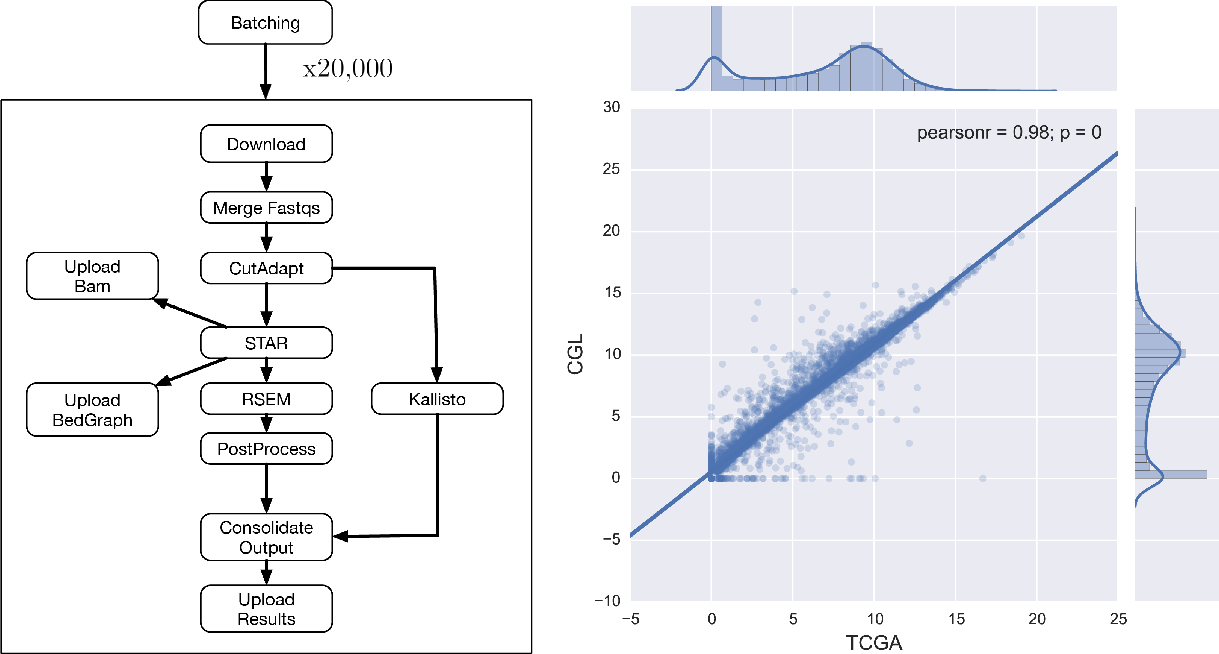
**(Left)** A dependency graph of the RNA-seq pipeline we developed (called CGL). CutAdapt was used to remove extraneous adapters, STAR was used for alignment and read coverage, and RSEM and Kallisto were used to produce quantification data. **(Right)** A scatter plot showing the Pearson correlation between the results of the TCGA best-practices pipeline and the CGL pipeline. 10,000 randomly selected sample/gene pairs were subset from the entire TCGA cohort and the normalized counts were plot against each other; this process was repeated 5 times with no change in Pearson correlation. The unit for counts is: log2(norm_counts+1).

## Portability

Toil runs within multiple cloud environments including Amazon Web Services (AWS), Microsoft Azure, Google Cloud, OpenStack, and within traditional high performance computing (HPC) environments running GridEngine or Slurm and distributed systems running Apache Mesos^11–13^. Toil also runs on a single machine, such as a laptop or workstation, to allow for interactive development, and can be installed with a single command. This portability stems from pluggable backend APIs for machine provisioning, job scheduling, and file management (**Supplementary Note 2**). Implementation of these APIs facilitates straight-forward extension of Toil to new compute environments. Toil manages intermediate files and checkpointing through a job store, which can be an object store like S3 or a network file-system such as NFS. The flexibility of the backend APIs allow a single script to be run on any supported compute environment, paired with any job store, without requiring any modifications to the source code.

**Figure 2.**
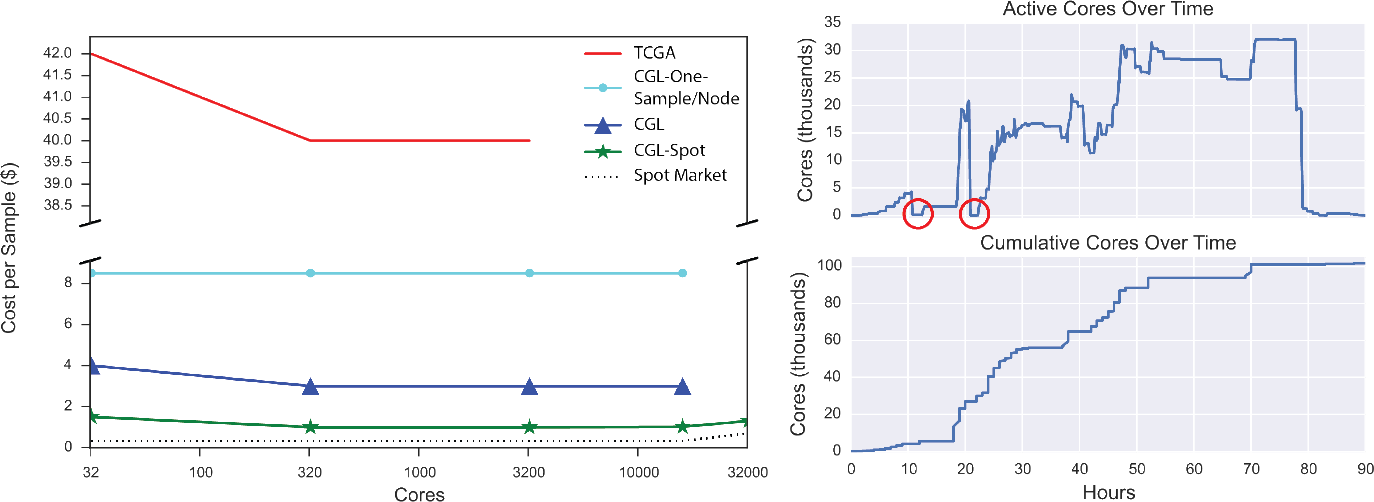
**(Left)** Scaling tests were run to ascertain the price per sample at varying cluster sizes for the different analysis methods. TCGA (red) shows the cost of running the TCGA best practice pipeline as re-implemented as a Toil workflow (for comparison). CGL-One-Sample/Node (cyan) shows the cost of running the revised Toil pipeline, one sample per node. CGL (blue) denotes the pipeline running samples across many nodes. CGL-Spot (green) is as CGL, but denotes the pipeline run on the Amazon spot market. The slight rise in cost per sample at 32,000 cores was due to a couple factors: aggressive instance provisioning directly affected the spot price (dotted line), and saving bam and bedGraph files for each sample. **(Right)** Tracking number of cores during the recompute. The two red circles indicate where all worker nodes were terminated and subsequently restarted shortly thereafter.

## Scale and Efficiency

To demonstrate the scalability and efficiency of Toil, we used a single Toil script to recompute read-mappings and isoform expression values for 19,952 samples spanning 4 major studies — three cancer-related: The Cancer Genome Atlas (TCGA)^1^, Therapeutically Applicable Research To Generate Effective Treatments (TARGET)^14^, and Pacific Pediatric Neuro-Oncology Consortium (PNOC)^15^, and one broad multi-tissue study: the Genotype Tissue Expression Project (GTEx)^16^. The data included almost every RNA-seq sample from the datasets and occupied a total of 108 Terabytes. This pipeline uses STAR to generate read mappings^17^ and read coverage graphs, and performs isoform quantification using both RSEM^18^ and Kallisto^19^ (**Fig. 1, Supplementary Note 3**). Processing the samples on up to 32,000 AWS cores took 90 hours of wall time, 368,000 jobs and 1, 325,936 core hours. The cost per sample was $1.30 — a more than thirtyfold reduction in cost, and a similar reduction in time, over a comparable TCGA best-practices workflow^5^, while achieving a 98% gene-level concordance with the previous pipeline’s expression predictions (**Fig. 1, Fig. 2, Supplementary Fig. 1**). Notably, we estimate the pipeline, without STAR and RSEM, can be used to generate quantifications for $0.19/sample with Kallisto. To illustrate portability, the same pipeline was run on the I-SPY2 dataset (156 samples) using a private HPC cluster, achieving similar per sample performance (Supplementary Table 1). 20 Expression level signal graphs (read coverage) of the GTEx data (7304 samples from 53 tissues, 570 donors) are available from a UCSC Genome Browser^21^ public track hub (Supplementary Fig. 2). Gene and isoform quantifications for this consistent, union dataset are publicly accessible through UCSC Xena^20^ and available for direct access through S3 (**Supplementary Fig. 3, Supplementary Note 5**).

To achieve these time and cost efficiencies, Toil includes numerous performance optimizations (**Supplementary Note 4**). Toil implements a leader/worker pattern for job scheduling, where the leader makes major decisions regarding the jobs delegated to workers. To reduce pressure on the leader, workers intelligently decide whether they are capable of running jobs immediately downstream to their assigned task (in terms of resource requirements and workflow dependencies). Frequently, next-generation sequencing workflows are I/O bound due to the large volume of data analysed. To mitigate this, Toil implements file caching and streaming. Where possible, successive jobs that share files are scheduled upon a single node, and caching prevents the need for repeated transfers from the file store transparently. Toil is robust to job failure — workflows can be resumed after any combination of leader and worker failures. Such robustness permits workflows to use low-cost preemptable machines, that can be terminated by the provider at short notice and are currently available at a significant discount on AWS and Google. We estimate the use of preemptable machines on AWS lowered the cost of our RNA-seq compute 2.5 fold, despite encountering over 2,000 premature terminations (Fig. 2). Toil also supports fine-grained resource requirements, enabling each job to specify its core, memory, and local storage needs for scheduling efficiency.

## Summary

A key motivation for developing Toil was to provide a free and open-source solution for large-scale biomedical computation. While there is extensive history of open-source workflow execution software^22–24^, the shift to cloud platforms and the advent of standard workflow languages is changing the landscape. Having portable and open workflow software that supports open community standards for workflow specification frees groups to move their computation according to cost, time, and existing data location. We argue such flexibility is not only efficient, but transformative, in that it allows the envisioning of larger, more comprehensive analyses, and for other groups to quickly reproduce results using precisely the original computations. To illustrate that point, had we used the original TCGA best-practices RNA-seq pipeline to analyse the constructed dataset, it would have cost about $800,000, but through algorithmic efficiencies and Toil we reduced that cost to $26,071. We anticipate the union expression dataset we have computed will be widely used — it being the first to quantify consistently across both TCGA and GTEx, therefore combining the largest public RNA-seq repositories of somatic and germline samples.

## Acknowledgements

This work was supported by (BD2K) the National Human Genome Research Institute of the National Institutes of Health under Award Number 5U54HG007990 and (Cloud Pilot) the National Cancer Institute of the National Institutes of Health under the Broad Institute Subaward Number 5417071-5500000716. The UCSC Genome Browser work was supported by the NHGRI award 5U41HG002371 (Corporate Sponsors) Research reported in this publication was also supported by gifts from Microsoft, Google, and Amazon. The content is solely the responsibility of the authors and does not necessarily represent the official views of the National Institutes of Health or our corporate sponsors.

## Author Contributions

J.V., A.R., and B.P wrote the manuscript. J.V., A.R., A.N., J.A., C.K., J.N., H.S., P.A., S.F., B.O. and B.P. contributed to Toil development. F.A.N. and A.M. contributed to Toil-Spark integration. J.V. wrote the RNA-seq pipeline and automation software. M.H. and C.B. contributed WDL and cloud support. P.A., S.Z. contributed CWL support. J.Z., B.C. and M.G. hosted quantification results on UCSC Xena. K.R. hosted GTEx results in UCSC Genome Browser. W.J.K, J.Z., S.Z., G.G., D.A.P., A.D.J., D.H., and B.P. provided scientific leadership and project oversight.

## Additional information

### Data Deposition statement

Data is available from this project at the Toil xena hub at https://genome-cancer.soe.ucsc.edu/proj/site/xena/datapages/?host=https://toil.xenahubs.net

### Required

Reprints and permissions information is available at http://www.nature.com/reprints

## Supplementary information for *Rapid and efficient analysis of20,000 RNA-seq samples with Toil*

For convenience, a version of Toil’s official documentation is included at the end of the supplement and is linked to from parts of the supplementary information.

An up-to-date version of Toil’s documentation can be found here: http://toil.readthedocs.org/

The Toil source code is freely viewable at: https://github.com/BD2KGenomics/toil

A repository of Toil pipelines is available at: https://github.com/BD2KGenomics/toil-scripts

### Supplementary Note 1 - Toil Architecture and Frontend API

(Appendix - Chapter 6: Developing a Workflow) Starting from ‘Hello World’, this section guides the user through all the key pieces of information needed to start developing Toil scripts and pipelines.

(Appendix - Chapter 7: Toil API) Explanation of Toil’s front-end syntax for creating and running jobs, storing and retrieving intermediate files, and useful features like subgraph encapsulation and promises.

(Appendix - Chapter 8: Toil Architecture) A high-level overview of Toil’s architecture, design choices, and optimizations.

### Supplementary Note 2 - Toil Implementation APIs

(Appendix - Chapter 9: The Batch System Interface) Describes the methods and functions that comprise the abstractBatchSystem class. The batch system manages the scheduling and supervision of jobs.

(Appendix - Chapter 10: Job Store Interface) Describes the methods and functions that comprise the absractJobStore class. The job store is where intermediate files are stored during runtime along with information pertaining to job completion which allows for resumability.

### Supplementary Note 3 - RNA-seq Analysis

#### Description

We describe the RNA-seq pipeline used in this recompute, referred to as the CGL RNA-seq pipeline. The pipeline uses STAR^1^ as its primary aligner which uses suffix arrays to achieve its mapping speed at the cost of requiring 40 GB of memory in which to load the array. RSEM^2^ was selected as an expression quantifier as it was used in the TCGA best practices pipeline and results were directly comparable between the two pipelines which allowed for validation. The other quantification tool in the pipeline is Kallisto^3^, a rapid transcript-level quantification tool that can analyze thirty million unaligned paired-end RNA-Seq reads in under 5 minutes. Kallisto is fast for two main reasons: it doesn’t require alignment to a reference genome, and it exploits the fact that it is more important to know which read came from which transcript, and not *where* within the transcript it is located. STAR, RSEM, and Kallisto all require input files to be generated before they can be used for their primary function. All input directories or files were generated from the HG38 reference genome^4^ and Gencode’s v23 comprehensive CHR annotation file^5^. The reference genome did not contain alternative sequences and had the overlapping genes from the PAR locus removed (chrY:10,000-2,781,479 and chrY:56,887,902-57,217,415) due to duplicate gene conflicts with the X chromosome.

#### Methods

The CGL RNA-seq pipeline was written in Python using the Toil API. This script was run on a distributed cluster in AWS using the Mesos batch system and CGCloud to provision and grow the cluster. A separate script was written to collect metrics in 30 minute increments and terminate instances that had grown idle.

We use Amazon’s server-side encryption, with customer-provided encryption keys (SSE-C), to encrypt all of our sample data securely in S3. We have an in-house system of generating per-file SSE-C keys that are derived from a master encryption key that has been left out of the following method so that it is reproducible.

1. Create head node in EC2

a. Launch EC2 ubuntu 14.04 instance (we used an m4.xlarge instance type)
b. `git clone` https://github.com/BD2KGenomics/cgcloud
c. `cd cgcloud`
d. `git checkout f630984b56a68ccdce51e5943a4aba9f7a18470b`
e. `virtualenv venv`
f. `source venv/bin/activate`
g. `pip install boto tqdm`
h. `make develop sdist`
i. Add the following to the bash profile (~/.profile):

i. `export CGCLOUD_ZONE=us-west-2a`
ii. `export CGCLOUD_PLUGINS=cgcloud.mesos:cgcloud.toil`
j. Generate a private key with ssh-keygen if necessary.
k. `cgcloud register-key ~/.ssh/id_rsa`
l. Create Amazon Machine Image with necessary prerequisites:

i. `cgcloud create-IT toil-box`
m. Create a credentials file in ~/.boto by adding the following:

i. `[Credentials]`
ii. `aws_access_key_id = ACCESS_KEY`
iii. `aws_secret_access_key = SECRET_KEY`
n. Create a folder, mkdir shared_dir, that contains:

i. https://github.com/BD2KGenomics/toil-scripts/blob/toil-recompute/src/toil_scripts/rnaseq_cgl/rnaseq_cgl_pipeline.py
ii. config.txt that contains one sample UUID,URL per line
2. Launch initial cluster. We used a bid price of $1.00, and us-west-2a or us-west-2b for the zone.

a. `eval $(ssh-agent); ssh-add`
b. cgcloud create-cluster-leader-instance-type params.leader_type-instance-type c3.8xlarge-share shared_dir-num-workers 1-cluster-name rnaseq-recompute-spot-bid <bid price>-leader-on-demand-zone <availability zone>-ssh-opts "-o UserKnownHostsFile=/dev/null-o StrictHostKeyChecking=no" toil
3. Launch pipeline by creating a screen session on the master, then passing the command to launch the pipeline script in the screen session.

a. `cgcloud ssh-cluster-name rnaseq-recompute toil-leader-o—UserKnownHostsFile=/dev/null-o StrictHostKeyChecking=no screen-dmS rnaseq-recompute`
b. cgcloud ssh-cluster-name params.cluster_name toil-leader-o UserKnownHostsFile=/dev/null-o StrictHostKeyChecking=no screen-S params.cluster_name-X stuff "python /home/mesosbox/shared/rnaseq_cgl_pipeline.py aws:us-west-2:rnaseq-recompute-config /home/mesosbox/shared/config.txt-retryCount 2-s3_dir cgl-rnaseq-recompute-batchSystem=mesos-mesosMaster mesos-master:5050-workDir=/var/lib/toil-wiggle-save_bam >& log.txt\n"
4. Begin metric collection

a. Download metric collection and cluster down-sizing script: https://github.com/jvivian/one_off_scripts/blob/master/toil_uber_script.py
b. `screen-S metrics`
c. `python toil_uber_script.py launch-metrics-c rnaseq-recompute`
d. Exit the screen: `ctrl+a d`
5. Grow cluster. We grew the cluster in increments of 100 nodes so we could monitor our impact on the spot market price and performance of the pipeline.
6. `screen-S grow`
7. `cgcloud grow-cluster-cluster-name rnaseq-recompute-list-num-workers <number>-spot-bid <bid price<-instance-type c3.8xlarge-z <availability zone> toil`

A heuristic to grow the cluster, given some target cluster size, is currently being developed, but wasn’t available in time for the recompute. The cluster was instead grown manually, and automatically downsized as nodes grew idle during periodic metric collection.

#### Input Files

STAR, RSEM, and Kallisto indexes were all derived from the same reference genome and annotation file. Kallisto requires a transcriptome instead of a complete reference genome to build its index file, which was created using BEDtool’s *fastaFromBed.*

- Reference genome - HG38 (no alt analysis)^4^ with overlapping genes from the PAR locus removed (chrY:10,000-2,781,479 and chrY:56,887,902-57,217,415).
- Annotation file - Gencode V23 comprehensive CHR^5^.
- STAR

- sudo docker run-v $(pwd):/data quay.io/ucsc_cgl/star-runThreadN 32-runMode genomeGenerate-genomeDir /data/genomeDir-genomeFastaFiles hg38.fa-sjdbGTFfile gencode.v2 3.annotation.gtf
- Kallisto

sudo docker run-v $(pwd):/data quay.io/ucsc_cgl/kallisto index-i hg38.gencodeV2 3.transcripts.idx transcriptome_hg38_gencodev23.fasta
- RSEM

`sudo docker run-v $(pwd):/data-entrypoint=rsem-prepare-reference jvivian/rsem-p 4-gtf gencode.v2 3.annotation.gtf hg38.fa hg38`

#### Source Code

A current implementation of the TCGA RNA-seq pipeline in Toil and the CGL RNA-seq pipeline can be found here: https://github.com/BD2KGenomics/toil-scripts/tree/master/src/toil_scripts

Script used to launch cluster, launch pipeline, collect metrics, and terminate idle instances: https://github.com/jvivian/one_off_scripts/blob/master/toil_uber_script.py

### Supplementary Note 4 Toil Optimizations

(Appendix - Chapter 8, Section 1: Toil Optimizations) Toil implements several optimizations designed for scalability. This section details some of the most important optimizations.

### Supplementary Note 5 - Data Availability

The RNA-seq recompute produced both quantification and wiggle track format data. The GTEx^6^ wiggle tracks were used by the UC Santa Cruz Browser^7^ team to assemble comprehensive tracks as seen in Supplementary Figure 2. To access the GTEx data on the UCSC Browser, go to the UCSC Genome Browser Track Hubs Page, click the **My Hubs** tab, and paste the following URL: http://hgdownload.soe.ucsc.edu/hubs/gtex/hub.txt into the the **URL** box and hit **Add Hub**. The quantification data is being hosted on UCSC Xena^8^, a platform that can be used to download the raw tables as well as explore the data through a variety of different visualizations as seen in Supplementary Figure 3.

Accession of the raw RNA-seq recompute data is available from S3. Our group has enabled **requester-pays** on the bucket to avoid the significant costs associated with data egress out of AWS. Users can transfer data to EC2 for free.

1. Create an account with AWS
2. Install s3cmd: `pip install s3cmd`
3. Type `s3cmd ––configure`, and follow the prompts to setup s3cmd.
4. Type `s3cmd ls s3://cgl-rnaseq-recompute-fixed` to list the bucket contents
5. Type `s3cmd get ––requester-pays [S3 URL]` to get any file in the bucket.

Download example for a random TCGA sample:

`s3cmd get––requester-pays \ s3://cgl-rnaseq-recompute-fixed/tcga/014ee344-2844-4bc4-842e-e6d6e1e8fd9b.tar.gz`

Due to the large size of the bucket, and number of files, the command s3cmd sync usually hangs. To download all of the data associated with a particular project, download any of the manifests located in the root directory and iterate over the s3 URLs to download all of the samples: for i in $(cat tcga-manifest); do s3cmd get ––requester-pays $i; done

Manifests are available using the following commands:

- `s3cmd get ––requester-payss3://cgl-rnaseq-recompute-fixed/tcga-manifest`
- `s3cmd get ––requester-payss3://cgl-rnaseq-recompute-fixed/gtex-manifest`
- `s3cmd get ––requester-payss3://cgl-rnaseq-recompute-fixed/target-manifest`
- `s3cmd get ––requester-payss3://cgl-rnaseq-recompute-fixed/pnoc-manifest`

### Supplementary Note 6 - Spark Support

A feature unique to Toil is support for long-lived services that can run alongside jobs in a workflow. This feature is necessary for supporting tools that run on distributed systems such as Apache Spark^11^ or Hadoop, legacy high-performance computing systems like the Message Passing Interface (MPI), or a high performance SQL database. There has been significant recent interest in using Apache Spark for processing genomic data, with significant projects including ADAM^12,13^, Hadoop-BAM^14^, and the upcoming release of the Genome Analysis Toolkit (GATK 4.0).

To use one of these processing frameworks, the framework is usually configured and started on a cluster of nodes. In a typical workflow management system, jobs are only connected by producer/consumer relationships: the output of a job is consumed by another job. However, persistent services, such as an Apache Spark cluster, cannot be covered easily by a strict producer/consumer relationship. Specifically, the cluster must be alive whenever a job that is using it is alive, and jobs do not use the output of the cluster, rather, they dynamically interact with it. This is implemented in Toil via the Service Job interface Appendix - Chapter 6, Section 11: Services). A Service Job persists as long as the children jobs that can access it are alive. A single service can spawn subservices, as is necessary for supporting complex services like Apache Spark.

Apache Spark is built around a leader-worker architecture, where an application driver submits tasks to the Spark leader, which then schedules the tasks on the workers in the cluster. We have implemented code in the `toil.lib.spark` module that spawns a Spark cluster using the Service Job interface. This works by first spawning a Spark cluster leader service job. Once this job has started up, we spawn the requisite number of Spark worker nodes. The IP address of the Spark leader node is passed to the Spark worker service jobs, and to the child jobs that will access the cluster using Toil’s promise functionality. The child jobs run the Spark application drivers, and attach to the Spark cluster using the IP address of the leader. Currently, we support Apache Spark 1.5.2 running on top of Apache Hadoop Distributed File System (HDFS) 2.6.2, which is deployed using Docker containers. The presence of this functionality enables the rapid setup of a Spark cluster on any batch system that Toil can run on.

To validate our Spark cluster implementation, we performed *k*-mer counting on the HG00096 sample from the 1000 Genomes. We used a naäve *k*-mer counter implemented using ADAM^12,13^. The results of our run can be found in Supplementary Figure 4. We used a Toil cluster running on AWS using the `r3.8xlarge` instance type. This instance type has 32 cores, 244 Gibibytes (GiB) of memory, and two 320 GiB SSDs. We scaled the number of Spark workers in our workflow from one to eight in four logarithmic steps. As can be seen in Supplementary Figure 4, the runtime of the *k*-mer counter improves by a factor of two with each step.

### Supplementary Note 7 - CWL Support

The Common Workflow Langauge (CWL) is a joint effort to establish a standardized specification for writing workflows used by the scientific community^15^. Tools are described using CWL’s Doman Specific Language (DSL), which details a tool’s inputs, outputs, and commands. These tools are wired together as *steps* inside a workflow, also written in CWL. Installing Toil with the CWL extra provides an entrypoint to **cwltoil** that allows execution of CWL workflows and tools using Toil’s infrastructure.

To run a CWL workflow using the Toil engine, assuming /**data** is a location with plenty of storage:

- Install `s3cmd` if not installed (see Supplementary Note 5)
- `s3cmd get ––requester-pays s3://cgl-rnaseq-recompute-fixed/cwl-example.tar.gz`
- `tar-xvf cwl-example.tar.gz && cd cwl-example`
- `virtualenv ~/cwl && source ~/cwl/bin/activate`
- `pip install toil[cwl,aws]`
- `cwltoil wf-alignment.cwl alignment-only.json—workDir=/data ––jobStore=/data/cwl-jobStore`

To run a sample on an AWS cluster running the Mesos batchsystem, run the following command on the leader:

`cwltoil wf-alignment.cwl alignment-only.json—batchSystem=mesos \ ––mesosMaster=mesos-master:5050 ––jobStore=aws:us-west-2:cwltoil-test`

### Supplementary Figure 1 Metrics

**Supplementary Figure 1.**
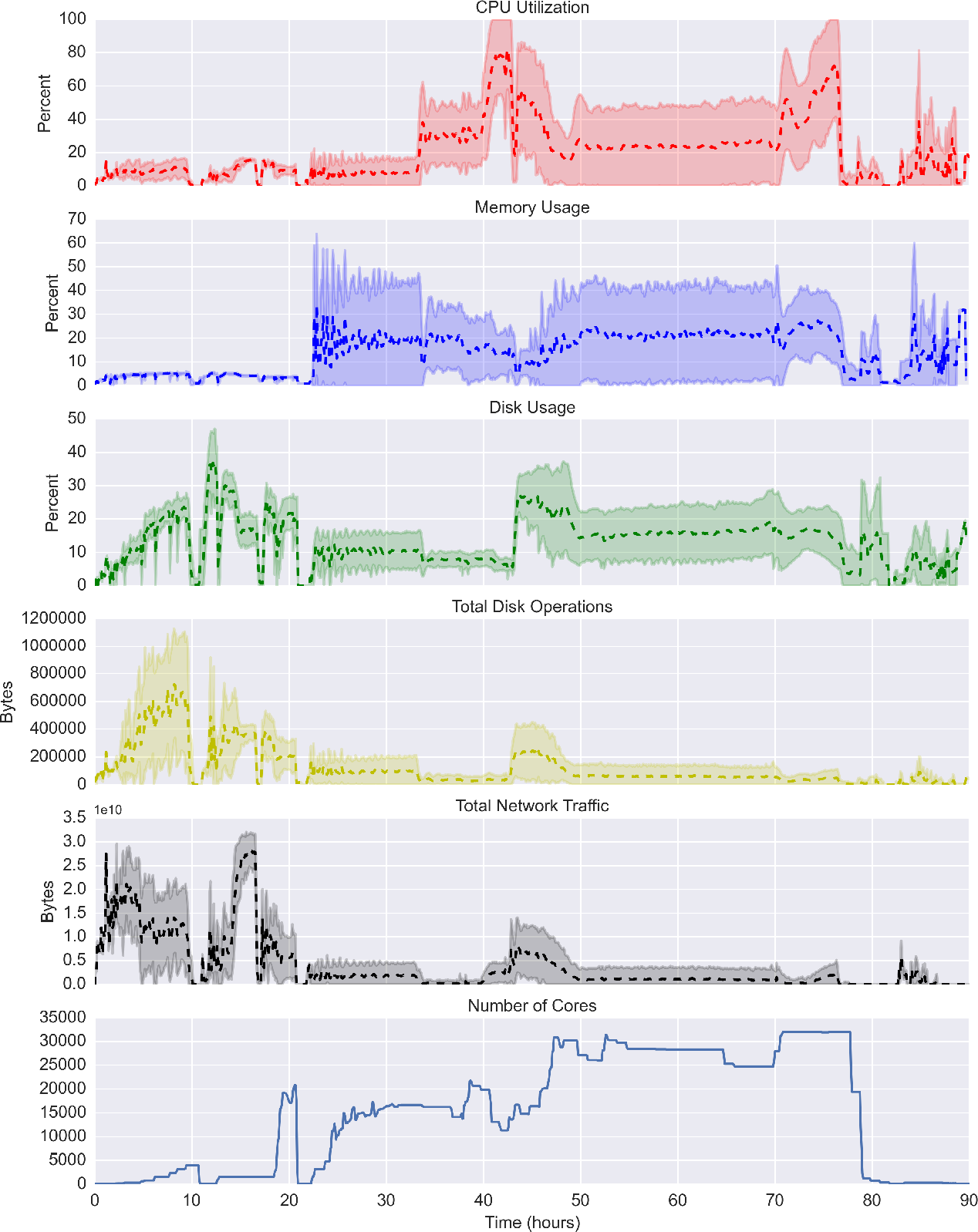
Aggregate metrics collected during the Toil RNA-seq recompute. Metrics were collected every 5 minutes and averaged across every instance running at that time point. Complete termination of the Mesos framework occurred around hours 10 and 20, from which the pipeline was resumed.

### Supplementary Figure 2 Browser

**Supplementary Figure 2.**
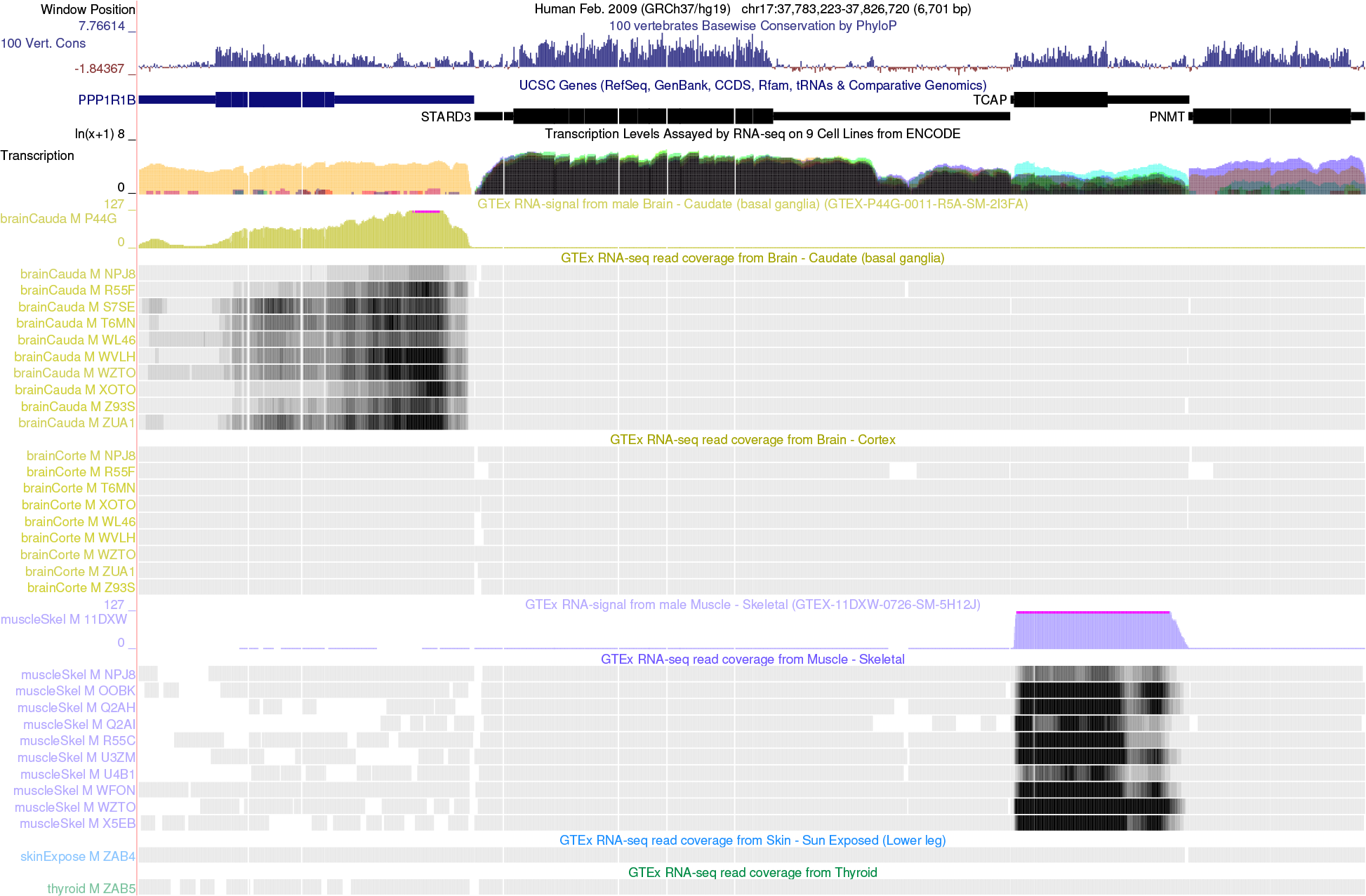
6700 bp exon-focused view of a 43 Kbp region of human chromosome 17 where GTEx RNA-seq highlights tissue-specific expression of two genes. The TCAP (titin cap protein) is highly expressed in muscle tissue, while PP1R1B (a therapeutic target for neurologic disorders) shows expression in brain basal ganglia but not muscle (or brain cortex). In this UCSC Genome Browser view, 33 samples from 5 tissues were selected for display, from the total 7304 (in 53 tissues) available on the GTEx public track hub. The hub is available on both hg19 (GRCh37) and hg38 (GRCh38) human genome assemblies. The hg19 tracks were generated using the UCSC liftOver tool to transform coordinates from the hg38 bedGraph files generated by STAR2 in the Toil pipeline. The browser display was configured to use the Multi-region exon view.

### Supplementary Figure 3 Xena

**Supplementary Figure 3.**
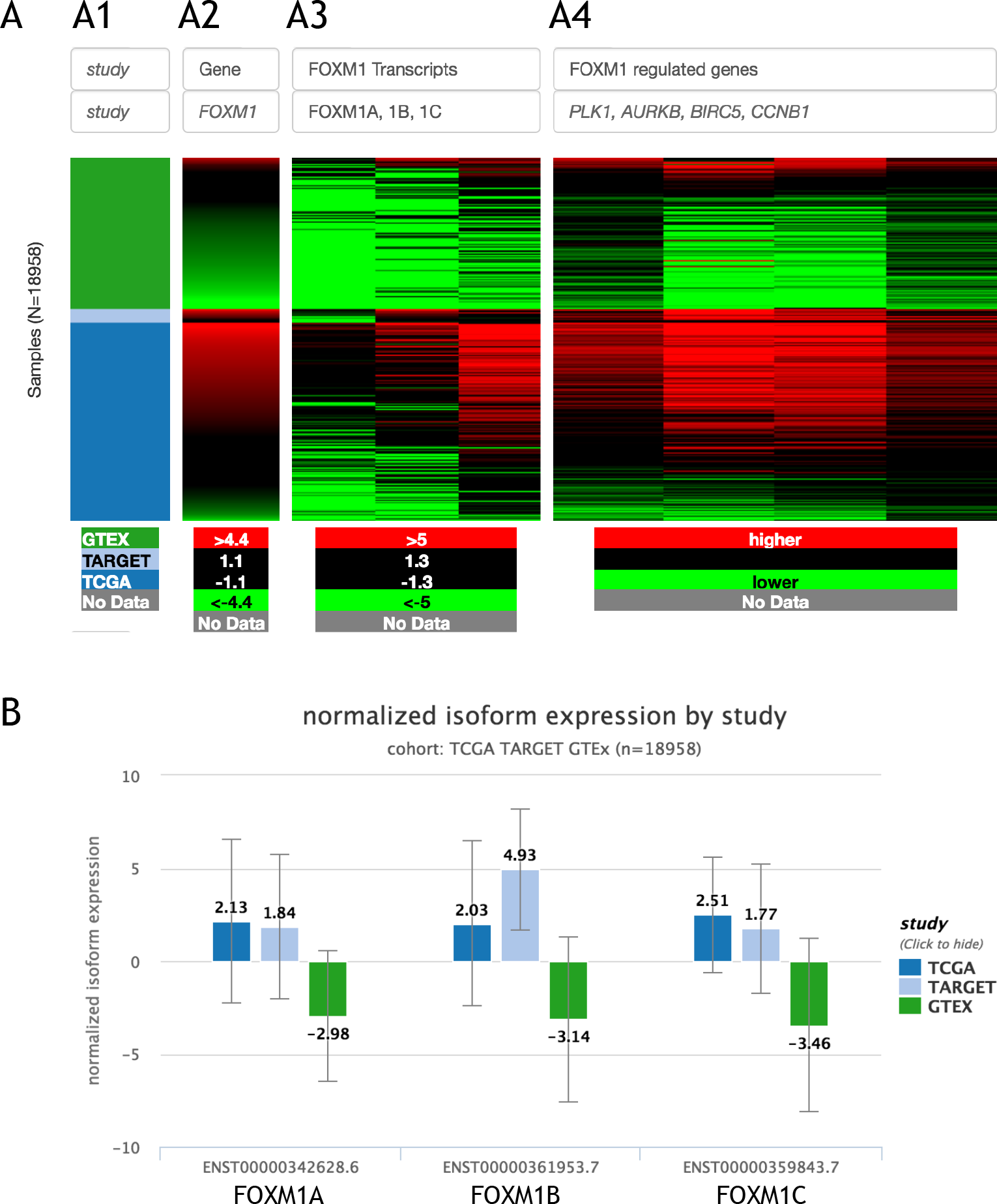
FOXM1 is a member of the FOX family of transcription factors. FOXM1 plays a key role in cell cycle progression. FOXM1 regulates expression of a large array of G2/M-specific genes, such as PLK1, CCNB2, BIRC5, and AURKB^9^ There are three FOXM1 isoforms, A, B and C. FOXM1A is known to a transcriptional repressor, while FOXM1B and 1C are known to be transcritional activators. (**A**) We exam FOXM1 gene expression (**A2**), isoform A, B, C expression (**A3**), and expression of four cell cycle genes (A4) in the TCGA, TARGET, GTEx toil RNA-seq re-compute results, using UCSC Xena^8^. A cohort of 19,948 samples drawn from all three studies shows the FOXM1B and FOXM1C are significantly up-regulated in tumors comparing to GTEx normal samples. Toil recompute expression estimation are hosted on Toil hub^10^ (**B**) Summary view of FOXM1 A, B, and C isoform expression in the TCGA, TARGET and GTEx samples.

### Supplementary Figure 4 Spark Support

**Supplementary Figure 4.**
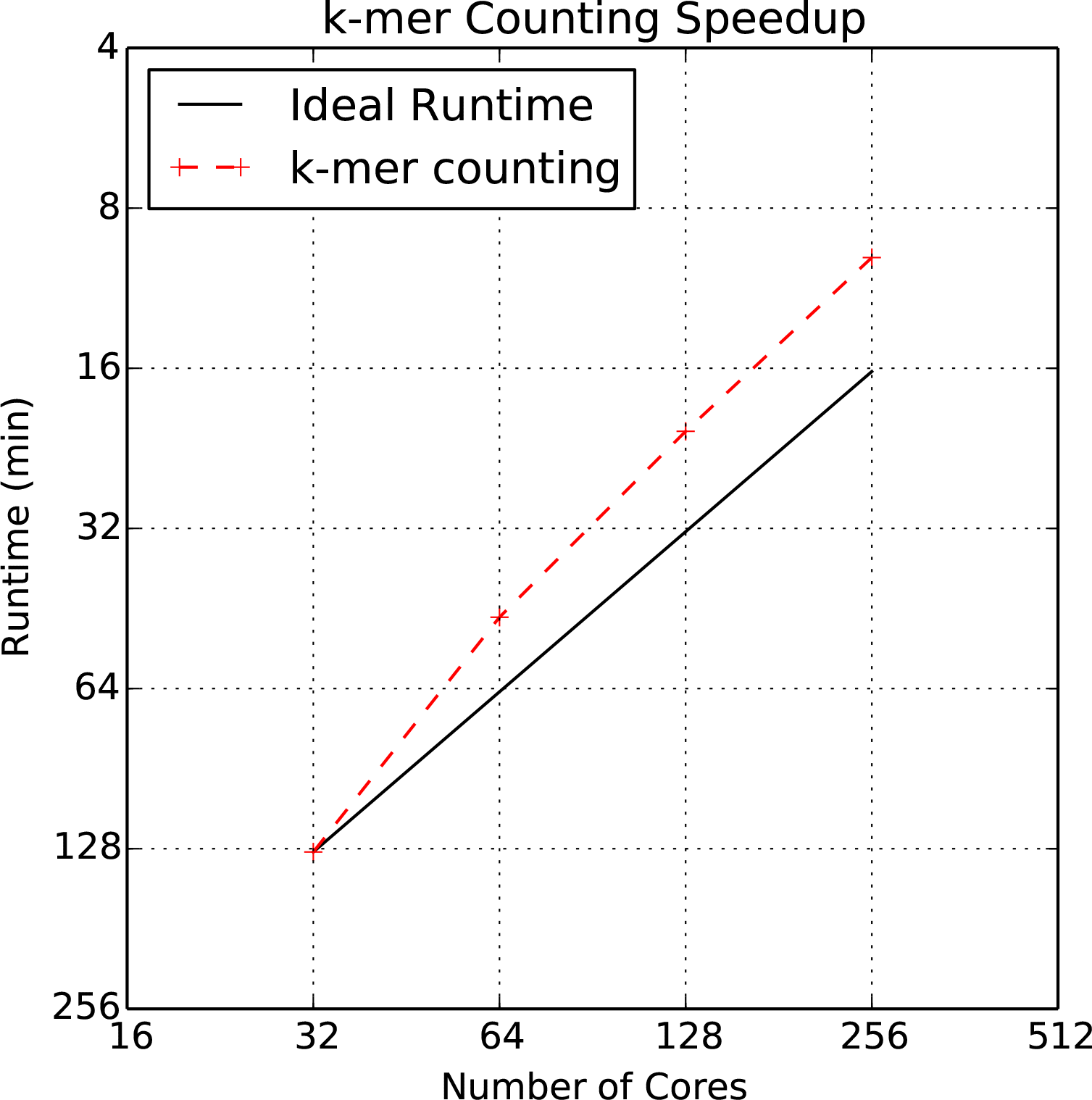
Spark runtimes and *k*-mer count distribution. We plotted the runtime of the *k*-mer counting experiment against the number of cores being used for counting. Note that the time axis is flipped: the experiments with more cores took less time to complete.

### Supplementary Table 1 OpenStack I-SPY Analysis

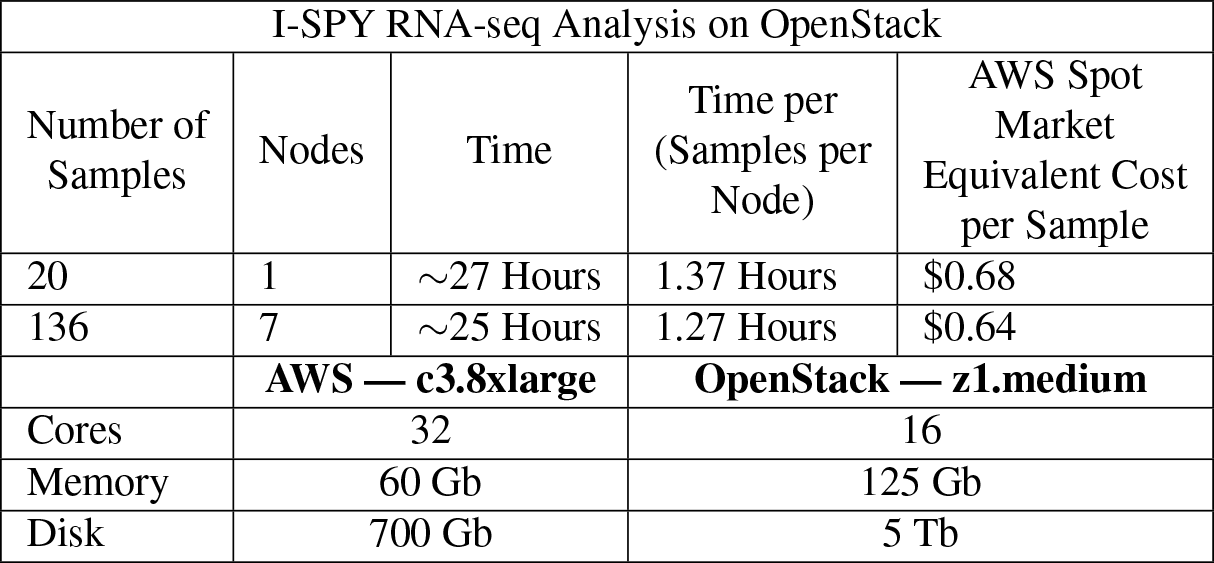
To demonstrate Toil on a non-AWS system, 156 samples were processed from the I-SPY breast cancer dataset on the UC Santa Cruz OpenStack cluster. Samples and inputs were stored in the associated Ceph object store for locality and cost-free transfer to OpenStack. These samples were run using Toil’s singleMachine batch system, which allowed us to use the local job store implementation, meaning files could be locally copied instead of being uploaded to a remote job store (like S3). Samples processed faster due to a few factors: smaller sample size, local job store is faster than AWS job store, and the large disk space meant no bottlenecks for jobs with large disk requirements.

| I-SPY RNA-seq Analysis on OpenStack |  |  |  |  |
| --- | --- | --- | --- | --- |
| Number of Samples | Nodes | Time | Time per (Samples per Node) | AWS Spot Market Equivalent Cost per Sample |
| 20 | 1 | ~27 Hours | 1.37 Hours | \$0.68 |
| 136 | 7 | ~25 Hours | 1.27 Hours | \$0.64 |
|  | **AWS — c3.8xlarge** |  | **OpenStack — z1.medium** |  |
| Cores | 32 |  | 16 |  |
| Memory | 60 Gb |  | 125 Gb |  |
| Disk | 700 Gb |  | 5 Tb |  |

## UCSC Computational Genomics Lab

Toil is a workflow engine entirely written in Python. It features:

- Easy installation, e.g. `pip install toil`.
- Common Workflow Language (CWL) support Complete support for the draft-3 CWL specification, allowing it to execute CWL workflows.
- Cross platform support Develop and test on your laptop then deploy on any of the following:

- Commercial clouds: - Amazon Web Services (including the spot market) - Microsoft Azure - Google Compute Engine
- Private clouds: - OpenStack
- High Performance Computing Environments: - GridEngine - Apache Mesos - Parasol - Individual multi-core machines
- A small API Easily mastered, the Python user API for defining and running workflows is built upon one core class.
- Complete file and stream management: Temporary and persistent file management that abstracts the details of the underlying file system, providing a uniform interface regardless of environment. Supports both atomic file transfer and streaming interfaces, and provides encryption of user data.
- Scalability: Toil can easily handle workflows concurrently using hundreds of nodes and thousands of cores.
- Robustness: Toil workflows support arbitrary worker and leader failure, with strong check-pointing that always allows resumption.
- Efficiency: Caching, fine grained, per task, resource requirement specifications, and support for the AWS spot market mean workflows can be executed with little waste.
- Declarative and dynamic workflow creation: Workflows can be declared statically, but new jobs can be added dynamically during execution within any existing job, allowing arbitrarily complex workflow graphs with millions of jobs within them.
- Support for databases and services: For example, Apache Spark clusters can be created quickly and easily integrated within a toil workflow as a service, with precisely defined time start and end times that fits with the flow of other jobs in the workflow.
- Open Source: An Apache license allows unrestricted use, incorporation and modification.

Contents:

## INSTALLATION

### 1.1 Prerequisites

- Python 2.7.x
- pip > 7.x

### 1.2 Basic installation

To setup a basic Toil installation use

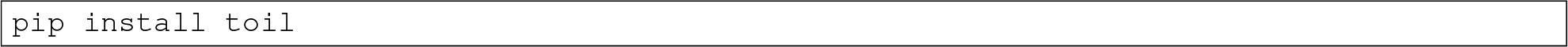

Toil uses setuptools’ extras mechanism for dependencies of optional features like support for Mesos or AWS. To install Toil with all bells and whistles use

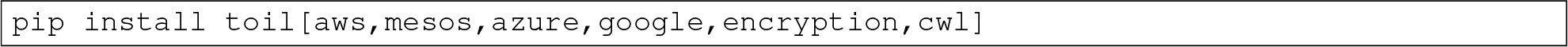

Here’s what each extra provides:

- The `aws` extra provides support for storing workflow state in Amazon AWS. This extra has no native dependencies.
- The `google` extra is experimental and stores workflow state in Google Cloud Storage. This extra has no native dependencies.
- The `azure` extra stores workflow state in Microsoft Azure Storage. This extra has no native dependencies.
- The `mesos` extra provides support for running Toil on an Apache Mesos cluster. Note that running Toil on SGE (GridEngine), Parasol or a single machine does not require an extra. The `mesos` extra requires the following native dependencies:

–*Apache Mesos*
–*Python headers and static libraries*
- The `encryption` extra provides client-side encryption for files stored in the Azure and AWS job stores. This extra requires the following native dependencies:

–*Python headers and static libraries*
–*Libffi headers and library*
- The `cwl` extra provides support for running workflows written using the Common Workflow Language.

##### Apache Mesos

Only needed for the `mesos` extra. Toil has been tested with version 0.25.0. Mesos can be installed on Linux by following the instructions on https://open.mesosphere.com/getting-started/install/. The Homebrew package manager has a formula for Mesos such that running `brew install mesos` is probably the easiest way to install Mesos on OS X. This assumes, of course, that you already have Xcode and Homebrew.

Please note that even though Toil depends on the Python bindings for Mesos, it does not explicitly declare that dependency and they will **not** be installed automatically when you run `pip install toil[mesos]`. You need to install the bindings manually. The Homebrew formula for OS X installs them by default. On Ubuntu you will need to download the appropriate.egg from https://open.mesosphere.com/downloads/mesos/ and install it using `easy_install-a <path_to_egg>`. Note that on Ubuntu Trusty you may need to upgrade `protobuf` via `pip install ––upgrade protobuf` **before** running the above `easy_install` command.

If you intend to install Toil with the mesos extra into a virtualenv, be sure to create that virtualenv with

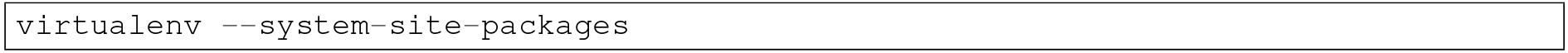

Otherwise, Toil will not be able to import the `mesos.native` module.

##### Python headers and static libraries

Only needed for the `mesos` and `encryption` extras. The Python headers and static libraries can be installed on Ubuntu/Debian by running `sudo apt-get install build-essential python-dev` and accordingly on other Linux distributions. On Mac OS X, these headers and libraries are installed when you install the Xcode command line tools by running `xcode-select ––install`, assuming, again, that you have Xcode installed.

##### Libffi headers and library

Libffiis only needed for the `encryption` extra. To install Libffi on Ubuntu, run `sudo apt-get install libffi-dev`. On Mac OS X, run `brew install libffi`. This assumes, of course, that you have Xcode and Homebrew installed.

### 1.3 Building & testing

For developers and people interested in building the project from source the following explains how to setup virtualenv to create an environment to use Toil in.

After cloning the source and cd-ing into the project root, create a virtualenv and activate it:

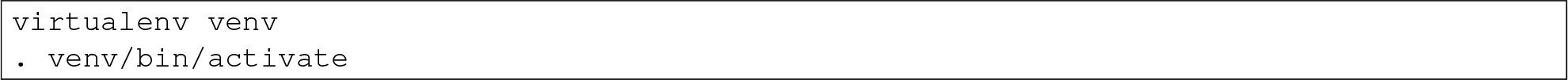

Simply running from the project root will print a description of the available Makefile targets.

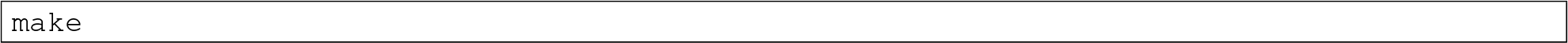

Once you created and activated the virtualenv, the first step is to install the build requirements. These are additional packages that Toil needs to be tested and built, but not run:

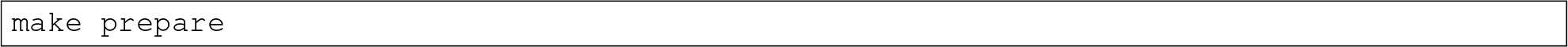

Once the virtualenv has been prepared with the build requirements, running

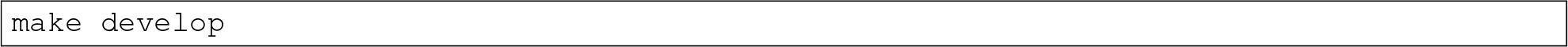

will create an editable installation of Toil and its runtime requirements in the current virtualenv. The installation is called *editable* (also known as a development mode installation) because changes to the Toil source code immediately affect the virtualenv. Optionally, set the extras variable to ensure that make develop installs support for optional extras. Consult setup.py for the list of supported extras. To install Toil in development mode with all extras run

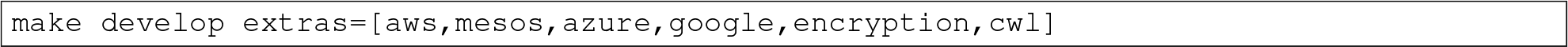

Note that some extras have native dependencies as listed in *Basic installation.* Be sure to install them before running the above command. If you get

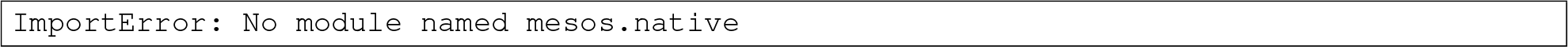

make sure you install Mesos and the Mesos egg as described in *Apache Mesos* and be sure to create the virtualenv with ––system-site-packages.

To build the docs, run `make develop` with all extras followed by

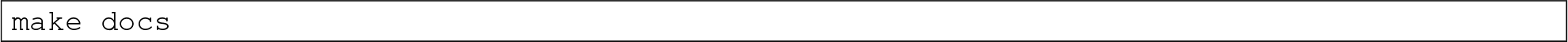

To invoke the tests (unit and integration) use

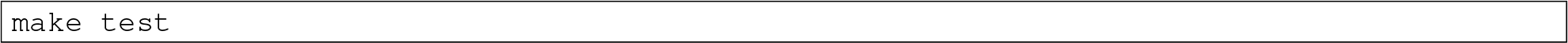

Run an individual test with

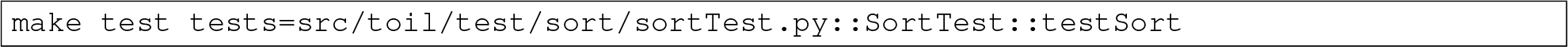

The default value for `tests` is "`src`" which includes all tests in the src subdirectory of the project root. Tests that require a particular feature will be skipped implicitly. If you want to explicitly skip tests that depend on a currently installed *feature*, use

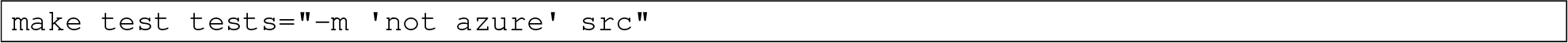

This will run only the tests that don’t depend on the `azure` extra, even if that extra is currently installed. Note the distinction between the terms *feature* and *extra*. Every extra is a feature but there are features that are not extras, the `gridengine` and `parasol` features fall into that category. So in order to skip tests involving both the Parasol feature and the Azure extra, the following can be used:

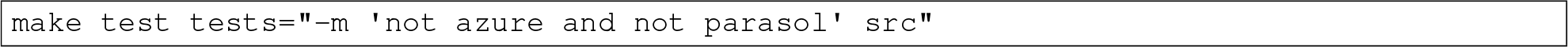

#### 1.3.1 Running Mesos tests

See *Apache Mesos*. Be sure to create the virtualenv with `––system-site-packages` to include the Mesos Python bindings. Verify by activating the virtualenv and running ‥ `pip list | grep mesos`. On OS X, this may come up empty. To fix it, run the following:

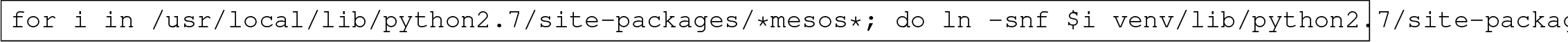

## CLOUD INSTALLATION

### 2.1 Installation on AWS for distributed computing

We use CGCloud to provision instances and clusters in AWS. Thorough documentation of CGCloud can be found in the CGCloud-core and CGCloud-toil documentation. Brief steps will be provided to those interested in using CGCloud for provisioning.

#### 2.1.1 CGCloud in a nutshell

Setting up clusters with CGCloud has the benefit of coming pre-packaged with Toil and Mesos, our preferred batch system for running on AWS. If you encounter any issues following these steps, check official documentation which contains Troubleshooting sections.

1. `virtualenv ~/cgcloud`
2. `source ~/cgcloud/bin/activate`
3. `pip install cgcloud-core`
4. `pip install cgcloud-toil`
5. **Add the following to your ~/.profile, use the appropriate region for your account.** 5a. `export CGCLOUD_ZONE=us-west-2a` 5b. `export CGCLOUD_PLUGINS=`"`cgcloud.toil:$CGCLOUD_PLUGINS`"
6. Setup credentials for your AWS account in `~/.aws/credentials`:

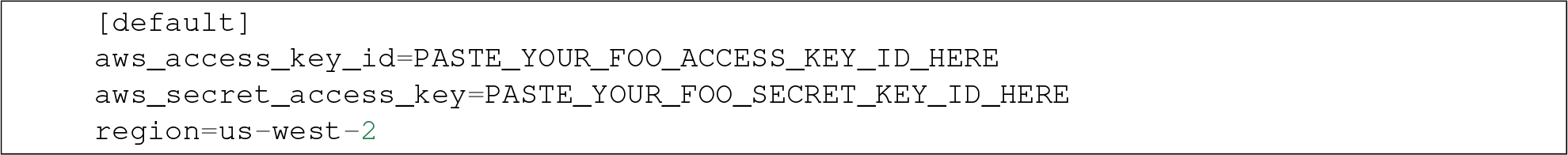
7. **Register your SSH key. You can create one with ssh-keygen.** 7a. `cgcloud register-key ~/.ssh/id_rsa.pub`
8. **Create a template *toil-box* which will contain necessary prerequisites** 8a. `cgcloud create-IT toil-box`
9. **Create a small leader/worker cluster** 9a. `cgcloud create-cluster toil-s 2-t m3.large`
10. SSH into the leader: `cgcloud ssh toil-leader`

At this point, any toil script can be run on the distributed AWS cluster following instructions in *Running on AWS*.

### 2.2 Installation on Azure

While CGCloud does not currently support cloud providers other than Amazon, Toil comes with a cluster template to facilitate easy deployment of clusters running Toil on Microsoft Azure. The template allows these clusters to be created and managed through the Azure portal.

Detailed information about the template is available here.

To use the template to set up a Toil Mesos cluster on Azure, follow these steps.

1. Make sure you have an SSH RSA public key, usually stored in `~/.ssh/id_rsa.pub`. If not, you can use `ssh-keygen-t rsa` to create one.
2. Click on the deploy button above, or navigate to https://portal.azure.com/#create/Microsoft.Template/uri in your browser.
3. If necessary, sign into the Microsoft account that you use for Azure.
4. You should be presented with a screen resembling the following:

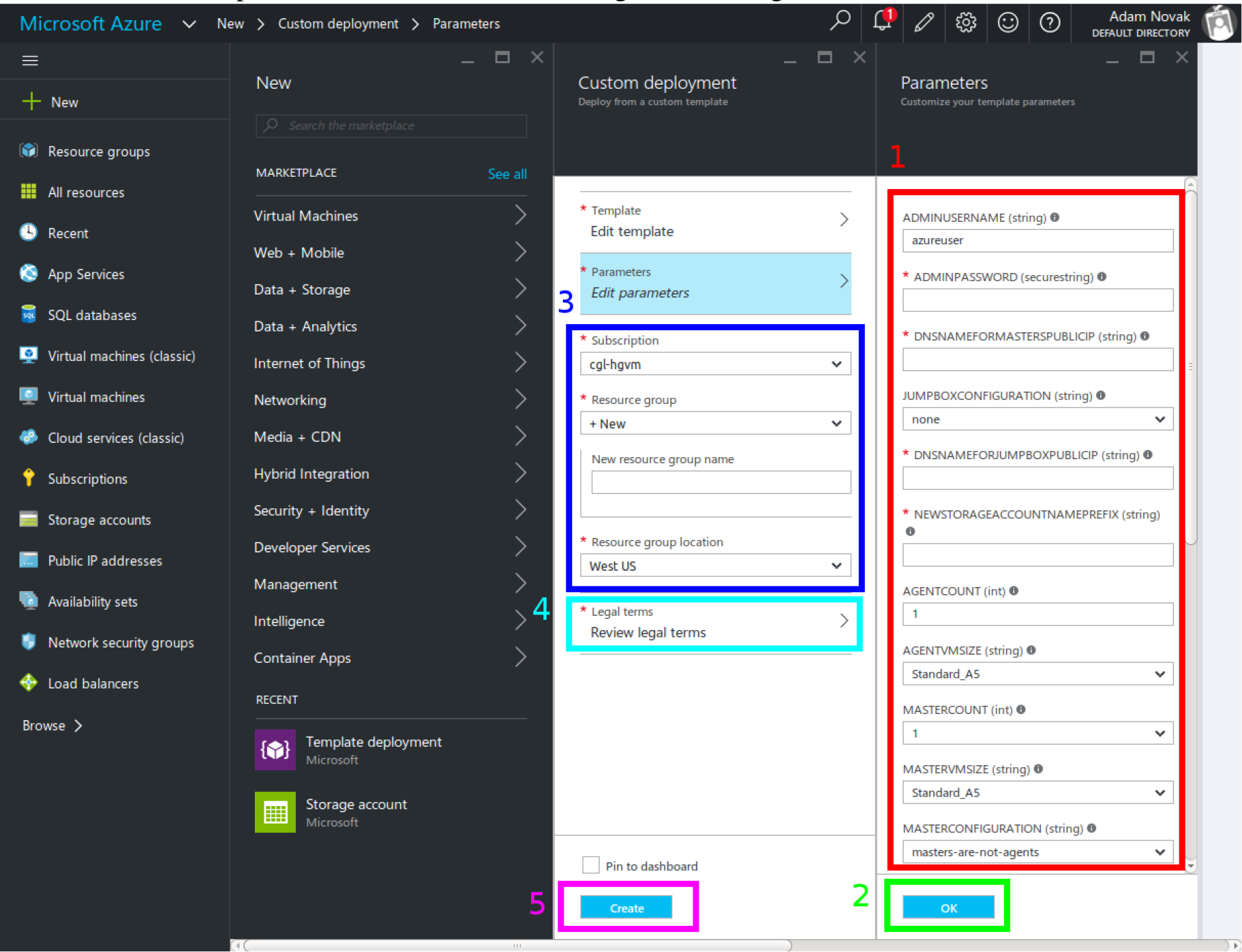
5. Fill out the form on the far right (marked “1” in the image), giving the following information. Important fields for which you will want to override the defaults are in bold:

a. **AdminUsername**: Enter a username for logging into the cluster. It is easiest to set this to match your username on your local machine.
b. **AdminPassword**: Choose a strong root password. Since you will be configuring SSH keys, you will not actually need to use this password to log in in practice, so choose something long and complex and store it safely.
c. **DnsNameForMastersPublicIp**: Enter a unique DNS name fragment to identify your cluster within your region. For example, if you are putting your cluster in `westus`, and you choose `awesomecluster`, your cluster’s public IP would be assigned the name `awesomecluster.westus.cloudapp.azure.com`.
d. JumpboxConfiguration: If you would like, you can select to have either a Linux or Windows “jumpbox” with remote desktop software set up on the cluster’s internal network. By default this is turned off, since it is unnecessary.
e. DnsNameForJumpboxPublicIp: If you are using a jumpbox, enter another unique DNS name fragment here to set its DNS name. See `DnsNameForMastersPublicIp` above.
f. **NewStorageAccountNamePrefix**: Enter a globally unique prefix to be used in the names of new storage accounts created to support the cluster. Storage account names must be 3 to 24 characters long, include only numbers and lower-case letters, and be globally unique. Since the template internally appends to this prefix, it must be shorter than the full 24 characters. Up to 20 should work.
g. **AgentCount**: Choose how many agents (i.e. worker nodes) you want in the cluster. Be mindful of your Azure subscription limits on both VMs (20 per region by default) and total cores (also 20 per region by default); if you ask for more agents or more total cores than you are allowed, you will not get them all, errors will occur during template instantiation, and the resulting cluster will be smaller than you wanted it to be.
h. **AgentVmSize**: Choose from the available VM instance sizes to determine how big each node will be. Again, be mindful of your Azure subscription’s core limits. Also be mindful of how many cores and how much disk and memory your Toil jobs will need: if any requirement is greater than that provided by an entire node, a job may never be scheduled to run.
i. MasterCount: Choose the number of “masters” or leader nodes for the cluster. By default only one is used, because although the underlying Mesos batch system supports master failover, currently Toil does not. You can increase this if multiple Toil jobs will be running and you want them to run from different leader nodes. Remember that the leader nodes also count against your VM and core limits.
j. MasterVmSize: Select one of the available VM sizes to use for the leader nodes. Generally the leader node can be relatively small.
k. MasterConfiguration: This is set to `masters-are-not-agents` by default, meaning that the leader nodes will not themselves run any jobs. If you are worried about wasting unused computing power on your leader nodes, you can set this to `masters-are-agents` to allow them to run jobs. However, this may slow them down for interactive use, making it harder to monitor and control your Toil workflows.
l. JumpboxVmSize: If you are using a jumpbox, you can select a VM instance size for it to use here. Again, remember that it counts against your Azure subscription limits.
m. ClusterPrefix: This prefix gets used to generate the internal hostnames of all the machines in the cluster. You can use it to give clusters friendly names to differentiate them. It has to be a valid part of a DNS name; you might consider setting it to match `DnsNameForMastersPublicIp`. You can also leave it at the default.
n. SwarmEnabled: You can set this to `true` to install Swarm, a system for scheduling Docker containers. Toil does not use Swarm, and Swarm has a tendency to allocate all the cluster’s resources for itself, so you should probably leave this set to `false` unless you also find yourself needing a Swarm cluster.
o. MarathonEnabled: You can set this to `true` to install Marathon, a scheduling system for persistent jobs run in Docker containers. It also has nothing to do with Toil, and should probably remains et to `false`.
p. ChronosEnabled: You can set this to `true` to install Chronos, which is a way to periodically run jobs on the cluster. Unless you find yourself needing this functionality, leave this set to `false`. (All these extra frameworks are here because the Toil Azure template was derived from a Microsoft template for a generic Mesos cluster, offering these services.)
q. ToilEnabled: You should leave this set to `true`. If you set it to `false`, Toil will not be installed on the cluster, which rather defeats the point.
r. **SshRsaPublicKey**: Replace `default` with your SSH public key contents, beginning with `ssh-rsa`. Paste in the whole line. Only one key is supported, and as the name suggests it must be an RSA key. This enables SSH key-based login on the cluster.
s. GithubSource: If you would like to install Toil from a nonstandard fork on Github (for example, installing a version inclusing your own patches), set this to the Github fork (formatted as `<username>`/`<reponame<`) from which Toil should be downloaded and installed. If not, leave it set to the default of `BD2KGenomics/toil`.
t. **GithubBranch**: To install Toil from a branch other than master, enter the name of its branch here. For example, for the latest release of Toil 3.1, enter `releases/3.1.x`. By default, you will get the latest and greatest Toil, but it may have bugs or breaking changes introduced since the last release.
6. Click OK (marked “2” in the screenshot).
7. Choose a subscription and select or create a Resource Group (marked “3” in the screenshot). If creating a Resource Group, select a region in which to place it. It is recommended to create a new Resource Group for every cluster; the template creates a large number of Azure entitites besides just the VMs (like virtual networks), and if they are organized into their own Resource Group they can all be cleaned up at once when you are done with the cluster, by deleting the Resource Group.
8. Read the Azure terms of service (by clicking on the item marked “4” in the screenshot) and accept them by clicking the “Create” button on the right (not shown). This is the contract that you are accepting with Microsoft, under which you are purchasing the cluster.
9. Click the main “Create” button (marked “5” in the screenshot). This will kick off the process of creating the cluster.
10. Eventually you will receive a notification (Bell icon on the top bar of the Azure UI) letting you know that your cluster has been created. At this point, you should be able to connect to it; however, note that it will not be ready to run any Toil jobs until it is finished setting itself up.
11. SSH into the first (and by default only) leader node. For this, you need to know the `AdminUsername` and `DnsNameForMastersPublicIp` you set above, and the name of the region you placed your cluster in. If you named your user phoebe and named your cluster `toilisgreat`, and placed it in the `centralus`— region, the hostname of the cluster would be `toilisgreat`.`centralus`.`cloudapp`.`azure.com`, and you would want to connect as `phoebe`. SSH is forwarded through the cluster’s load balancer to the first leader node on port 2211, so you would run ssh phoebe@toilisgreat.centralus.cloudapp.azure.com-p 2211.
12. Wait for the leader node to finish setting itself up. Run `tail-f/var/log/azure/cluster-bootstrap.log` and wait until the log reaches the line completed mesos cluster configuration. At that point, kill `tail` with a `ctrl-c`. Your leader node is now ready.
13. At this point, you can start running Toil jobs, using the Mesos batch system (by passing `––batchSystem mesos ––mesosMaster 10.0.0.5:5050`) and the Azure job store (for which you will need a separate Azure Storage account set up, ideally in the same region as your cluster but in a different Resource Group). The nodes of the cluster may take a few more minutes to finish installing, but when they do they will report in to Mesos and begin running any scheduled jobs.
14. Whan you are done running your jobs, go back to the Azure portal, find the Resource Group you created for your cluster, and delete it. This will destroy all the VMs and any data stored on them, and stop Microsoft charging you money for keeping the cluster around. As long as you used a separate Asure Storage account in a different Resource Group, any information kept in the job stores and file stores you were using will be retained.

For more information about how your new cluster is organized, for information on how to access the Mesos Web UI, or for troubleshooting advice, please see the template documentation.

### 2.3 Installation on OpenStack

Our group is working to expand distributed cluster support to OpenStack by providing convenient Docker containers to launch Mesos from. Currently, OpenStack nodes can be setup to run Toil in **singleMachine** mode following the basic installation instructions: *Basic installation*

### 2.4 Installation on Google Compute Engine

Support for running on Google Cloud is experimental, and our group is working to expand distributed cluster support to Google Compute by writing a cluster provisioning tool based around a Dockerized Mesos setup. Currently, Google Compute Engine nodes can be configured to run Toil in **singleMachine** mode following the basic installation instructions: *Basic installation*

## RUNNING A WORKFLOW

### 3.1 Running quick start

Starting with Python, a Toil workflow can be run with just three steps.

1. `pip install toil`
2. Copy and paste the following code block into HelloWorld.py:

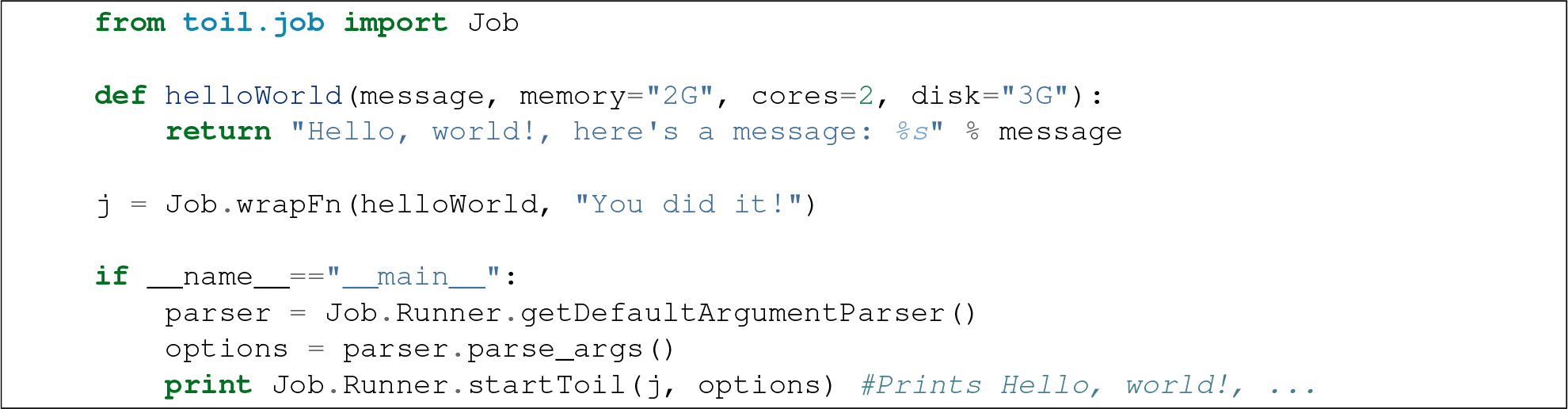
3. `python HelloWorld.py file:jobStore`

Now you have run Toil on **singleMachine** (default batch system) using the **FileStore** job store. The first positional argument after the. `py` is the location of the job store, a place where intermediate files are written to. In this example, a folder called **jobStore** will be created where **HelloWorld.py** is run from. Information on the jobStore can be found at *The job store interface.*

Run `python HelloWorld.py ––help` to see a complete list of available options.

For something beyond a hello world example, refer to *Running a Toil pipeline in detail*

### 3.2 Running CWL workflows

The Common Workflow Language (CWL) is an emerging standard for writing workflows that are portable across multiple workflow engines and platforms. To run workflows written using CWL, first ensure that Toil is installed with the “cwl” extra as described in *Basic installation.* This will install the executables `cwl-runner` and `cwltoil` (these are identical, where `cwl-runner` is the portable name for the default system CWL runner). To learn more about CWL, see the CWL User Guide.

To run in local batch mode, simply provide the CWL file and the input object file:

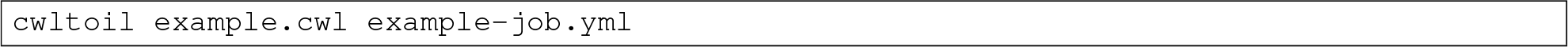

To run in cloud and HPC configurations, you may need to provide additional command line parameters to select and configure the batch system to use. Consult the appropriate sections.

### 3.3 Running a Toil pipeline in detail

For a detailed example and explanation, we’ll walk through running a pipeline that performs merge-sort on a temporary file.

1. Copy and paste the following code into **toil-sort-example.py**:

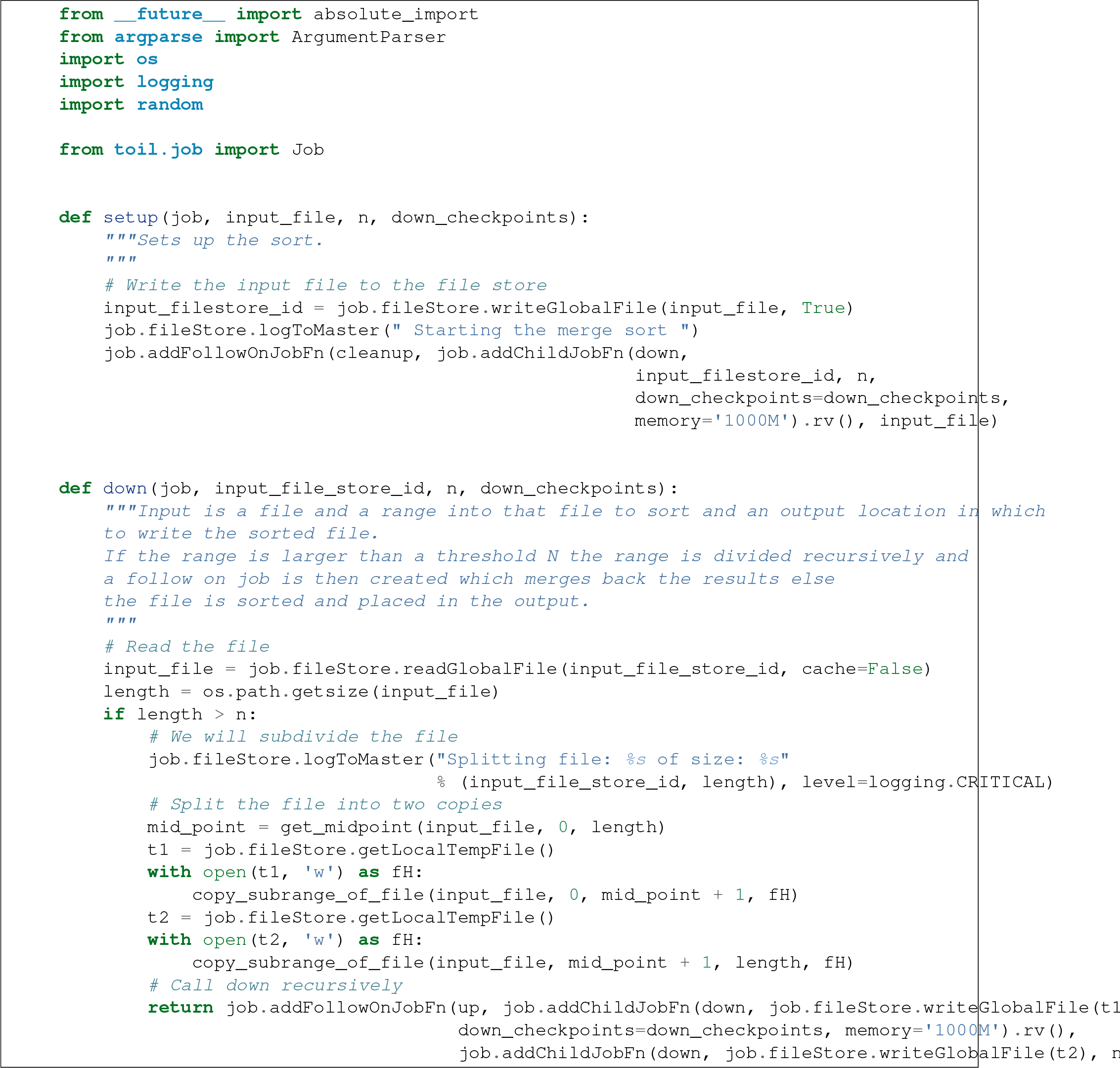

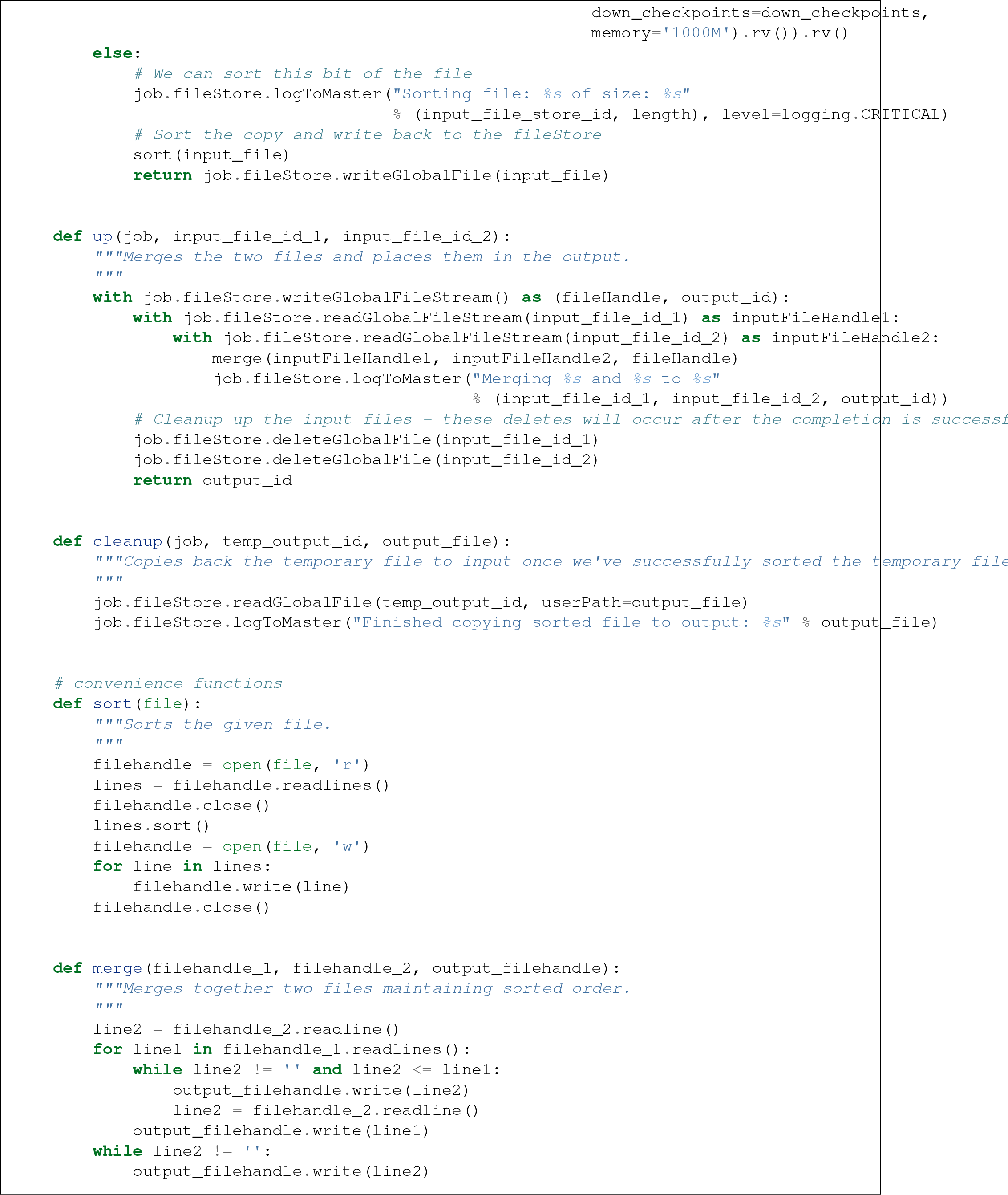

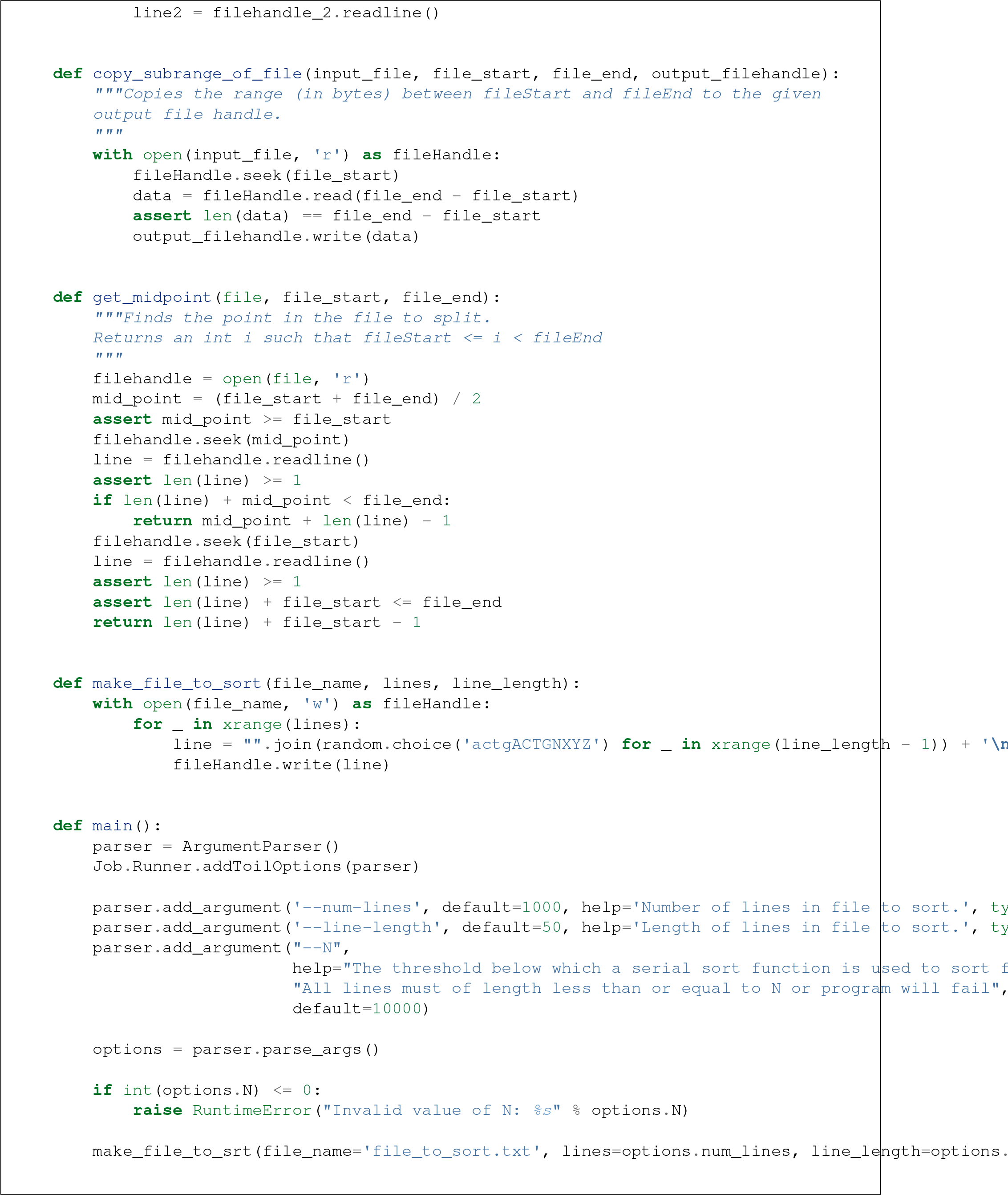

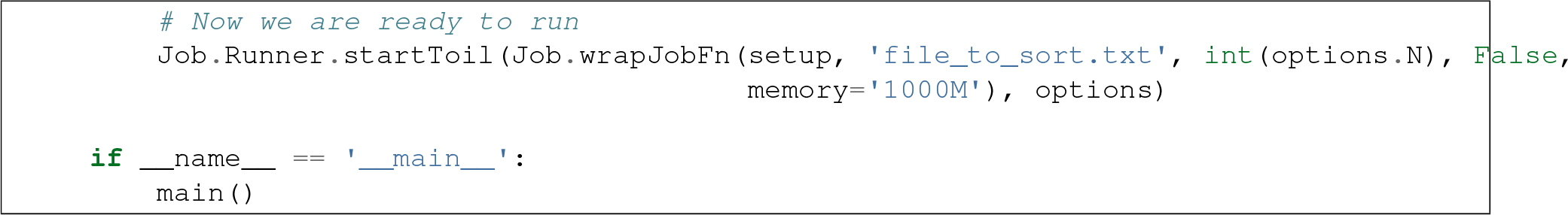
2. Run with default settings: `python toil-sort-example.py file:jobStore`.
3. Run with options: `python toil-sort-example.py file:jobStore ––num-lines 5000 ––line-length 10 ––workDir /tmp/`

The if __name__ == ′__main__ ′ boilerplate is required to enable Toil to import the job functions defined in the script into the context of a Toil *worker* process. By invoking the script you created the *leader process*. A worker process is a separate process whose sole purpose is to host the execution of one or more jobs defined in that script. When using the single-machine batch system (the default), the worker processes will be running on the same machine as the leader process. With full-fledged batch systems like Mesos the worker processes will typically be started on separate machines. The boilerplate ensures that the pipeline is only started once-on the leader-but not when its job functions are imported and executed on the individual workers.

Typing `python toil-sort-example.py ––help` will show the complete list of arguments for the workflow which includes both Toil’s and ones defined inside **toil-sort-example.py**. A complete explanation of Toil’s arguments can be found in *Command line interface and arguments*.

#### 3.3.1 Changing the log statements

When we run the pipeline, we see some logs printed to the screen. At the top there’s some information provided to the user about the environment Toil is being setup in, and then as the pipeline runs we get INFO level messages from the batch system that tell us when jobs are being executed. We also see both INFO and CRITICAL level messages that are in the user script. By changing the logLevel, we can change what we see output to screen. For only CRITICAL level messages: `python toil-sort-examply.py file:jobStore ––logLevel=critical`. This hides most of the information we get from the Toil run. For more detail, we can run the pipeline with `––logLevel=debug` to see a comprehensive output. For more information see *Logging*.

#### 3.3.2 Restarting after introducing a bug

Let’s now introduce a bug in the code, so we can understand what a failure looks like in Toil, and how we would go about resuming the pipeline. On line 30, the first line of the **down()** function, let’s add the line `assert 1==2`, ′`Test Error!`′. Now when we run the pipeline, `python toil-sort-example.py file:jobStore`, we’ll see a failure log under the header **– – – TOIL WORKER OUTPUT LOG– – –**, that contains the stack trace. We see a detailed message telling us that on line 30, in the down fuction, we encountered an error.

If we try and run the pipeline again, we get an error message telling us that a jobStore of the same name already exists. The default behavior for the job store is that it is not cleaned up in the event of failure so that you can restart it from the last succesful job. We can restart the pipeline by running `python toil-sort-example.py file:jobStore ––restart`. We can also change the number of times Toil will attempt to retry a failed job, `python toil-sort-example.py ––retryCount 2 ––restart`. You’ll now see Toil attempt to rerun the failed job, decrementing a counter until that job has exhausted the retry count. `––retryCount` is useful for non-systemic errors, like downloading a file that may experience a sporadic interruption, or some other non-deterministic failure.

To succesfully restart our pipeline, we can edit our script to comment out line 30, or remove it, and then run `python toil-sort-example.py ––restart`. The pipeline will successfully complete, and the job store will be removed.

#### 3.3.3 Getting stats from our pipeline run

We can execute the pipeline to let use retrieve statistics with `python toil-sort-example.py ––stats`. Our pipeline will finish successfully, but leave behind the job store. Now we can type `toil stats file:jobStore` and get back information about total runtime and stats pertaining to each job function.

We can then cleanup our jobStore by running `toil clean file:jobStore`

## RUNNING IN THE CLOUD

There are several recommended ways to run Toil jobs in the cloud. Of these, running on Amazon Web Services (AWS) is currently the best-supported solution.

On all cloud providers, it is recommended that you run long-running jobs on remote systems under `screen`. Simply type `screen` to open a new `screen‵ session. Later, type ‵‵ctrl-a` and then d to disconnect from it, and run `screen-r` to reconnect to it. Commands running under screen will continue running even when you are disconnected, allowing you to unplug your laptop and take it home without ending your Toil jobs.

### 4.1 Running on AWS

See *Installation on AWS for distributed computing* to get setup for running on AWS.

Having followed the *Running quick start* guide, the user can run their **HelloWorld.py** script on a distributed cluster just by modifiying the run command. Since our cluster is distributed, we’ll use the **AWS Jobstore** which creates a job store in S3 instead of on file system.

Place the HelloWorld.py script on the leader node, and run:

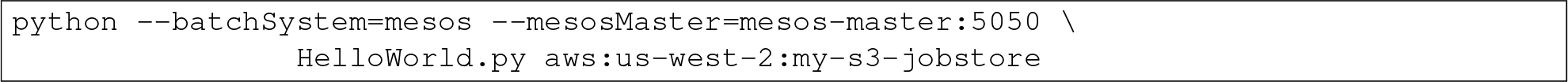

To run a CWL workflow:

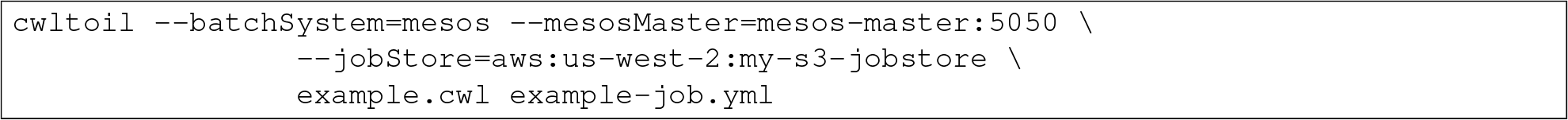

When running a CWL workflow on AWS, input files can be provided either on the local file system or in S3 buckets using s3:// URL references. Final output files will be copied to the local file system of the leader node.

### 4.2 Running on Azure

See *Installation on Azure* to get setup for running on Azure. This section assumes that you are SSHed into your cluster’s leader node.

The Azure templates do not create a shared filesystem; you need to use the **Azure Jobstore**, which needs an Azure Storage Account in which to store its job data. (Note that you can store multiple job stores in a single Azure Storage Account.)

To create a new Storage Account, if you do not already have one:

1. Click here, or navigate to https://portal.azure.eom/#create/Microsoft.StorageAccount in your browser.
2. If necessary, log into the Microsoft Account that you use for Azure.
3. Fill out the presented form. The **Name** for the account, notably, must be a 3-to-24-character string of letters and lowercase numbers that is globally unique. For **Deployment model**, choose “Resource manager”. For **Resource group**, choose or create a resource group **different than** the one in which you created your cluster. For **Location**, choose the **same** region that you used for your cluster.
4. Press the “Create” button. Wait for your Storage Account to be created; you should get a notification in the notifications area at the upper right.

Once you have a Storage Account, you need to authorize the cluster to access the Storage Account, by giving it the access key. To do find your Storage Account’s access key:

1. When your Storage Account has been created, open it up and click the “Settings” icon.
2. In the “Settings” panel, select “Access keys”.
3. Select the text in the “Key1” box and copy it to the clipboard, or use the copy-to-clipboard icon.

You then need to share the key with the cluster. To do this temporarily, for the duration of an SSH or screen session:

1. On the leader node, run `export AZURE_ACCOUNT_KEY=`"`<KEY>`", replacing `<KEY>` with the access key you copied from the Azure portal.

To do this permanently:

1. On the leader node, run `nano ~/.toilAzureCredentials`.
2. In the editor that opens, navigate with the arrow keys, and give the file the following contents:

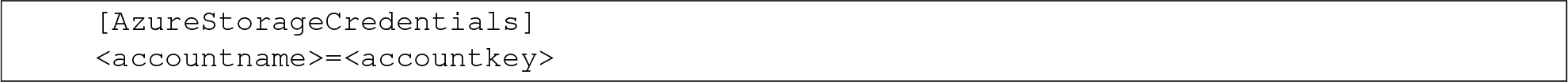 Be sure to replace `<accountname>` with the name that you used for your Azure Storage Account, and `<accountkey>` with the key you obtained above. (If you want, you can have multiple accounts with different keys in this file, by adding multipe lines. If you do this, be sure to leave the `AZURE_ACCOUNT_KEY` environment variable unset.)
3. Press `ctrl-o` to save the file, and `ctrl-x` to exit the editor.

Once that’s done, you are now ready to actually execute a job, storing your job store in that Azure Storage Account. Assuming you followed the *Running quick start* guide above, you have an Azure Storage Account created, and you have placed the Storage Account’s access key on the cluster, you can run the **HelloWorld.py** script by doing the following:

1. Place your script on the leader node, either by downloading it from the command line or typing or copying it into a command-line editor.
2. Run the command:

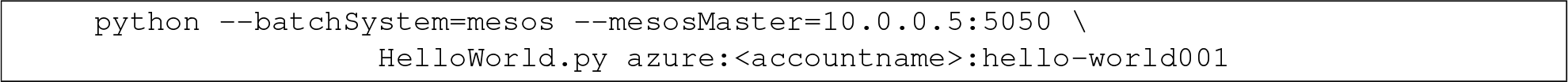 To run a CWL workflow:

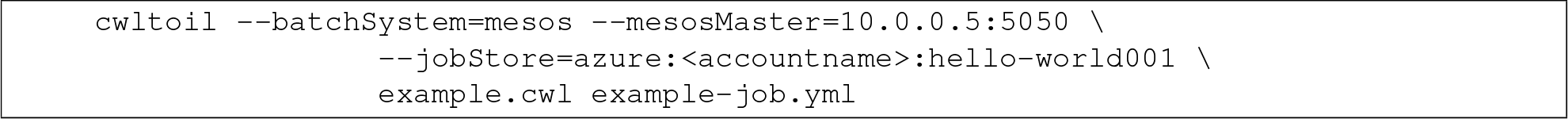 Be sure to replace `<accountname>` with the name of your Azure Storage Account.

Note that once you run a job with a particular job store name (the part after the account name) in a particular Storage Account, you cannot re-use that name in that account unless one of the following happens:

1. You are restarting the same job with the `––restart` option.
2. You clean the job store with `toil clean azure:<accountname>:<jobstore>`.
3. You delete all the items created by that job, and the main job store table used by Toil, from the account (destroying all other job stores using the account).
4. The job finishes successfully and cleans itself up.

### 4.3 Running on Open Stack

After getting setup with *Installation on OpenStack*, Toil scripts can be run just by designating a job store location as shown in *Running quick start*. The location of temporary directories Toil creates to run jobs can be specified with `––workDir:`

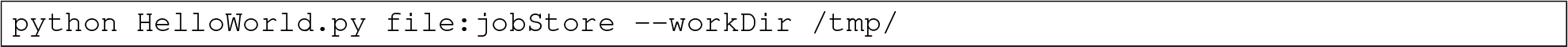

### 4.4 Running on Google Compute Engine

After getting setup with *Installation on Google Compute Engine*, Toil scripts can be run just by designating a job store location as shown in *Running quick start*.

If you wish to use the Google Storage job store, you must install Toil with the ‘google’ extra. Having done this, you must create a file named ‘.boto’ in your home directory with the following format:

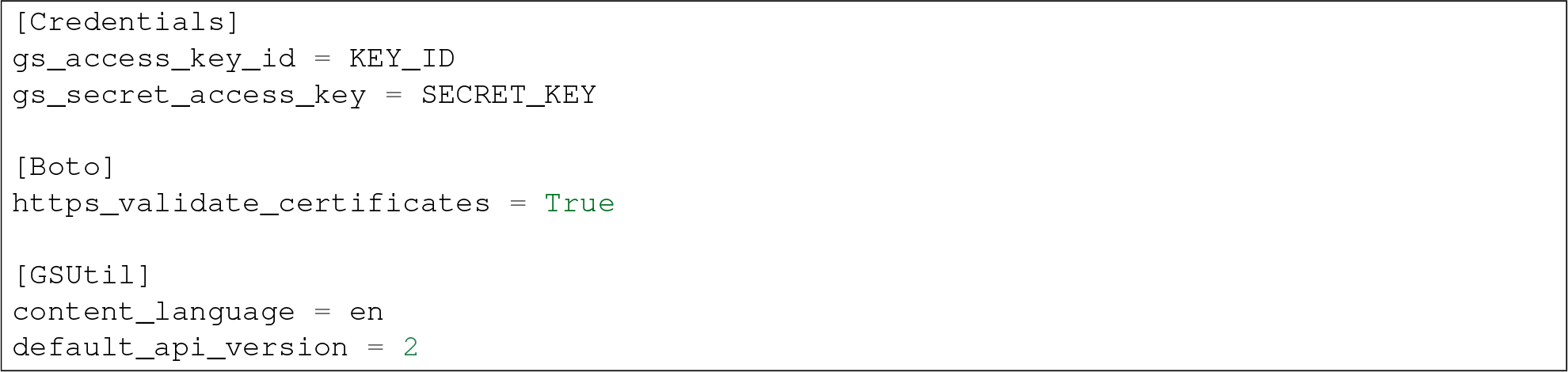

The gs_access_key_id and gs_secret_access_key can be generated by navigating to your Google Cloud Storage console and clicking on ‘Settings’. Then, on the Settings page, navigate to the Interoperability tab and click ‘Enable interoperability access’. On this page you can now click ‘Create a new key’ to generate an access key and a matching secret. Insert these into their respective places in the.boto file and you will be able to use a Google job store when invoking a Toil script, as in the following example:

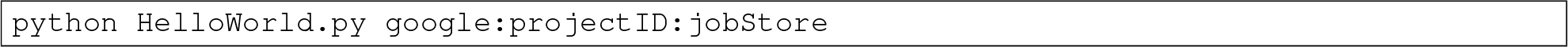

The ‘projectID’ component of the job store argument above refers your Google Cloud project ID in the Google Cloud Console, and will be visible in the console’s banner at the top of the screen. The ‘jobStore’ component is a name of your choosing that you will use to refer to this job store.

## COMMAND LINE INTERFACE AND ARGUMENTS

Toil provides many command line options when running a toil script (see*Running a workflow*), or using Toil to run a CWL script. Many of these are described below. For most Toil scripts executing ‘-help’ will show this list of options.

It is also possible to set and manipulate the options described when invoking a Toil workflow from within Python using `toil.job.Job.Runner.getDefaultOptions()`, e.g.:

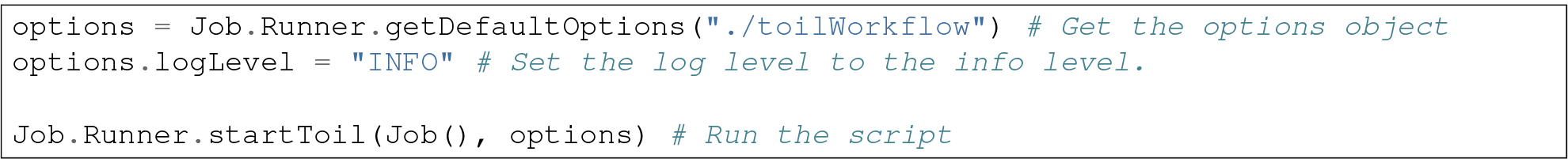

### 5.1 Logging

Toil hides stdout and stderr by default except in case of job failure. For more robust logging options (default is INFO), use `––logDebug` or more generally, use `––logLevel=`, which may be set to either `OFF` (or `CRITICAL`), `ERROR`, `WARN` (or `WARNING`), `INFO` or `DEBUG`. Logs can be directed to a file with `––logFile=`.

If large logfiles are a problem, `––maxLogFileSize` (in bytes) can be set as well as `––rotatingLogging`, which prevents logfiles from getting too large.

### 5.2 Stats

The `––stats` argument records statistics about the Toil workflow in the job store. After a Toil run has finished, the entrypoint `toil stats <jobStore>` can be used to return statistics about cpu, memory, job duration, and more. The job store will never be deleted with `––stats`, as it overrides `––clean`.

### 5.3 Restart

In the event of failure, Toil can resume the pipeline by adding the argument `––restart` and rerunning the python script. Toil pipelines can even be edited and resumed which is useful for development or troubleshooting.

### 5.4 Clean

If a Toil pipeline didn’t finish successfully, or is using a variation of `––clean`, the job store will exist until it is deleted. `toil clean <jobStore>` ensures that all artifacts associated with a job store are removed. This is particularly useful for deleting AWS job stores, which reserves an SDB domain as well as an S3 bucket.

The deletion of the job store can be modified by the `––clean` argument, and may be set to `always`, `onError`, `never`, or `onSuccess` (default).

Temporary directories where jobs are running can also be saved from deletion using the `––cleanWorkDir`, which has the same options as `––clean`. This option should only be run when debugging, as intermediate jobs will fill up disk space.

### 5.5 Batch system

Toil supports several different batch systems using the `––batchSystem` argument. More information in the *The batch system interface*.

### 5.6 Default cores, disk, and memory

Toil uses resource requirements to intelligently schedule jobs. The defaults for cores (1), disk (2G), and memory (2G), can all be changed using `––defaultCores`, `––defaultDisk`, and `––defaultMemory`. Standard suffixes like K, Ki, M, Mi, G or Gi are supported.

### 5.7 Job store

Running toil scripts has one required positional argument: the job store. The default job store is just a path to where the user would like the job store to be created. To use the *Running quick start* example, if you’re on a node that has a large /**scratch** volume, you can specify the jobstore be created there by executing: `python HelloWorld.py /scratch/my-job-store`, or more explicitly, `python HelloWorld.py file:/scratch/my-job-store`. Toil uses the colon as way to explicitly name what type of job store the user would like. Different types of job store options can be looked up in *The job store interface*.

### 5.8 Miscellaneous

Here are some additional useful arguments that don’t fit into another category.

- `––workDir` sets the location where temporary directories are created for running jobs.
- `––retryCount` sets the number of times to retry a job in case of failure. Useful for non-systemic failures like **HTTP** requests.
- `––sseKey` accepts a path to a 32-byte key that is used for server-side encryption when using the AWS job store.
- `––cseKey` accepts a path to a 256-bit key to be used for client-side encryption on Azure job store.
- `––setEnv <NAME=VALUE>` sets an environment variable early on in the worker

## DEVELOPING A WORKFLOW

This tutorial walks through the features of Toil necessary for developing a workflow using the Toil Python API.

### 6.1 Scripting quick start

To begin, consider this short toil script which illustrates defining a workflow:

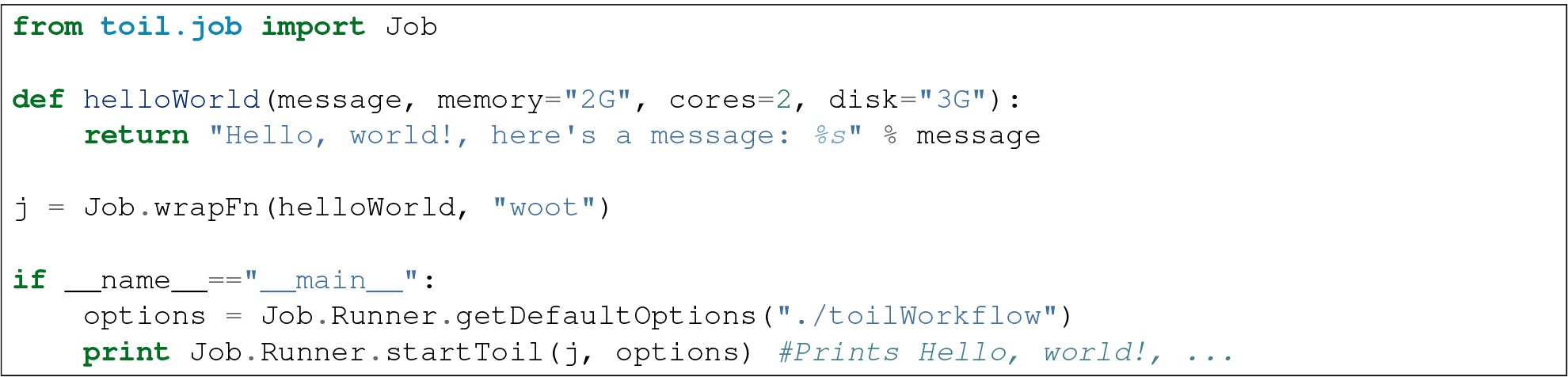

The workflow consists of a single job. The resource requirements for that job are (optionally) specified by keyword arguments (memory, cores, disk). The script is run using `toil.job.Job.Runner.getDefaultOptions()`. Below we explain the components of this code in detail.

### 6.2 Job basics

The atomic unit of work in a Toil workflow is a *job* (`toil.job.Job`). User scripts inherit from this base class to define units of work. For example, here is a more long-winded class-based version of the job in the quick start example:

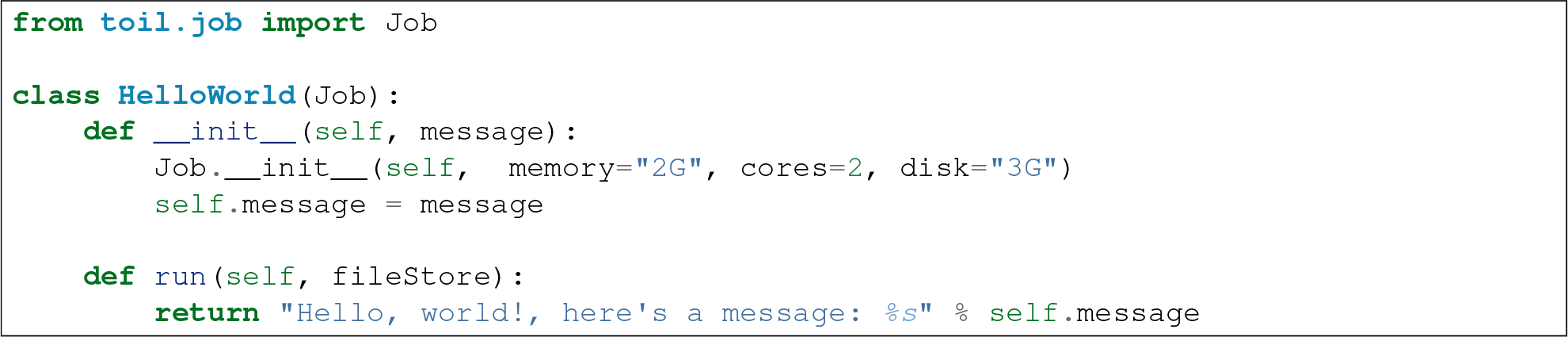

In the example a class, HelloWorld, is defined. The constructor requests 2 gigabytes of memory, 2 cores and 3 gigabytes of local disk to complete the work.

The `toil.job.Job.run()` method is the function the user overrides to get work done. Here it just logs a message using `toil.job.Job.FileStore.logToMaster()`, which will be registered in the log output of the leader process of the workflow.

### 6.3 Invoking a workflow

We can add to the previous example to turn it into a complete workflow by adding the necessary function calls to create an instance of HelloWorld and to run this as a workflow containing a single job. This uses the `toil.job.Job.Runner` class, which is used to start and resume Toil workflows. For example:

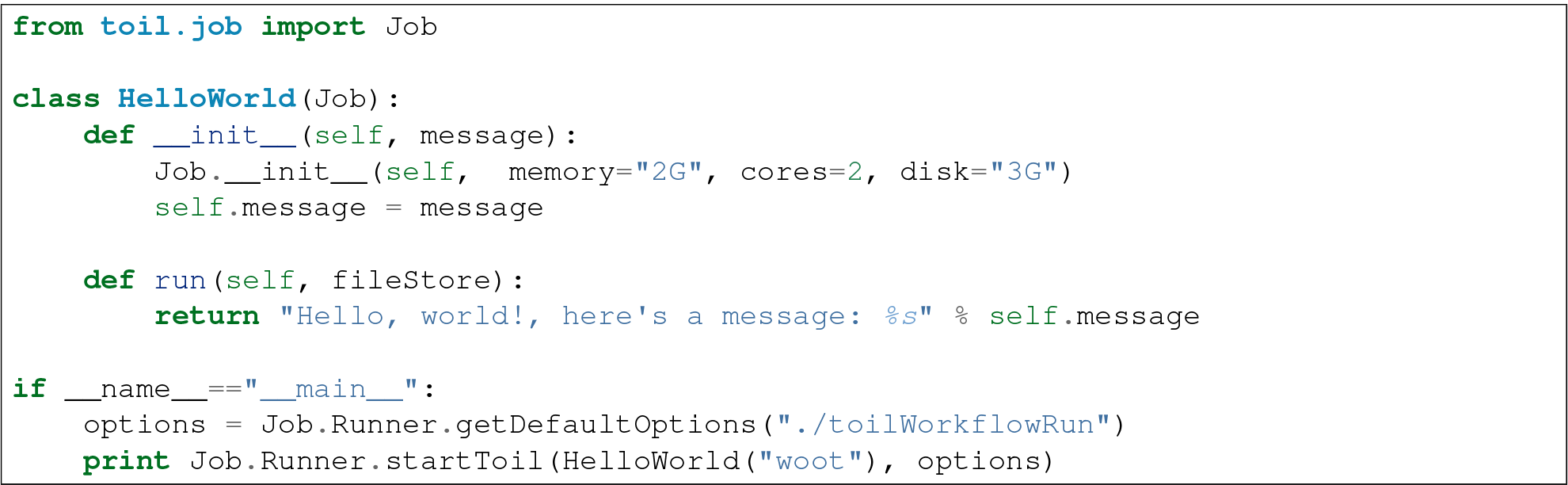

Alternatively, the more powerful `toil.common.Toil` class can be used to run and resume workflows. It is used as a context manager and allows for preliminary setup, such as staging of files into the job store on the leader node. An instance of the class is initialized by specifying an options object. The actual workflow is then invoked by calling the `toil.common.Toil.start()` method, passing the root job of the workflow, or, if a workflow is being restarted, `toil.common.Toil.restart()` should be used. Note that the context manager should have explicit if else branches addressing restart and non restart cases. The boolean value for these if else blocks is toil.options.restart.

For example:

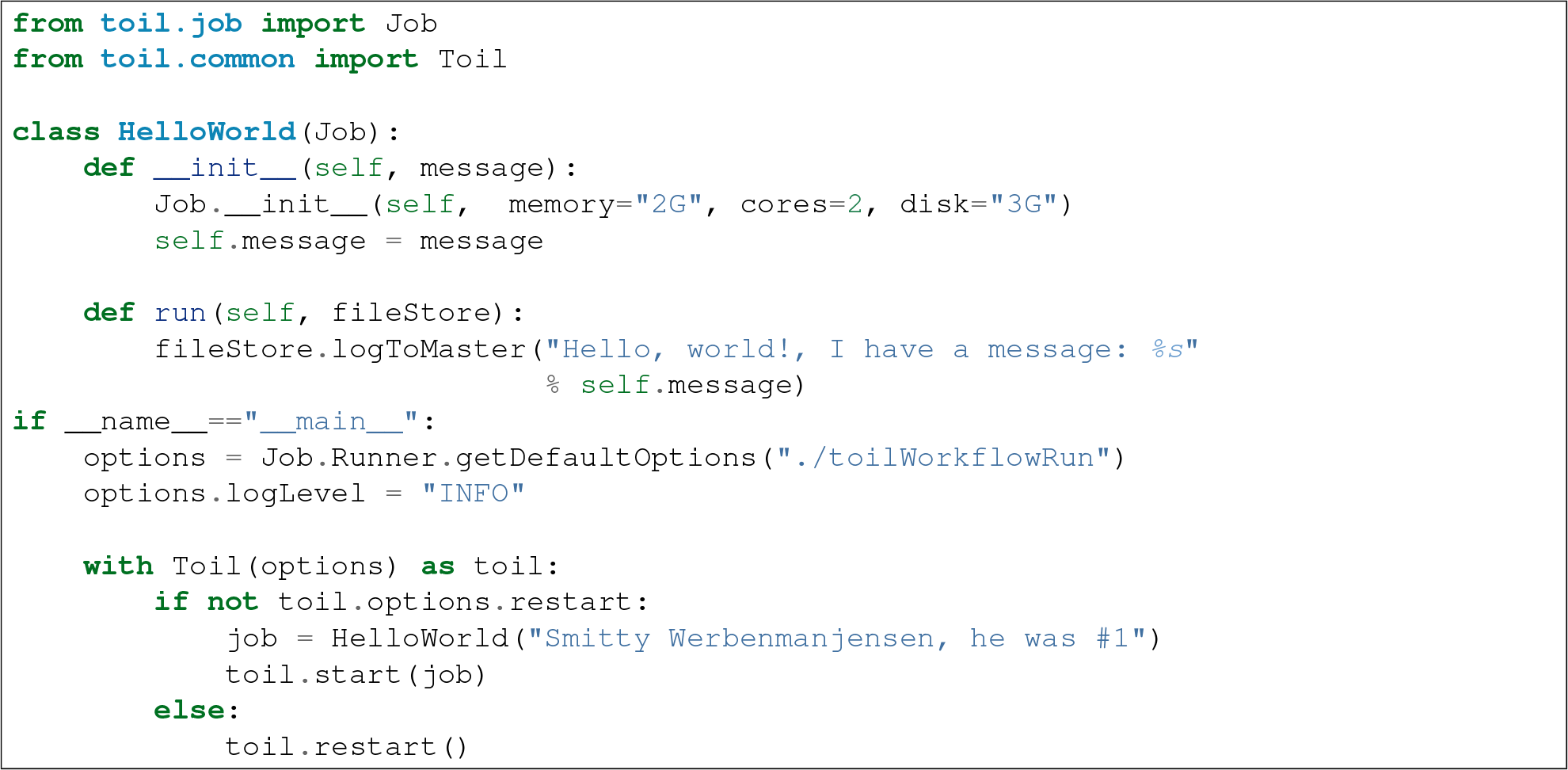

The call to `toil.job.Job.Runner.getDefaultOptions()` creates a set of default options for the workflow. The only argument is a description of how to store the workflow’s state in what we call a *job-store*. Here the job-storeis contained in a directory within the current working directory called “toilWorkflowRun”. Alternatively this string can encode other ways to store the necessary state, e.g. an S3 bucket or Azure object store location. By default the job-store is deleted if the workflow completes successfully.

The workflow is executed in the final line, which creates an instance of HelloWorld and runs it as a workflow. Note all Toil workflows start from a single starting job, referred to as the *root* job. The return value of the root job is returned as the result of the completed workflow (see promises below to see how this is a useful feature!).

### 6.4 Specifying arguments via the command line

To allow command line control of the options we can use the `toil.job.Job.Runner.getDefaultArgumentParser()` method to create a `argparse.ArgumentParser` object which can be used to parse command line options for a Toil script. For example:

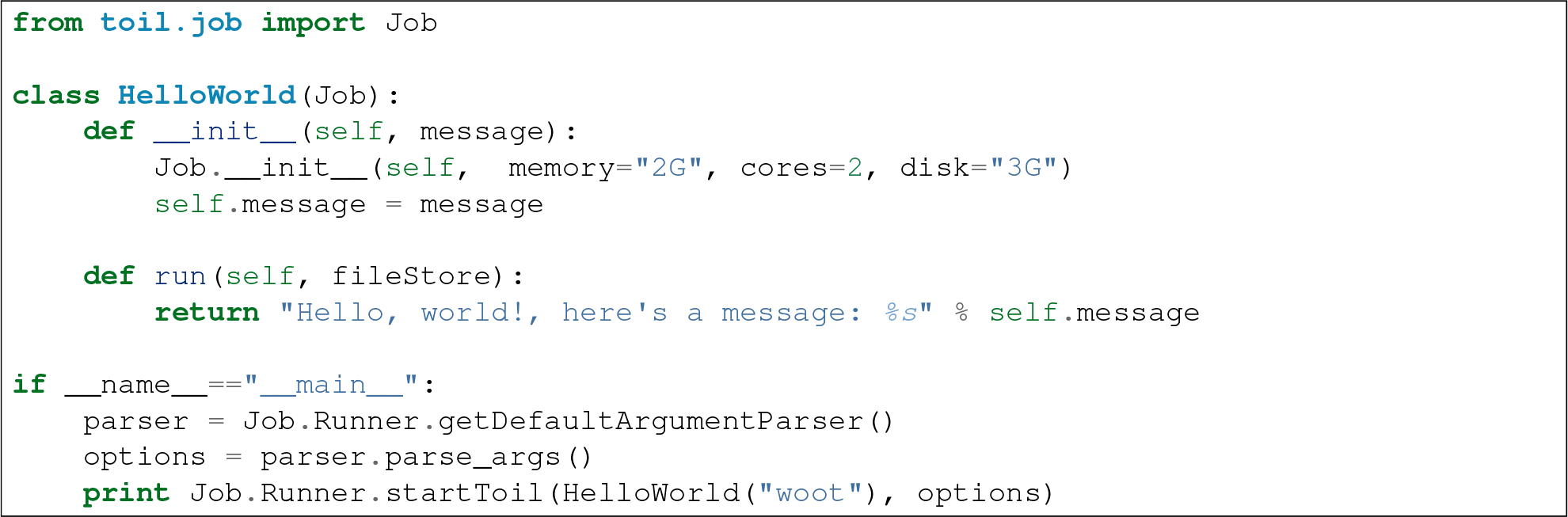

Creates a fully fledged script with all the options Toil exposed as command line arguments. Running this script with “-help” will print the full list of options.

Alternatively an existing `argparse.ArgumentParser` or `optparse.OptionParser` object can have Toil script command line options added to it with the `toil.job.Job.Runner.addToilOptions()` method.

### 6.5 Resuming a workflow

In the event that a workflow fails, either because of programmatic error within the jobs being run, or because of node failure, the workflow can be resumed. Workflows can only not be reliably resumed if the job-store itself becomes corrupt.

Critical to resumption is that jobs can be rerun, even if they have apparently completed successfully. Put succinctly, a user defined job should not corrupt its input arguments. That way, regardless of node, network or leader failure the job can be restarted and the workflow resumed.

To resume a workflow specify the “restart” option in the options object passed to `toil.job.Job.Runner.startToil()`. If node failures are expected it can also be useful to use the integer “retryCount” option, which will attempt to rerun a job retryCount number of times before marking it fully failed.

In the common scenario that a small subset of jobs fail (including retry attempts) within a workflow Toil will continue to run other jobs until it can do no more, at which point `toil.job.Job.Runner.startToil()` will raise a `toil.job.leader.FailedJobsException` exception. Typically at this point the user can decide to fix the script and resume the workflow or delete the job-store manually and rerun the complete workflow.

### 6.6 Functions and job functions

Defining jobs by creating class definitions generally involves the boilerplate of creating a constructor. To avoid this the classes `toil.job.FunctionWrappingJob` and `toil.job.JobFunctionWrappingTarget` allow functions to be directly converted to jobs. For example, the quick start example (repeated here):

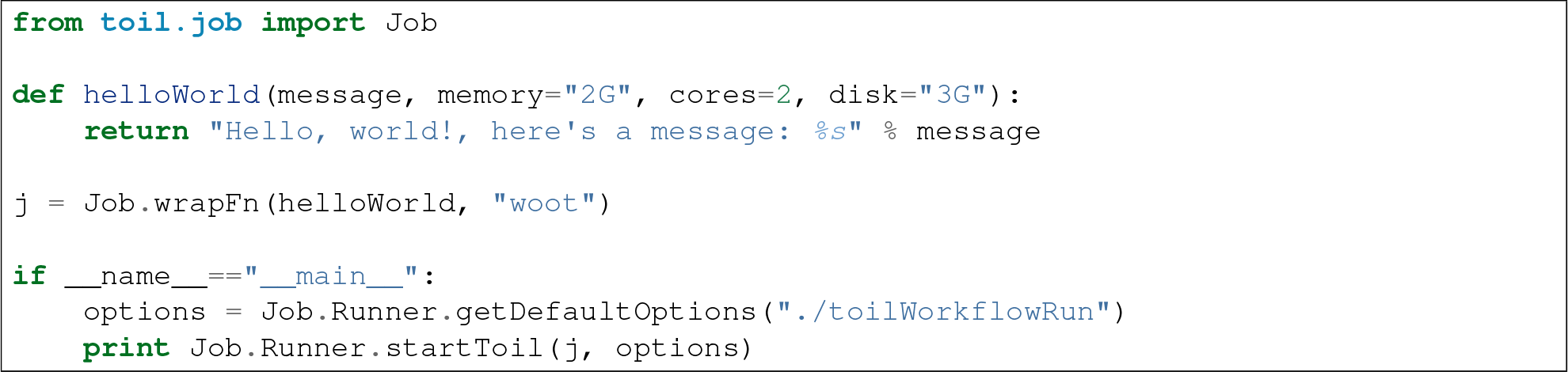

Is equivalent to the previous example, but using a function to define the job.

The function call:

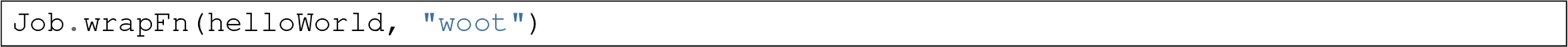

Creates the instance of the `toil.job.FunctionWrappingTarget` that wraps the function.

The keyword arguments *memory*, *cores* and *disk* allow resource requirements to be specified as before. Even if they are not included as keyword arguments within a function header they can be passed as arguments when wrapping a function as a job and will be used to specify resource requirements.

We can also use the function wrapping syntax to a *job function*, a function whose first argument is a reference to the wrapping job. Just like a *self* argument in a class, this allows access to the methods of the wrapping job, see `toil.job.JobFunctionWrappingTarget`. For example:

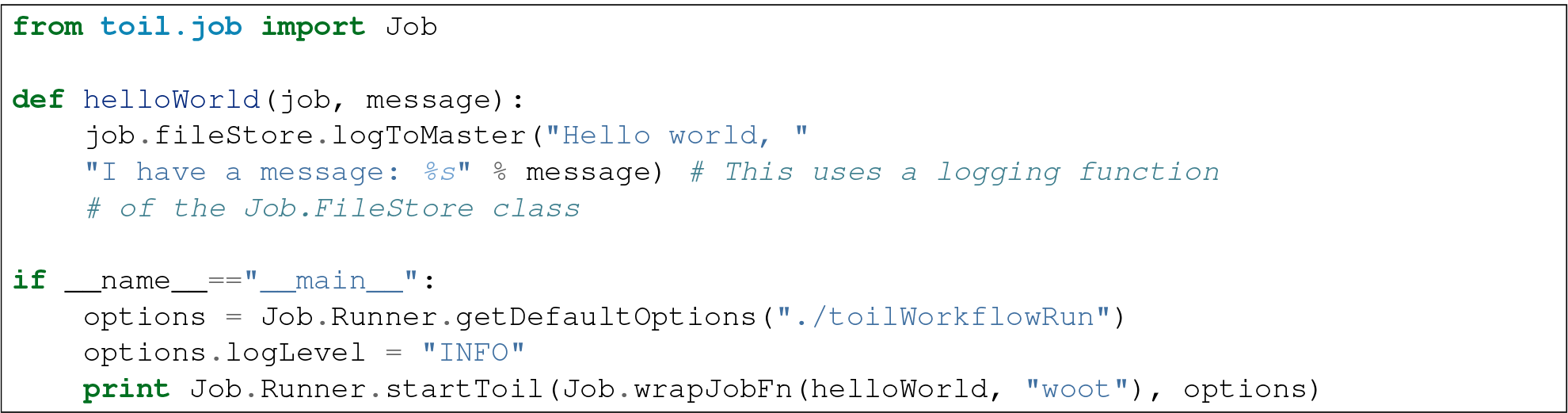

Here helloWorld2 is a job function. It accesses the `toil.job.Job.FileStore` attribute of the job to log a message that will be printed to the output console. Here the only subtle difference to note is the line:

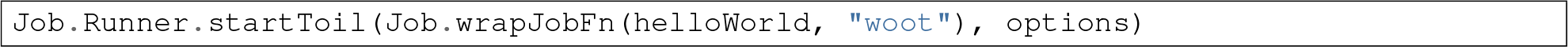

Which uses the function `toil.job.Job.wrapJobFn()` to wrap the job function instead of `toil.job.Job.wrapFn()` which wraps a vanilla function.

### 6.7 Workflows with multiple jobs

A *parent* job can have *child* jobs and *follow-on* jobs. These relationships are specified by methods of the job class, e.g. `toil.job.Job.addChild()` and `toil.job.Job.addFollowOn()`.

Considering a set of jobs the nodes in a job graph and the child and follow-on relationships the directed edges of the graph, we say that a job B that is on a directed path of child/follow-on edges from a job A in the job graph is a *successor* of A, similarly A is a *predecessor* of B.

A parent job’s child jobs are run directly after the parent job has completed, and in parallel. The follow-on jobs of a job are run after its child jobs and their successors have completed. They are also run in parallel. Follow-ons allow the easy specification of cleanup tasks that happen after a set of parallel child tasks. The following shows a simple example that uses the earlier helloWorld job function:

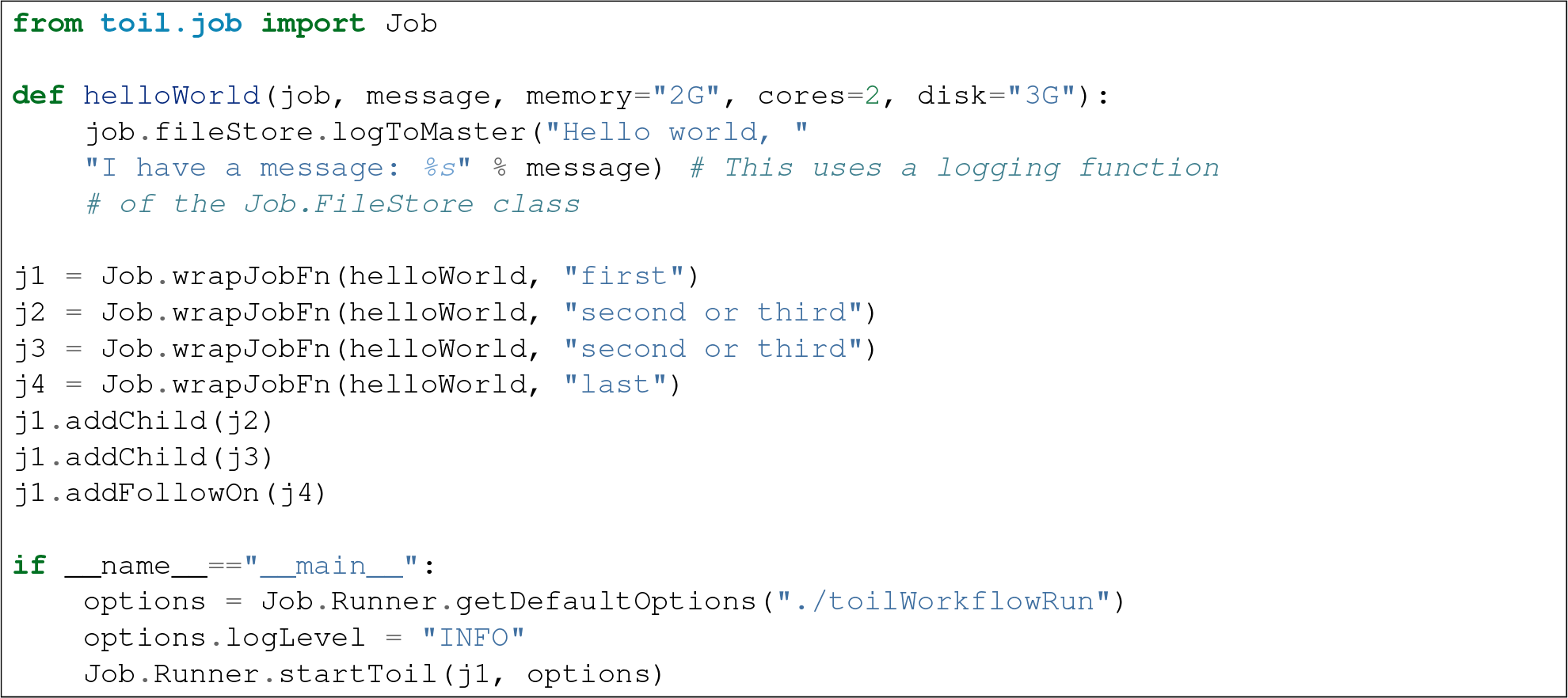

In the example four jobs are created, first j1 is run, then j2 and j3 are run in parallel as children of j1, finally j4 is run as a follow-on of j1.

There are multiple short hand functions to achieve the same workflow, for example:

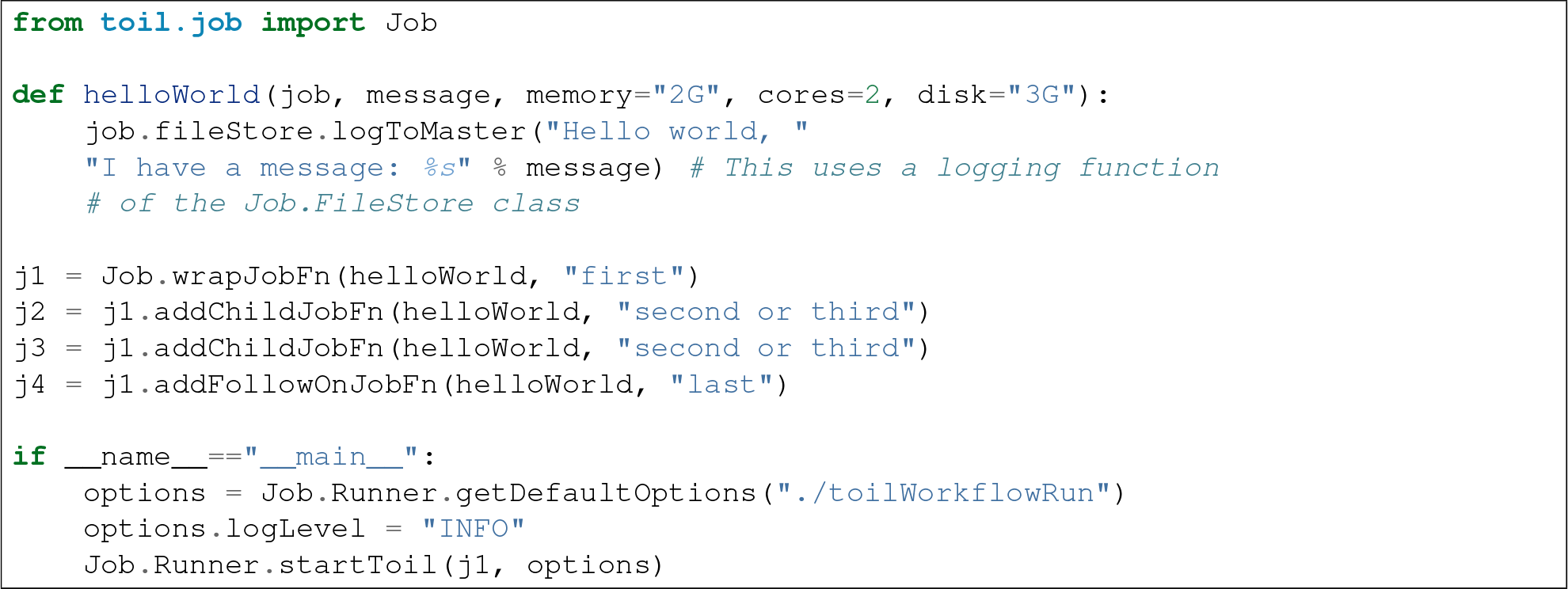

Equivalently defines the workflow, where the functions `toil.job.Job.addChildJobFn()` and `toil.job.Job.addFollowOnJobFn()` are used to create job functions as children or follow-ons of an earlier job.

Jobs graphs are not limited to trees, and can express arbitrary directed acylic graphs. For a precise definition of legal graphs see `toil.job.Job.checkJobGraphForDeadlocks()`. The previous example could be specified as a DAG as follows:

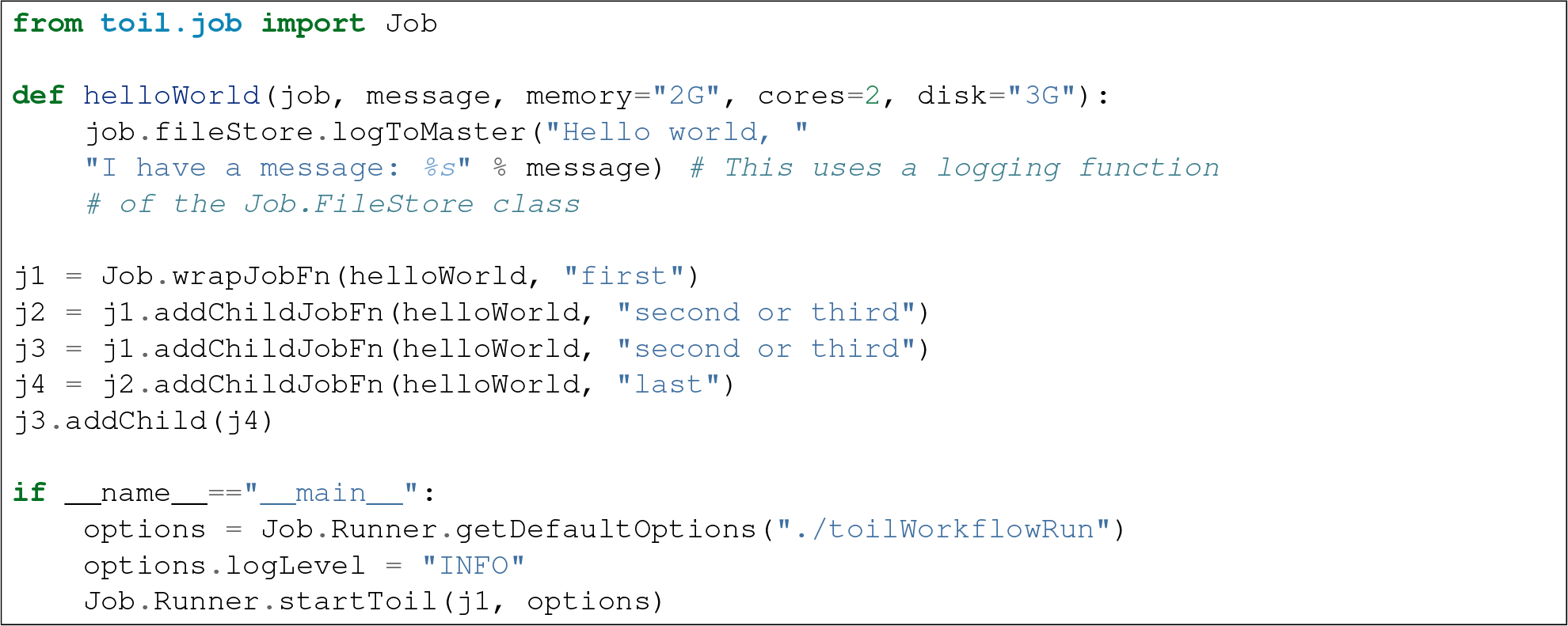

Note the use of an extra child edge to make j4 a child of both j2 and j3.

### 6.8 Dynamic job creation

The previous examples show a workflow being defined outside of a job. However, Toil also allows jobs to be created dynamically within jobs. For example:

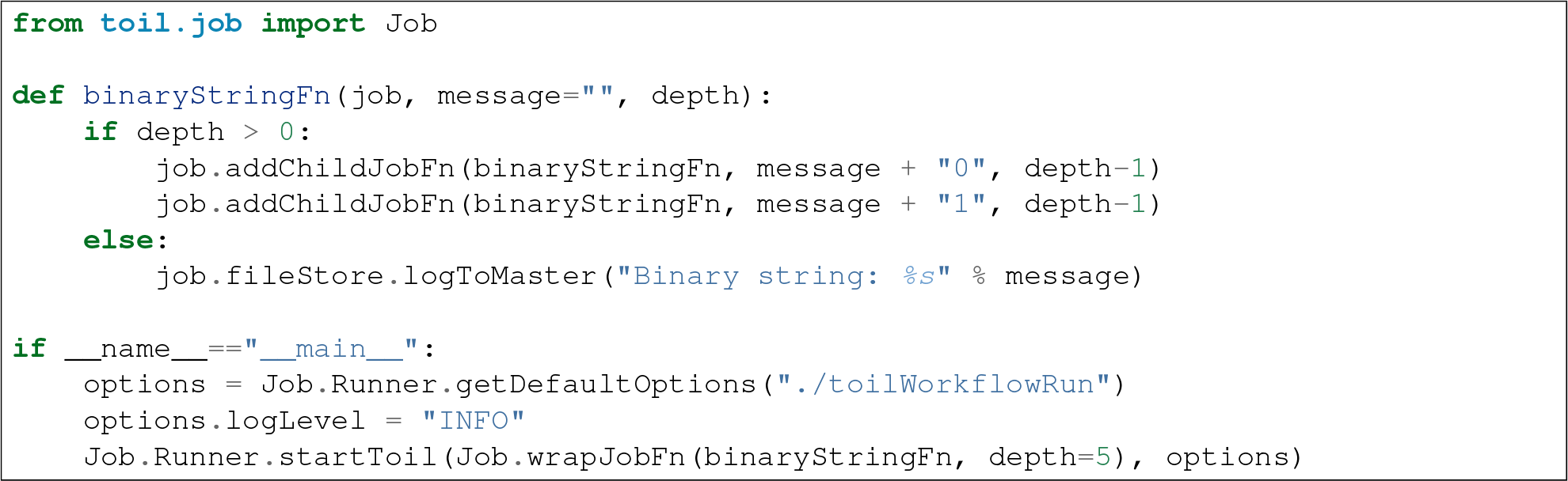

The binaryStringFn logs all possible binary strings of length n (here n=5), creating a total of 2^(n+2) − 1 jobs dynamically and recursively. Static and dynamic creation of jobs can be mixed in a Toil workflow, with jobs defined within a job or job function being created at run-time.

### 6.9 Promises

The previous example of dynamic job creation shows variables from a parent job being passed to a child job. Such forward variable passing is naturally specified by recursive invocation of successor jobs within parent jobs. This can also be achieved statically by passing around references to the return variables of jobs. In Toil this is achieved with promises, as illustrated in the following example:

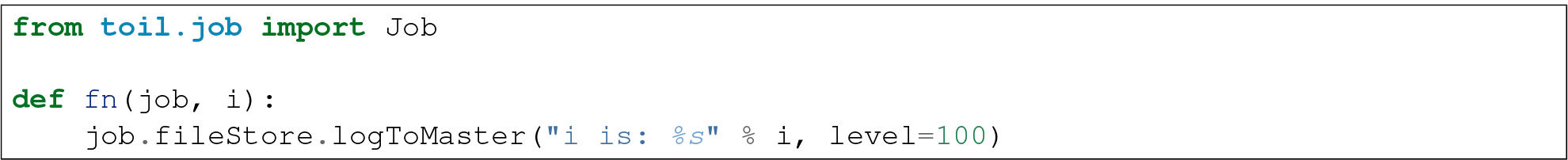

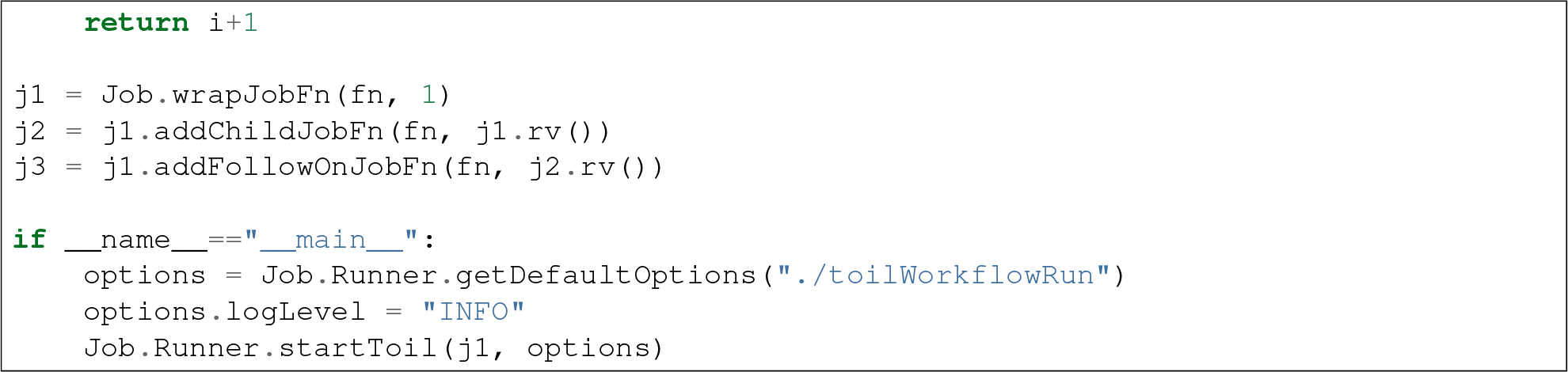

Running this workflow results in three log messages from the jobs: “i is 1” from *j1*, “i is 2” from *j2* and “i is 3” from j3.

The return value from the first job is *promised* to the second job by the call to `toil.job.Job.rv()` in the line:

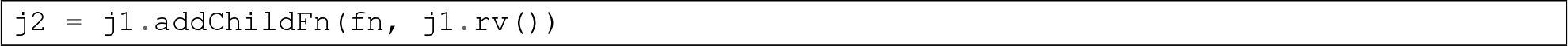

The value of *j1.rv()* is a *promise,* rather than the actual return value of the function, because j1 for the given input has at that point not been evaluated. A promise (`toil.job.Promise`) is essentially a pointer to the return value that is replaced by the actual return value once it has been evaluated. Therefore when j2 is run the promise becomes 2.

Promises can be quite useful. For example, we can combine dynamic job creation with promises to achieve a job creation process that mimics the functional patterns possible in many programming languages:

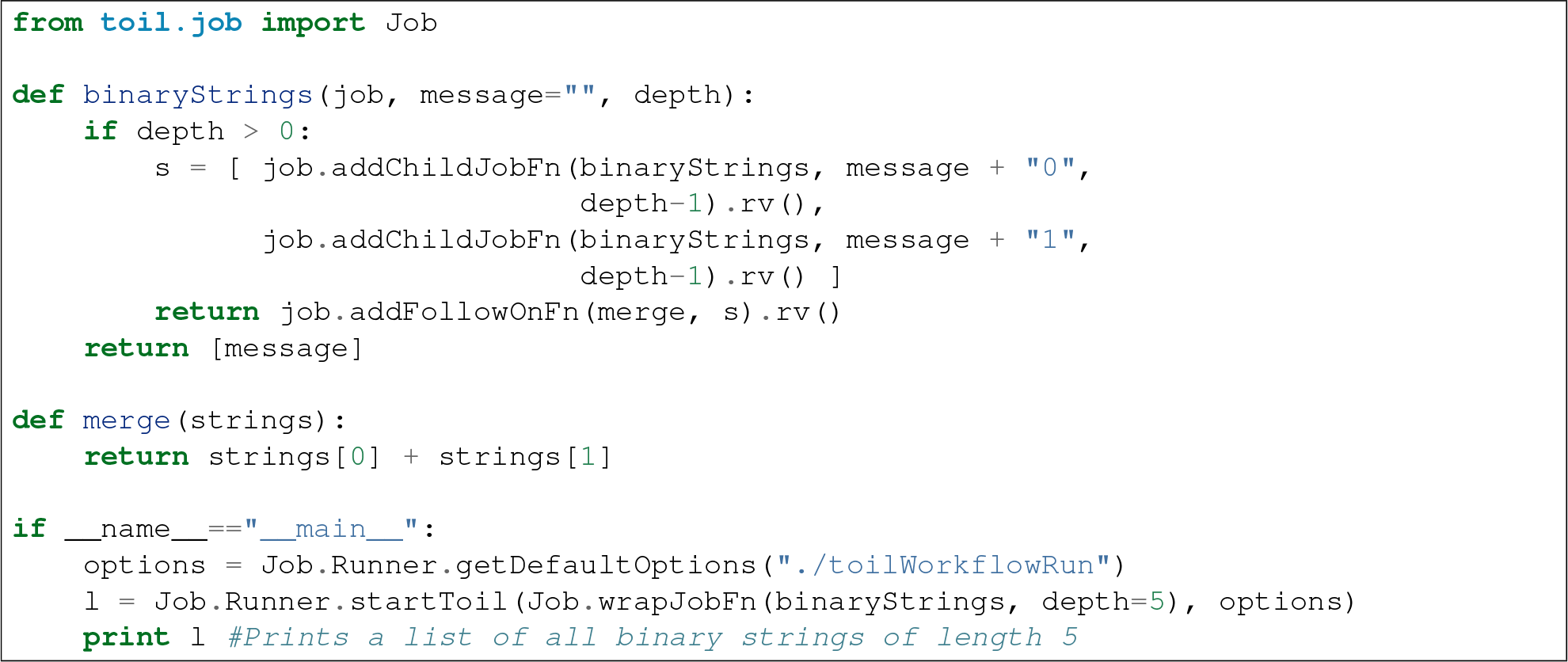

The return value *l* of the workflow is a list of all binary strings of length 10, computed recursively. Although a toy example, it demonstrates how closely Toil workflows can mimic typical programming patterns.

### 6.10 Managing files within a workflow

It is frequently the case that a workflow will want to create files, both persistent and temporary, during its run. The `toil.job.Job.FileStore` class is used by jobs to manage these files in a manner that guarantees cleanup and resumption on failure.

The `toil.job.Job.run()` method has a file-store instance as an argument. The following example shows how this can be used to create temporary files that persist for the length of the job, be placed in a specified local disk of the node and that will be cleaned up, regardless of failure, when the job finishes:

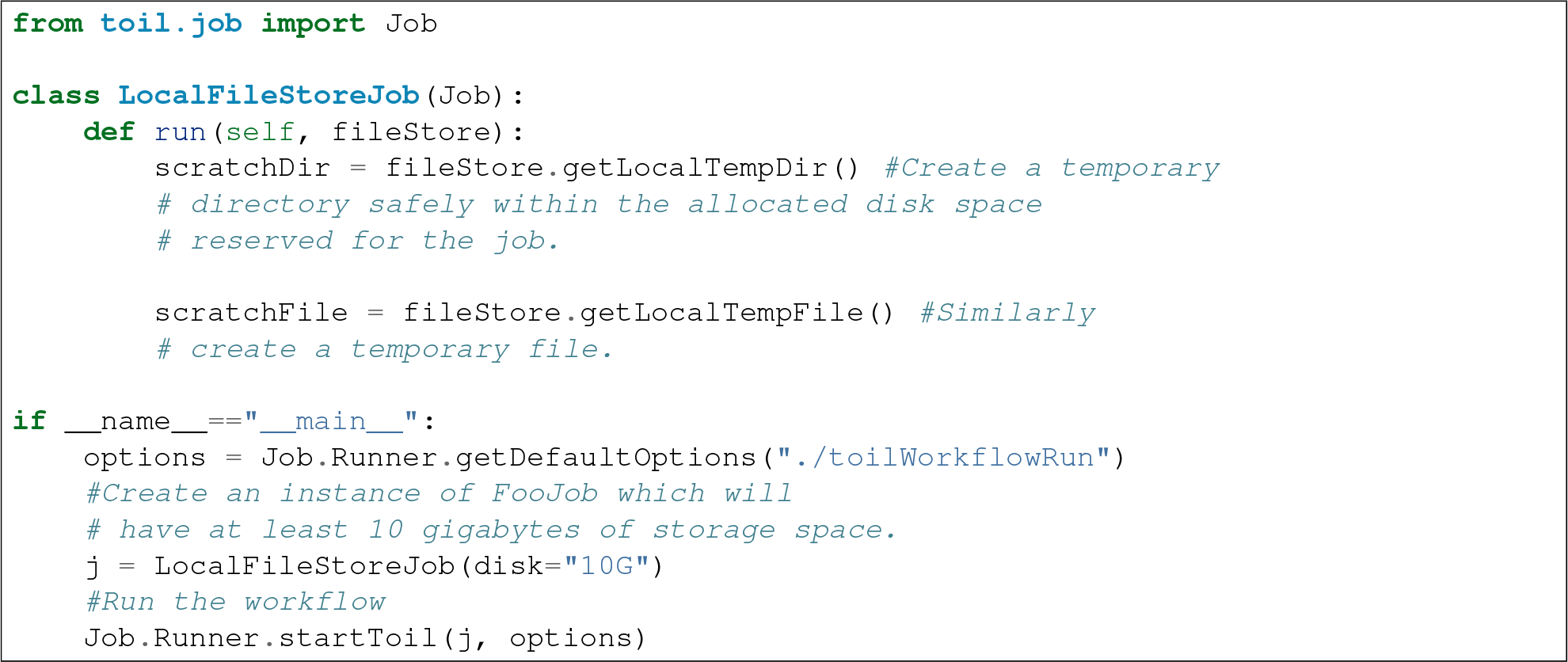

Job functions can also access the file-store for the job. The equivalent of the LocalFileStoreJob class is equivalently:

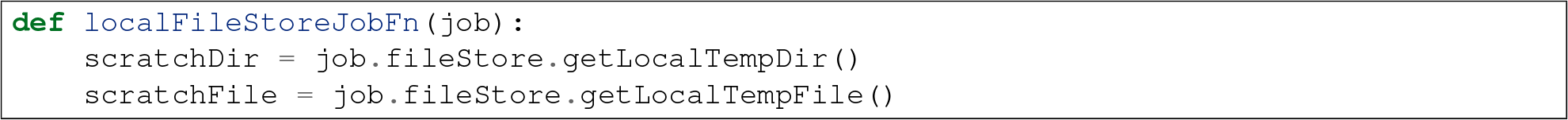

Note that the fileStore attribute is accessed as an attribute of the job argument.

In addition to temporary files that exist for the duration of a job, the file-store allows the creation of files in a *global* store, which persists during the workflow and are globally accessible (hence the name) between jobs. For example:

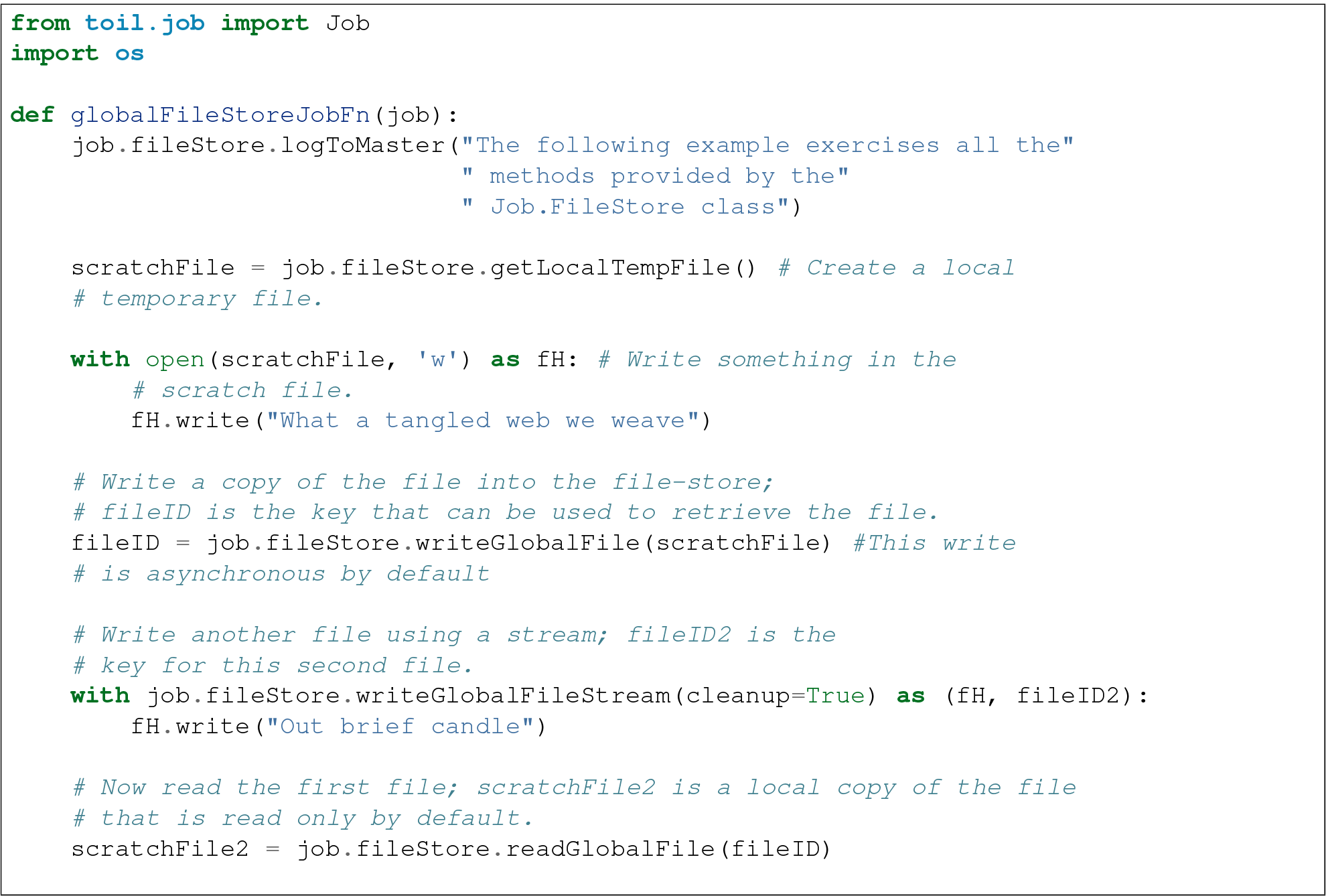

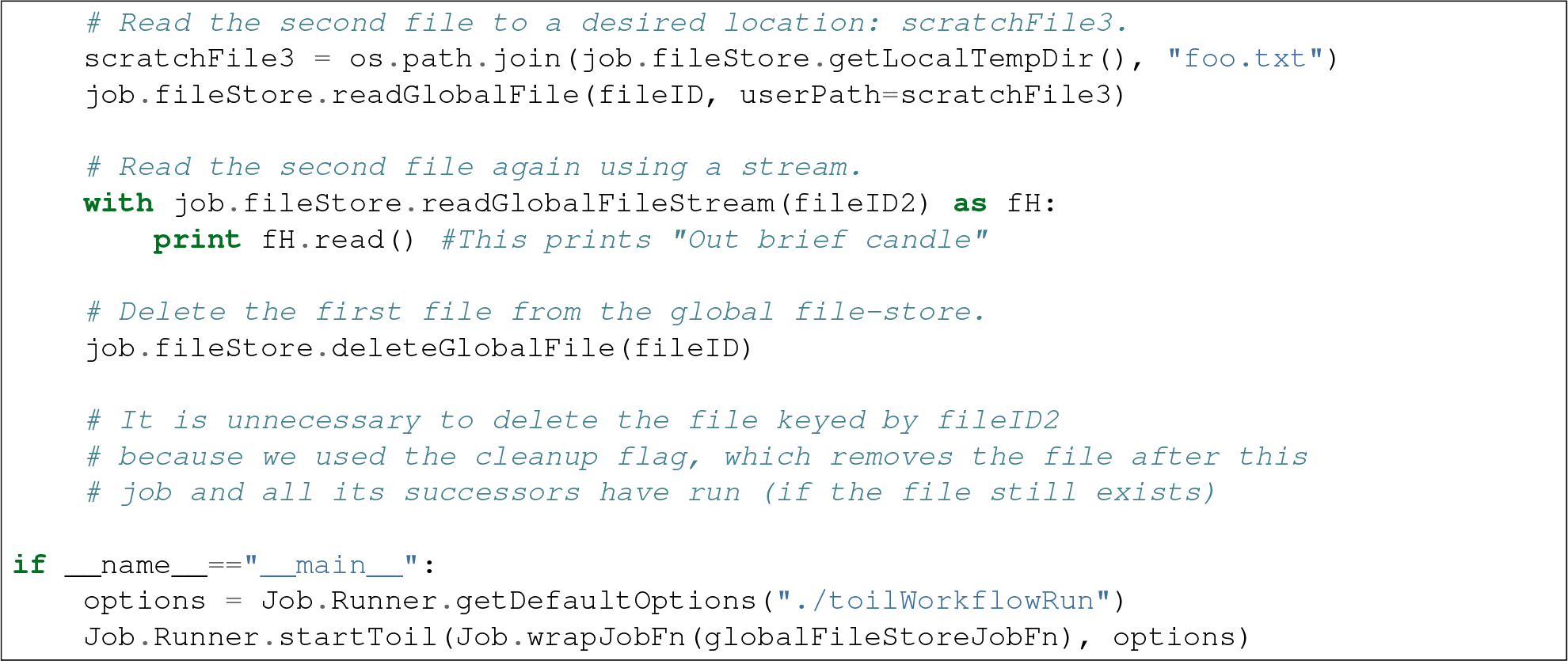

The example demonstrates the global read, write and delete functionality of the file-store, using both local copies of the files and streams to read and write the files. It covers all the methods provided by the file-store interface.

What is obvious is that the file-store provides no functionality to update an existing “global” file, meaning that files are, barring deletion, immutable. Also worth noting is that there is no file system hierarchy for files in the global file store. These limitations allow us to fairly easily support different object stores and to use caching to limit the amount of network file transfer between jobs.

#### 6.10.1 Staging of files into the job store

External files can be imported into or exported out of the job store prior to running a workflow when the `toil.common.Toil` context manager is used on the leader. The context manager provides methods `toil.common.Toil.importFile()`, and `toil.common.Toil.exportFile()` for this purpose. The destination and source locations of such files are described with URLs passed to the two methods. A list of the currently supported URLs can be found at `toil.jobStores.abstractJobStore.AbstractJobStore.importFile()`. To import an external file into the job store as a shared file, pass the optional `sharedFileName` parameter to that method.

If a workflow fails for any reason an imported file acts as any other file in the job store. If the workflow was configured such that it not be cleaned up on a failed run, the file will persist in the job store and needs not be staged again when the workflow is resumed.

Example:

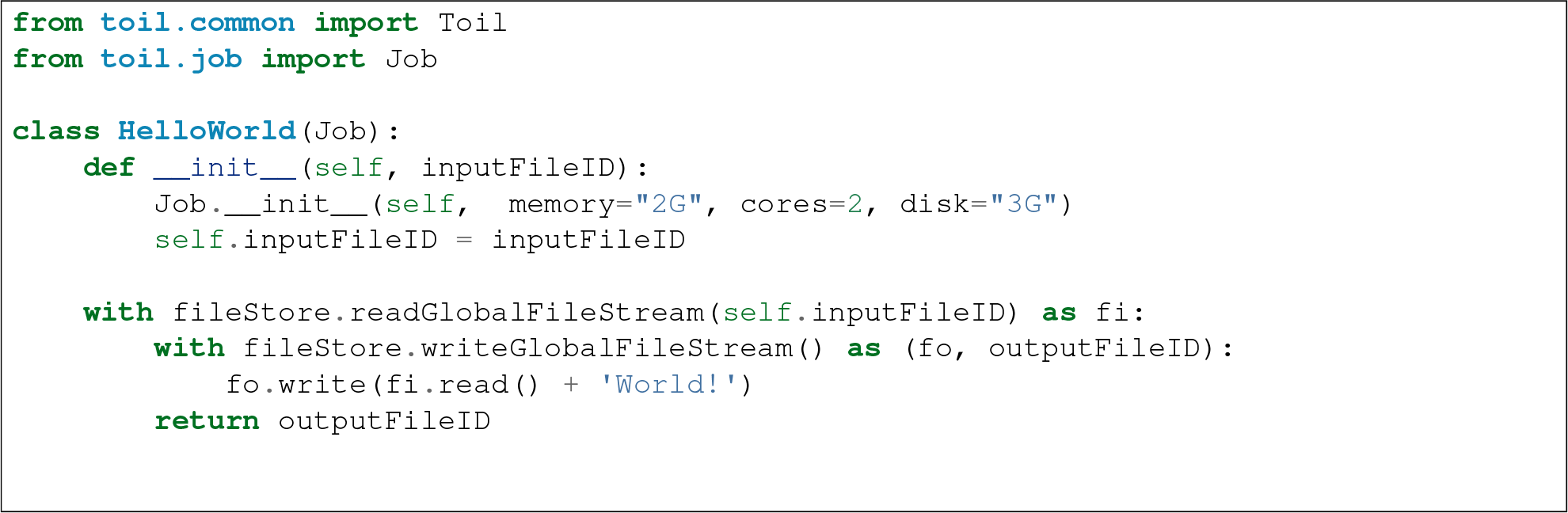

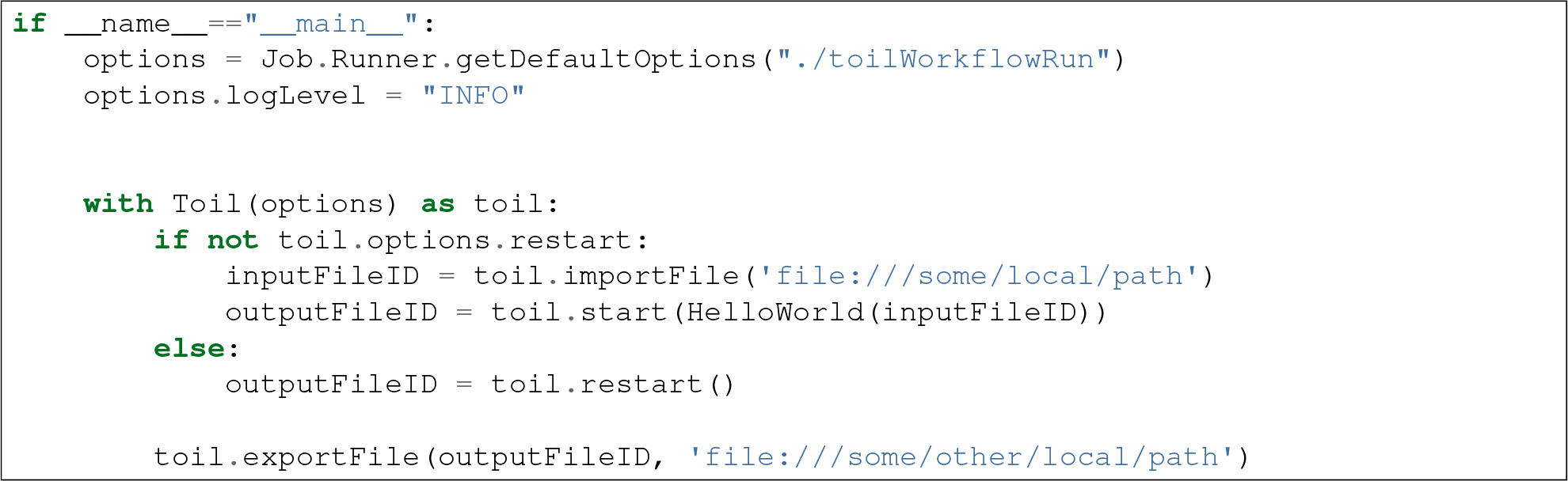

### 6.11 Services

It is sometimes desirable to run *services*, such as a database or server, concurrently with a workflow. The `toil.job.Job.Service` class provides a simple mechanism for spawning such a service within a Toil workflow, allowing precise specification of the start and end time of the service, and providing start and end methods to use for initialization and cleanup. The following simple, conceptual example illustrates how services work:

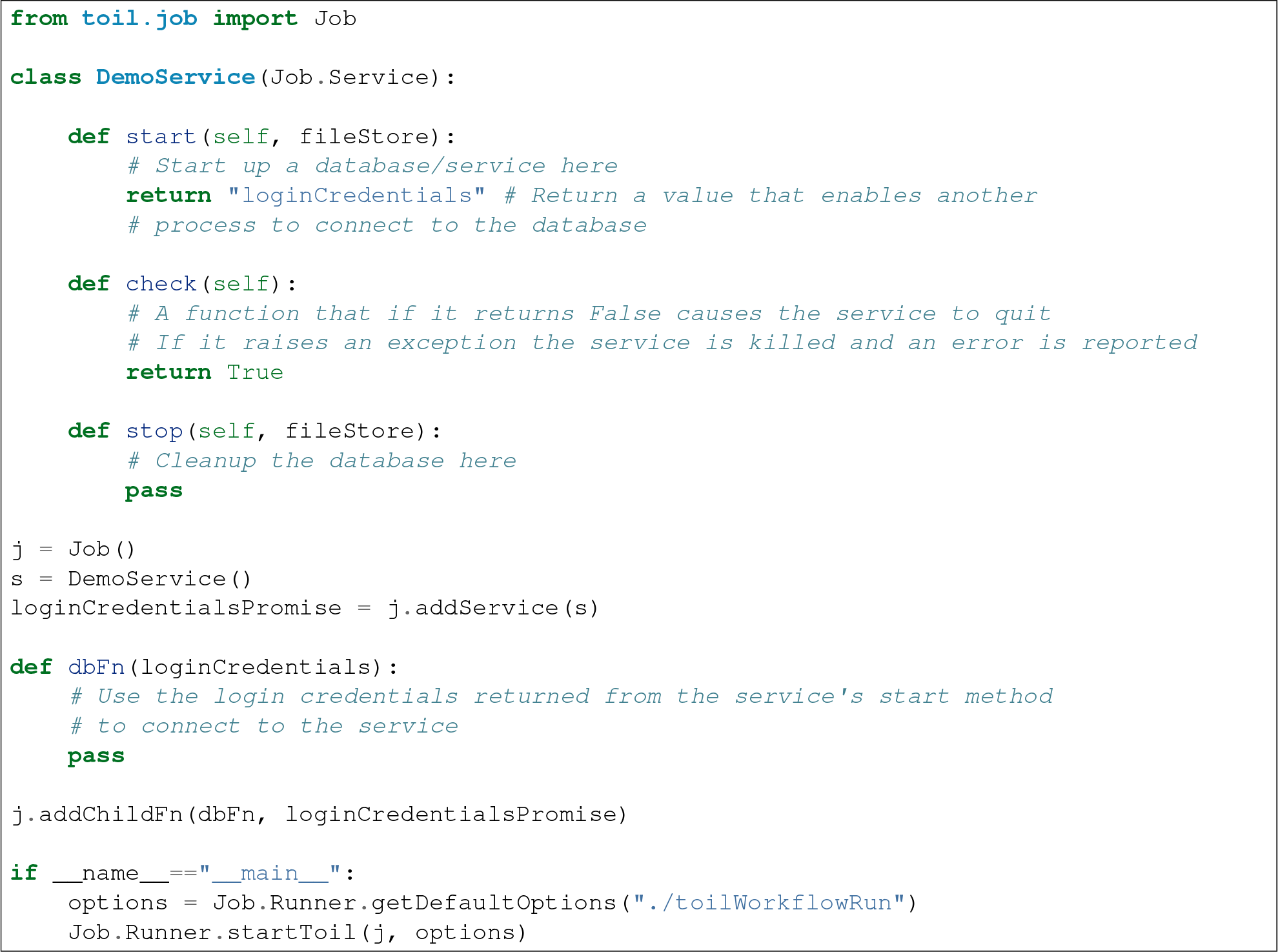

In this example the DemoService starts a database in the start method, returning an object from the start method indicating how a client job would access the database. The service’s stop method cleans up the database, while the service’s check method is polled periodically to check the service is alive.

A DemoService instance is added as a service of the root job *j*, with resource requirements specified. The return value from `toil.job.Job.addService()` is a promise to the return value of the service’s start method. When the promised is fulfilled it will represent how to connect to the database. The promise is passed to a child job of j, which uses it to make a database connection. The services of a job are started before any of its successors have been run and stopped after all the successors of the job have completed successfully.

Multiple services can be created per job, all run in parallel. Additionally, services can define sub-services using `toil.job.Job.Service.addChild()`. This allows complex networks of services to be created, e.g. Apache Spark clusters, within a workflow.

### 6.12 Checkpoints

Services complicate resuming a workflow after failure, because they can create complex dependencies between jobs. For example, consider a service that provides a database that multiple jobs update. If the database service fails and loses state, it is not clear that just restarting the service will allow the workflow to be resumed, because jobs that created that state may have already finished. To get around this problem Toil supports “checkpoint” jobs, specified as the boolean keyword argument “checkpoint” to a job or wrapped function, e.g.:

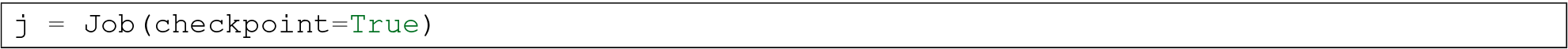

A checkpoint job is rerun if one or more of its successors fails its retry attempts, until it itself has exhausted its retry attempts. Upon restarting a checkpoint job all its existing successors are first deleted, and then the job is rerun to define new successors. By checkpointing a job that defines a service, upon failure of the service the database and the jobs that access the service can be redefined and rerun.

To make the implementation of checkpoint jobs simple, a job can only be a checkpoint if when first defined it has no successors, i.e. it can only define successors within its run method.

### 6.13 Encapsulation

Let A be a root job potentially with children and follow-ons. Without an encapsulated job the simplest way to specify a job B which runs after A and all its successors is to create a parent of A, call it Ap, and then make B a follow-on of Ap. e.g.:

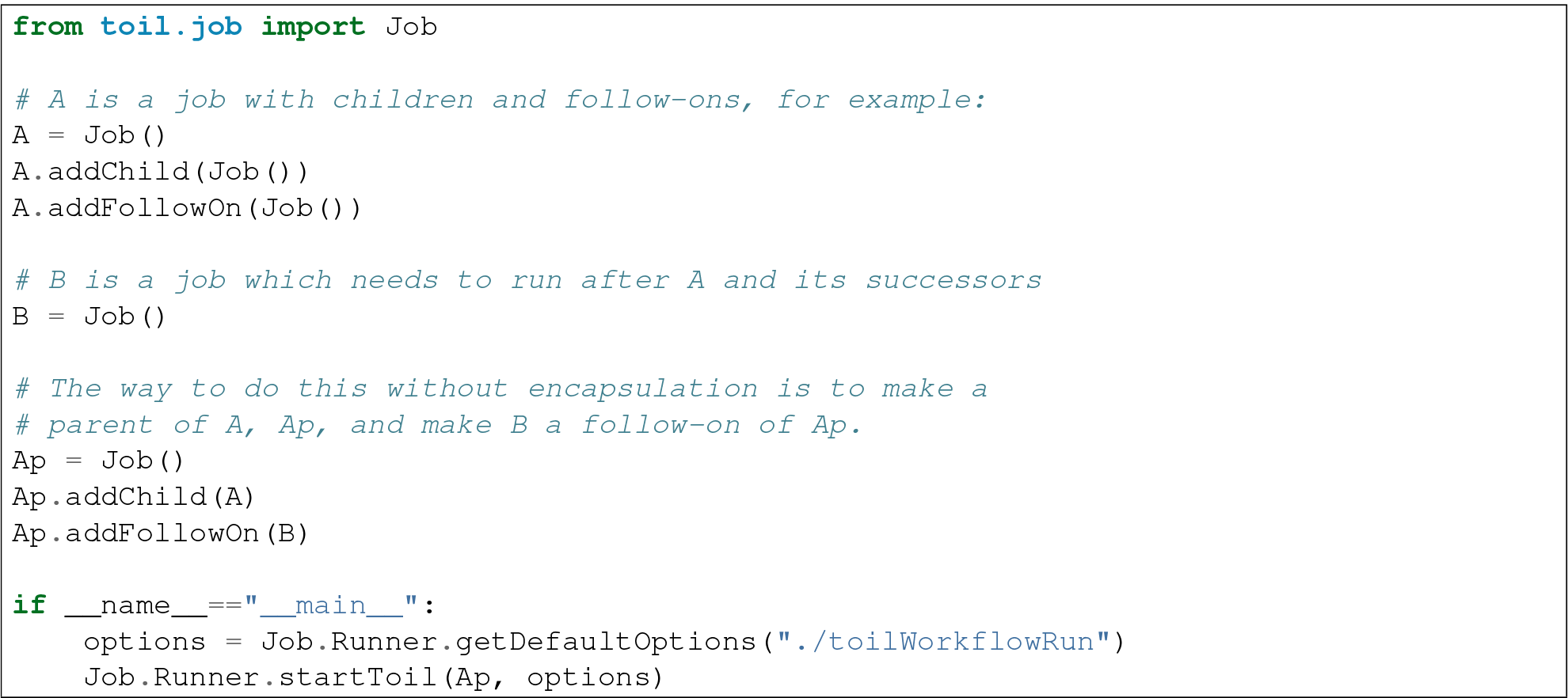

An *encapsulated* job of E(A) of A saves making Ap, instead we can write:

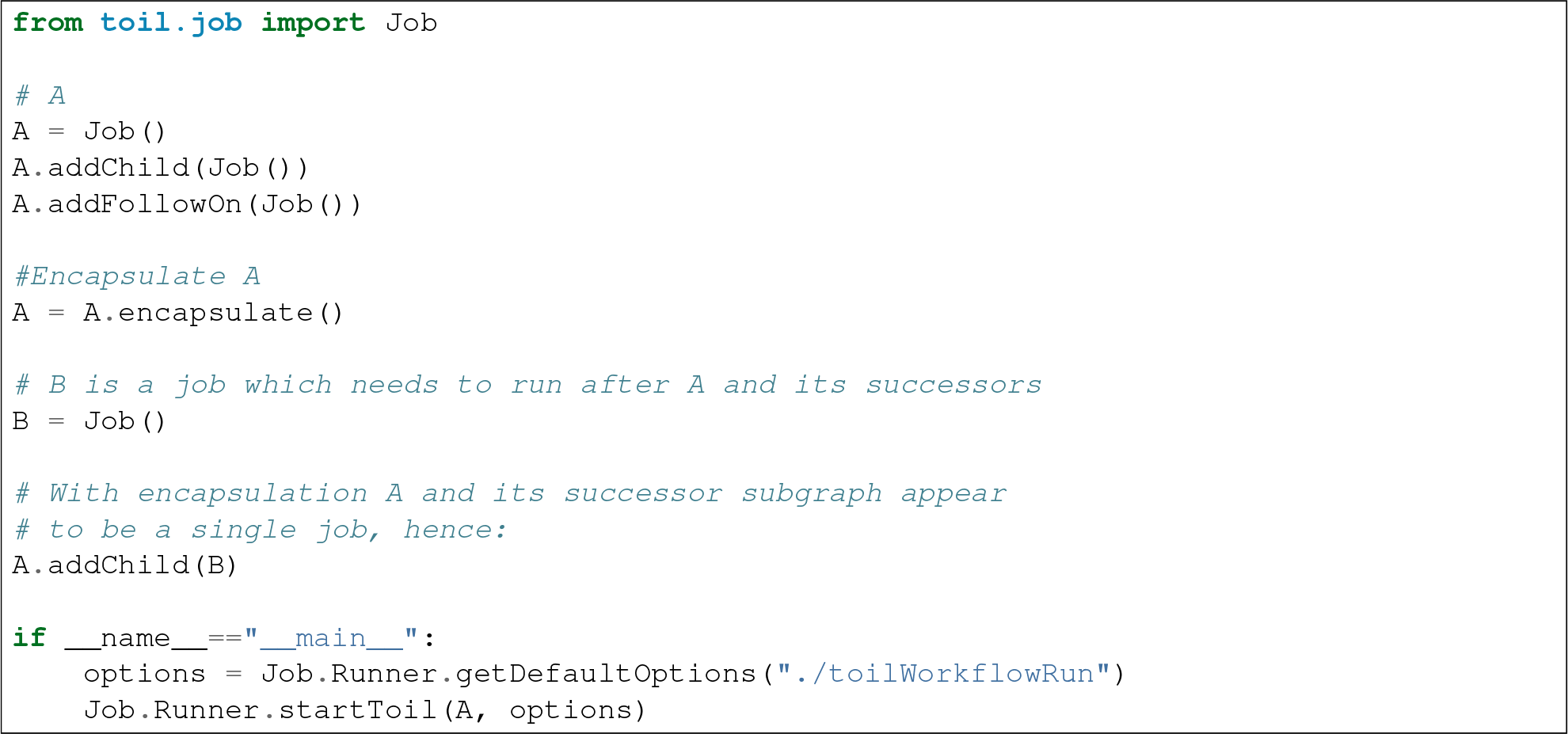

Note the call to `toil.job.Job.encapsulate()` creates the `toil.job.Job.EncapsulatedJob`.

## TOIL API

### 7.1 Job methods

Jobs are the units of work in Toil which are composed into workflows.

### 7.2 Job.FileStore

The FileStore is an abstraction of a Toil run’s shared storage.

### 7.3 Job.Runner

The Runner contains the methods needed to configure and start a Toil run.

### 7.4 Toil

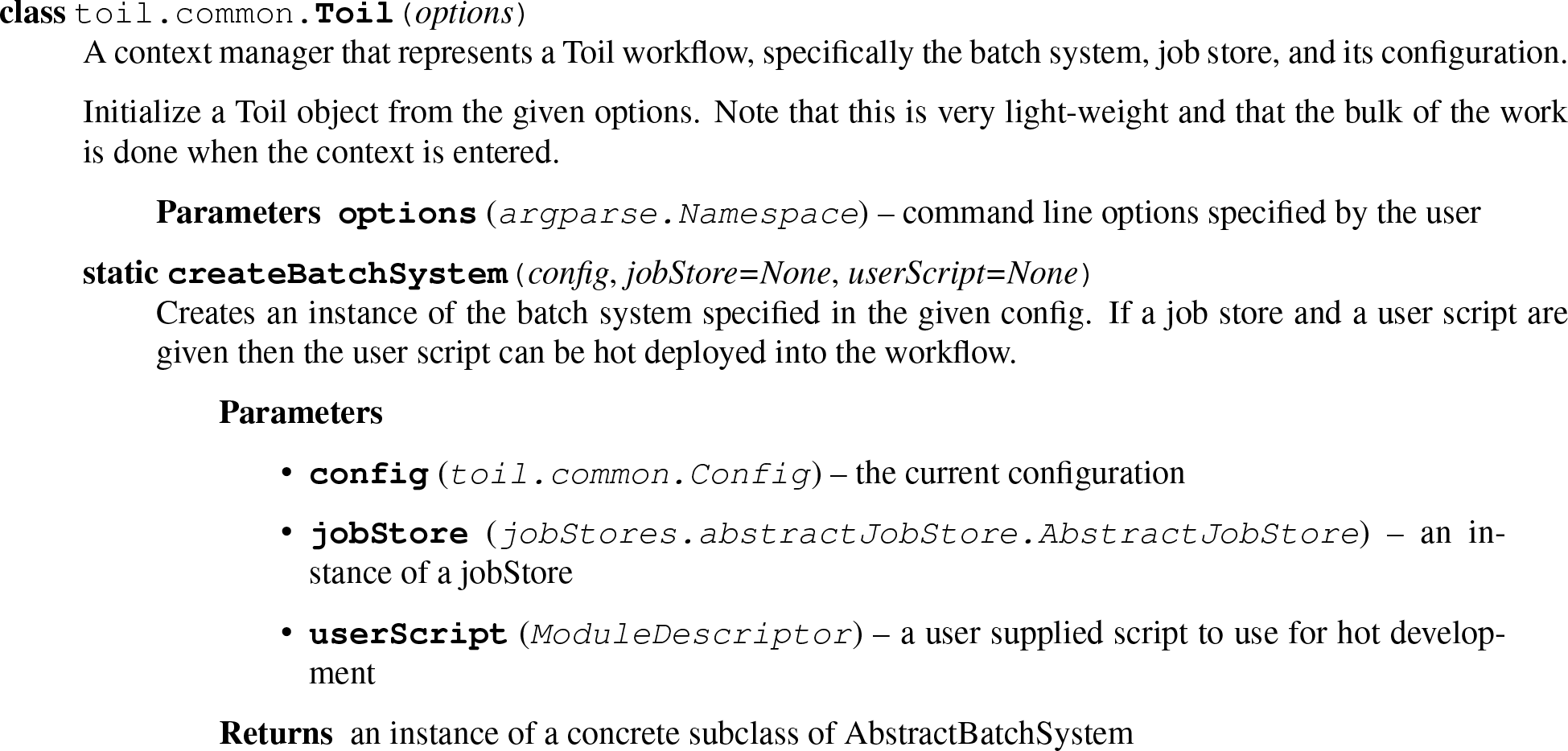

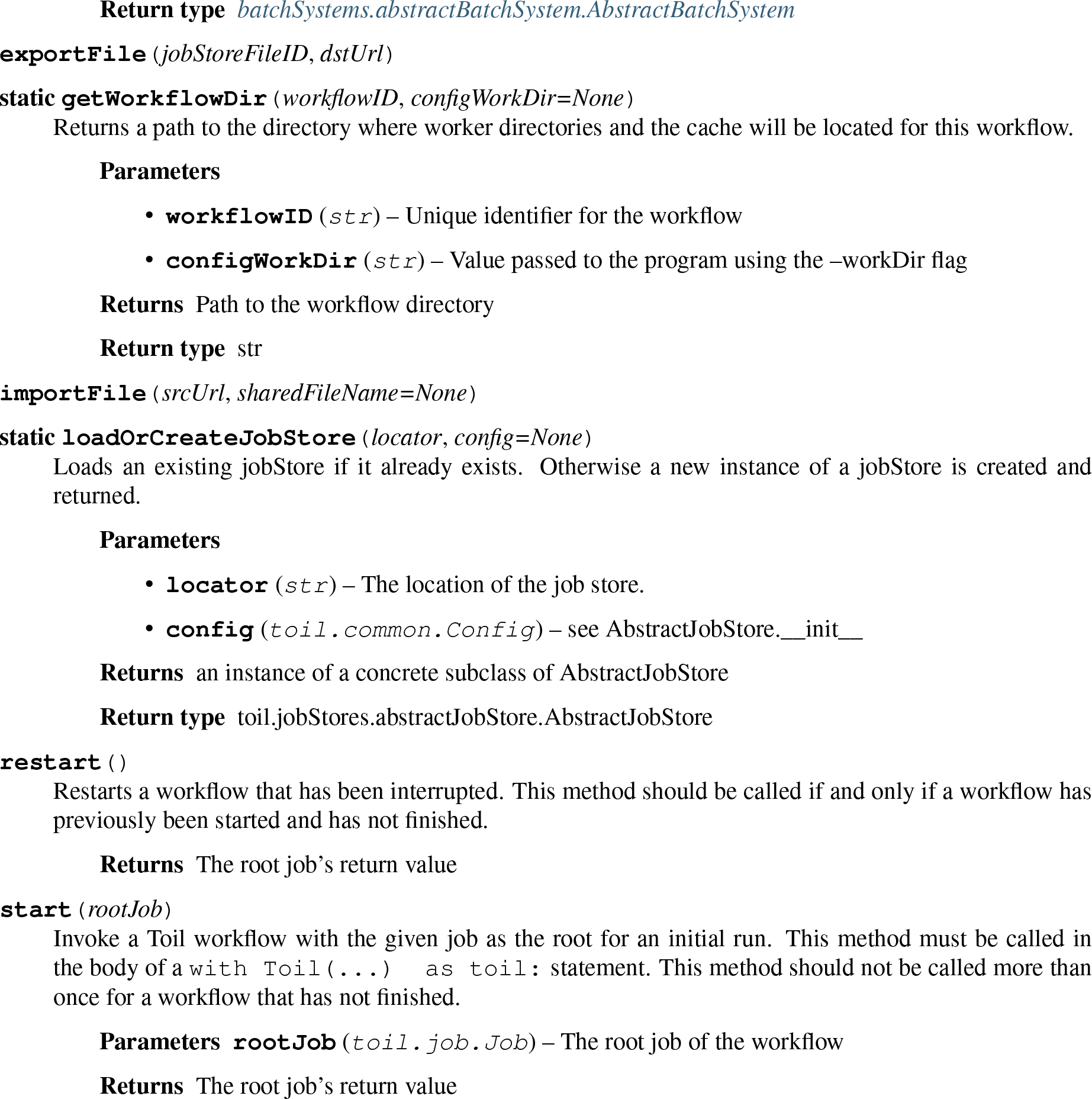

### 7.5 Job.Service

The Service class allows databases and servers to be spawned within a Toil workflow.

### 7.6 FunctionWrappingJob

The subclass of Job for wrapping user functions.

### 7.7 JobFunctionWrappingJob

The subclass of FunctionWrappingJob for wrapping user job functions.

### 7.8 EncapsulatedJob

The subclass of Job for *encapsulating* a job, allowing a subgraph of jobs to be treated as a single job.

### 7.9 Promise

The class used to reference return values of jobs/services not yet run/started.

### 7.10 Exceptions

Toil specific exceptions.

## TOIL ARCHITECTURE

The following diagram layouts out the software architecture of Toil.

**Fig. 8.1: Figure 1:**
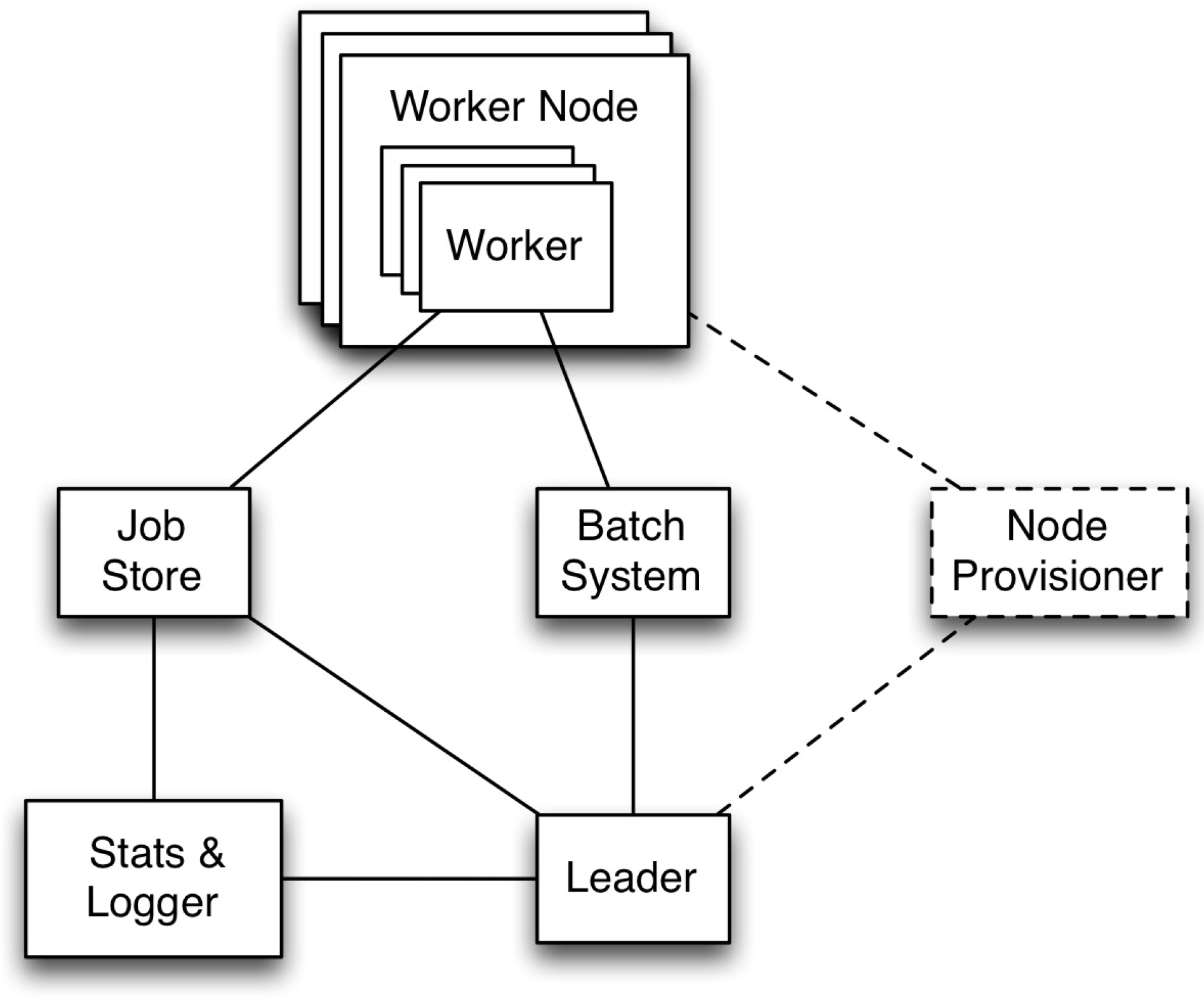
The basic components of the toil architecture. Note the node provisioning is coming soon.

These components are described below:

- **the leader:** The leader is responsible for deciding which jobs should be run. To do this it traverses the job graph. Currently this is a single threaded process, but we make aggressive steps to prevent it becoming a bottleneck (see *Read-only Leader* described below).
- **the job-store:** Handles all files shared between the components. Files in the job-store are the means by which the state of the workflow is maintained. Each job is backed by a file in the job store, and atomic updates to this state are used to ensure the workflow can always be resumed upon failure. The job-store can also store all user files, allowing them to be shared between jobs. The job-store is defined by the abstract class `toil.jobStores.AbstractJobStore`. Multiple implementations of this class allow Toil to support different back-end file stores, e.g.: S3, network file systems, Azure file store, etc.
- **workers:** The workers are temporary processes responsible for running jobs, one at a time per worker. Each worker process is invoked with a job argument that it is responsible for running. The worker monitors this job and reports back success or failure to the leader by editing the job’s state in the file-store. If the job defines successor jobs the worker may choose to immediately run them (see *Job Chaining* below).
- **the batch-system:** Responsible for scheduling the jobs given to it by the leader, creating a worker command for each job. The batch-system is defined by the abstract class class `toil.batchSystems.AbstractBatchSystem`. Toil uses multiple existing batch systems to schedule jobs, including Apache Mesos, GridEngine and a multi-process single node implementation that allows workflows to be run without any of these frameworks. Toil can therefore fairly easily be made to run a workflow using an existing cluster.
- **the node provisioner:** Creates worker nodes in which the batch system schedules workers. This is currently being developed. It is defined by the abstract class `toil.provisioners.AbstractProvisioner`.
- **the statistics and logging monitor:** Monitors logging and statistics produced by the workers and reports them. Uses the job-store to gather this information.

### 8.1 Optimizations

Toil implements lots of optimizations designed for scalability. Here we detail some of the key optimizations.

#### 8.1.1 Read-only leader

The leader process is currently implemented as a single thread. Most of the leader‘s tasks revolve around processing the state of jobs, each stored as a file within the job-store. To minimise the load on this thread, each worker does as much work as possible to manage the state of the job it is running. As a result, with a couple of minor exceptions, the leader process never needs to write or update the state of a job within the job-store. For example, when a job is complete and has no further successors the responsible worker deletes the job from the job-store, marking it complete. The leader then only has to check for the existence of the file when it receives a signal from the batch-system to know that the job is complete. This off-loading of state management is orthogonal to future parallelization of the leader.

#### 8.1.2 Job chaining

The scheduling of successor jobs is partially managed by the worker, reducing the number of individual jobs the leader needs to process. Currently this is very simple: if the there is a single next successor job to run and it’s resources fit within the resources of the current job and closely match the resources of the current job then the job is run immediately on the worker without returning to the leader. Further extensions of this strategy are possible, but for many workflows which define a series of serial successors (e.g. map sequencing reads, post-process mapped reads, etc.) this pattern is very effective at reducing leader workload.

#### 8.1.3 Preemptable node support

Critical to running at large-scale is dealing with intermittent node failures. Toil is therefore designed to always be resumable providing the job-store does not become corrupt. This robustness allows Toil to run on preemptible nodes, which are only available when others are not willing to pay more to use them. Designing workflows that divide into many short individual jobs that can use preemptable nodes allows for workflows to be efficiently scheduled and executed.

#### 8.1.4 Caching

Running bioinformatic pipelines often require the passing of large datasets between jobs. Toil caches the results from jobs such that child jobs running on the same node can directly use the same file objects, thereby eliminating the need for an intermediary transfer to the job store. Caching also reduces the burden on the local disks, because multiple jobs can share a single file. The resulting drop in I/O allows pipelines to run faster, and, by the sharing of files, allows users to run more jobs in parallel by reducing overall disk requirements.

To demonstrate the efficiency of caching, we ran an experimental internal pipeline on 3 samples from the TCGA Lung Squamous Carcinoma (LUSC) dataset. The pipeline takes the tumor and normal exome fastqs, and the tumor rna fastq and input, and predicts MHC presented neoepitopes in the patient that are potential targets for T-cell based immunotherapies. The pipeline was run individually on the samples on c3.8xlarge machines on AWS (60GB RAM,600GB SSD storage, 32 cores). The pipeline aligns the data to hg19-based references, predicts MHC haplotypes using PHLAT, calls mutations using 2 callers (MuTect and RADIA) and annotates them using SnpEff, then predicts MHC:peptide binding using the IEDB suite of tools before running an in-house rank boosting algorithm on the final calls.

To optimize time taken, The pipeline is written such that mutations are called on a per-chromosome basis from the whole-exome bams and are merged into a complete vcf. Running mutect in parallel on whole exome bams requires each mutect job to download the complete Tumor and Normal Bams to their working directories - An operation that quickly fills the disk and limits the parallelizability of jobs. The script was run in Toil, with and without caching, and Figure 2 shows that the workflow finishes faster in the cached case while using less disk on average than the uncached run. We believe that benefits of caching arising from file transfers will be much higher on magnetic disk-based storage systems as compared to the SSD systems we tested this on.

**Fig. 8.2: Figure 2:**
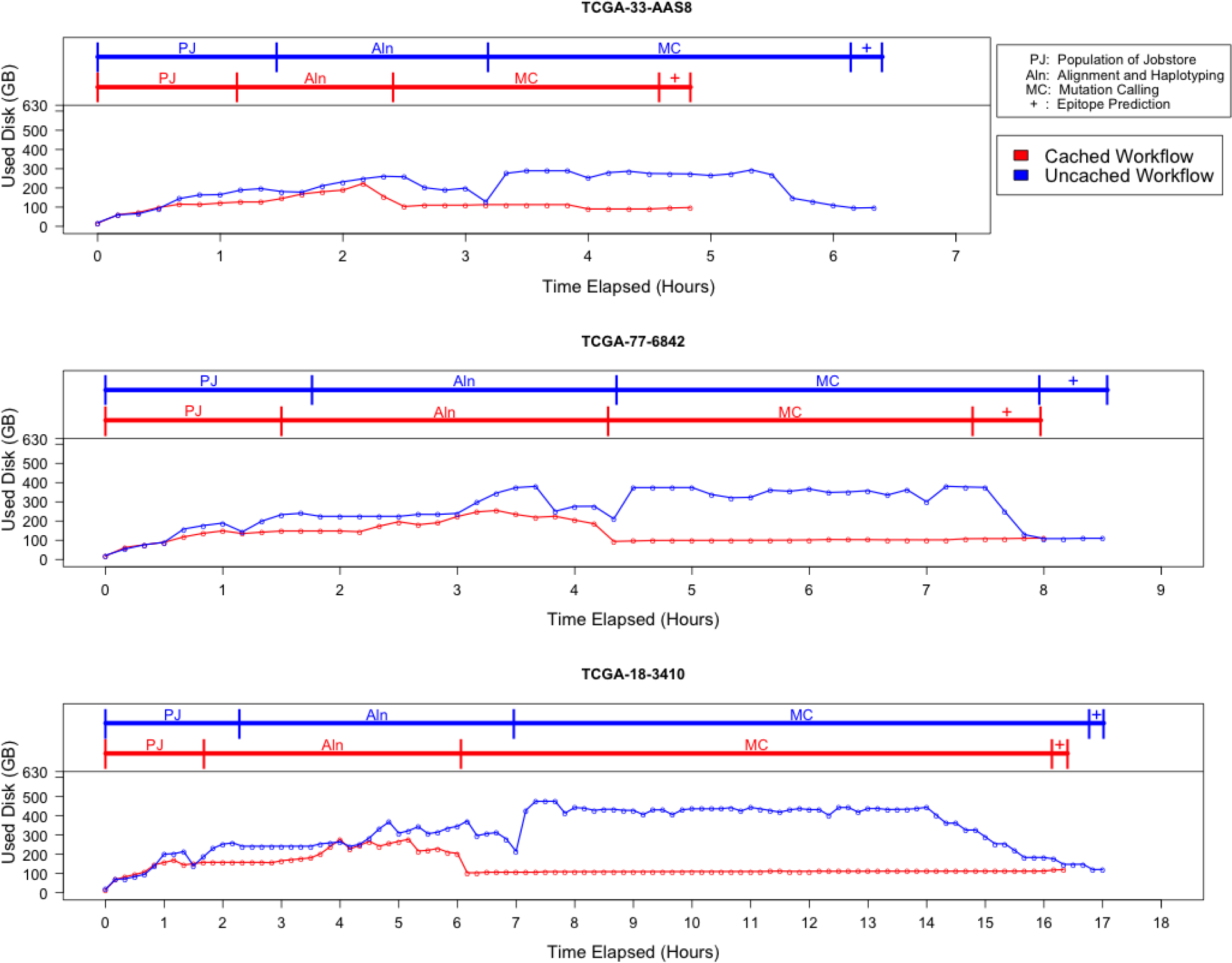
Efficiency gain from caching. The lower half of each plot describes the disk used by the pipeline recorded every 10 minutes over the duration of the pipeline, and the upper half shows the corresponding stage of the pipeline that is being processed. Since jobs requesting the same file shared the same inode, the effective load on the disk is considerably lower than in the uncached case where every job downloads a personal copy of every file it needs. We see that in all cases, the uncached run uses almost 300-400GB more that the uncached run in the resource heavy mutation calling step. We also see a benefit in terms of wall time for each stage since we eliminate the time taken for file transfers.

## THE BATCH SYSTEM INTERFACE

The batch system interface is used by Toil to abstract over different ways of running batches of jobs, for example GridEngine, Mesos, Parasol and a single node. The `toil.batchSystems.abstractBatchSystem.AbstractBatchSystem` API is implemented to run jobs using a given job management system, e.g. Mesos.

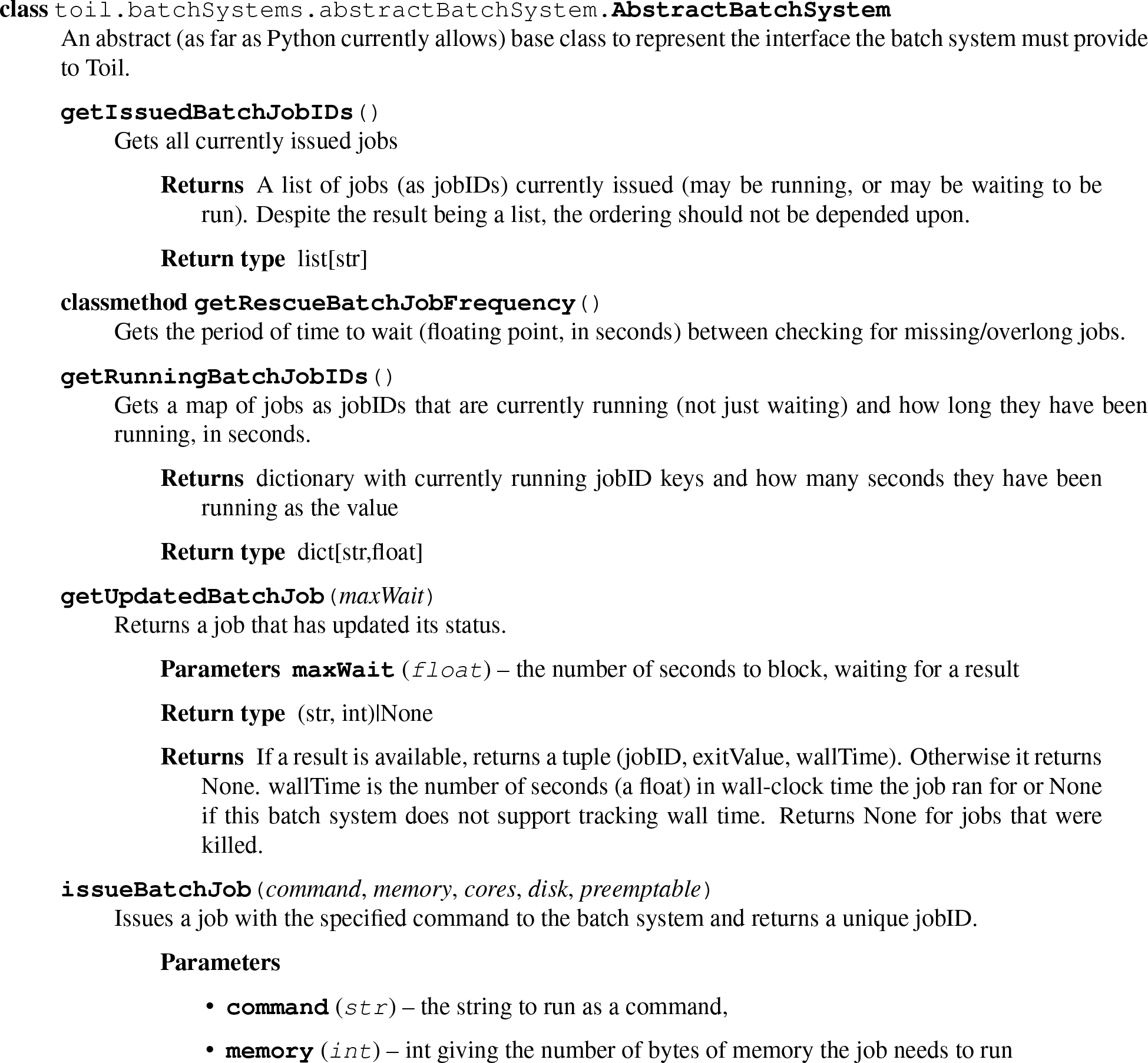

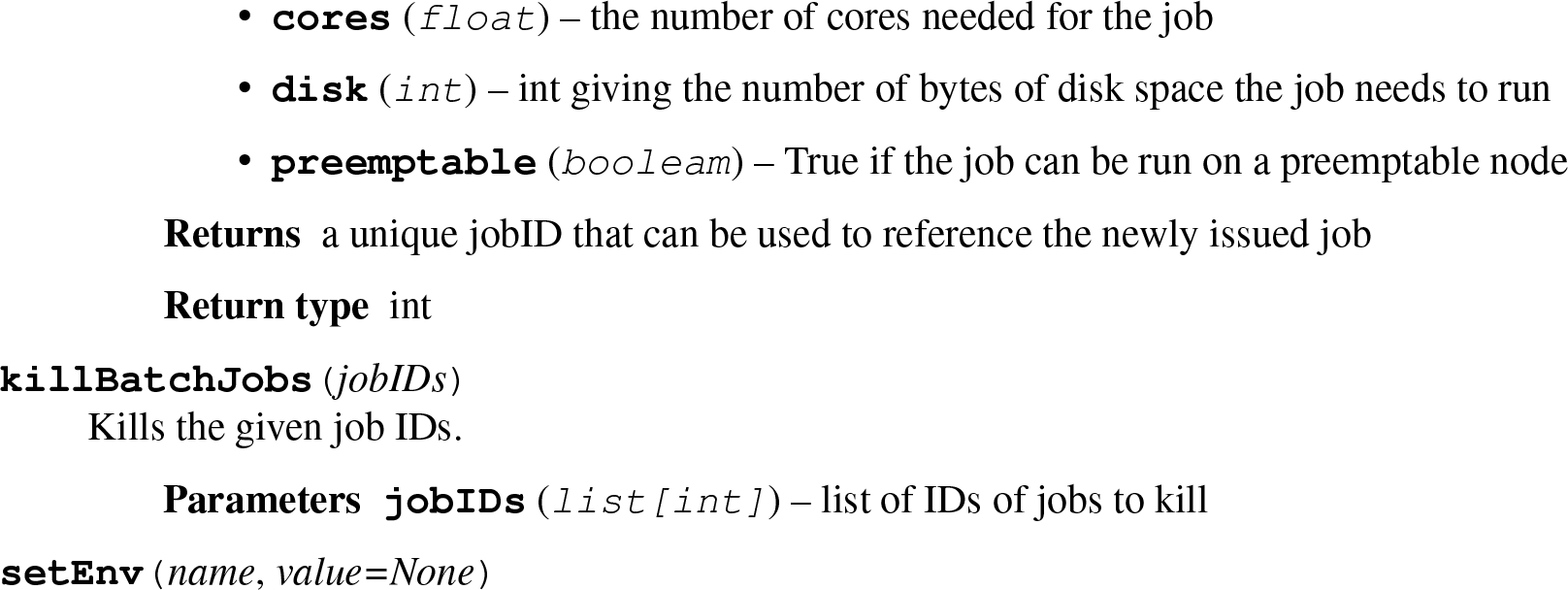

## THE JOB STORE INTERFACE

The job store interface is an abstraction layer that that hides the specific details of file storage, for example standard file systems, S3, etc. The `toil.jobStores.abstractJobStore.AbstractJobStore` API is implemented to support a give file store, e.g. S3. Implement this API to support a new file store.

## SRC

### 11.1 toil package

#### 11.1.1 Subpackages

**toil.batchSystems package Subpackages**

**toil.batchSystems.mesos package Subpackages**

**toil.batchSystems.mesos.test package**

**Module contents**

**Submodules**

**toil.batchSystems.mesos.batchSystem module**

**toil.batchSystems.mesos.conftest module**

**toil.batchSystems.mesos.executor module**

**Module contents**

**Submodules**

**toil.batchSystems.abstractBatchSystem module**

**toil.batchSystems.gridengine module**

**toil.batchSystems.jobDispatcher module**

**toil.batchSystems.lsf module**

~~~
toil.batchSystems.lsf.**bsub** (*bsubline*)
toil.batchSystems.lsf.**getjobexitcode**(*lsfJobID*)
toil.batchSystems.lsf.**prepareBsub**(*cpu*, *mem*)
~~~

**toil.batchSystems.parasol module**

**toil.batchSystems.parasolTestSupport module**

**module toil.batchSystems.singleMachine**

**toil.batchSystems.slurm module**

**Module contents**

**toil.cwl package Submodules**

**toil.cwl.conftest module**

**toil.cwl.cwltoil module**

**Module contents**

**toil.jobStores package**

**Subpackages**

**toil.jobStores.aws package**

**Submodules**

**toil.jobStores.aws.jobStore module**

**toil.jobStores.aws.utils module**

~~~
toil.jobStores.aws.utils.**bucket_location_to_http_url**(*location*)
toil.jobStores.aws.utils.**bucket_location_to_region**(*location*)
toil.jobStores.aws.utils.**connection_reset**(*e*)
toil.jobStores.aws.utils.**monkeyPatchSdbConnection**(*sdb*)
toil.jobStores.aws.utils.**no_such_sdb_domain**(*e*)
toil.jobStores.aws.utils.**region_to_bucket_location**(*region*)
toil.jobStores.aws.utils.**retry_s3**(*delays*=*(0*, *1*, *1*, *4*, *16*, *64)*, *timeout*=*300*, *predicate*=<*function retryable*_*s3*_*errors*>)
toil.jobStores.aws.utils.retry_sdb(*delays*=*(0*, *1*, *1*, *4*, *16*, *64)*, *timeout*=*300*, *predicate*=<*function retryable*_*sdb*_*errors*>)
toil.jobStores.aws.utils.**retryable_s3_errors**(*e*)
toil.jobStores.aws.utils.**retryable_sdb_errors**(*e*)
toil.jobStores.aws.utils.**retryable_ssl_error**(*e*)
toil.jobStores.aws.utils.**sdb_unavailable**(*e*)
~~~

**Module contents**

**Submodules**

**toil.jobStores.abstractJobStore module**

**toil.jobStores.azureJobStore module**

**toil.jobStores.conftest module**

**toil.jobStores.fileJobStore module**

**toil.jobStores.googleJobStore module**

**toil.jobStores.utils module**

~~~
toil.jobStores.utils.**never**(*exception*)
toil.jobStores.utils.**retry**(*delays*=*(0*, *1*, *1*, *4*, *16*, *64)*, *timeout*=*300*, *predicate*=<*function never*>)
~~~

Retry an operation while the failure matches a given predicate and until a given timeout expires, waiting a given amount of time in between attempts. This function is a generator that yields contextmanagers. See doctests below for example usage.

**Parameters**

- **delays** (`Iterable[float]`) an interable yielding the time in seconds to wait before each retried attempt, the last element of the iterable will be repeated.
- **timeout** (`float`) a overall timeout that should not be exceeded for all attempts together. This is a best-effort mechanism only and it won’t abort an ongoing attempt, even if the timeout expires during that attempt.
- **predicate** (`Callable[[Exception],bool]`) a unary callable returning True if another attempt should be made to recover from the given exception. The default value for this parameter will prevent any retries!

**Returns** a generator yielding context managers, one per attempt

**Return type** Iterator

Retry for a limited amount of time:

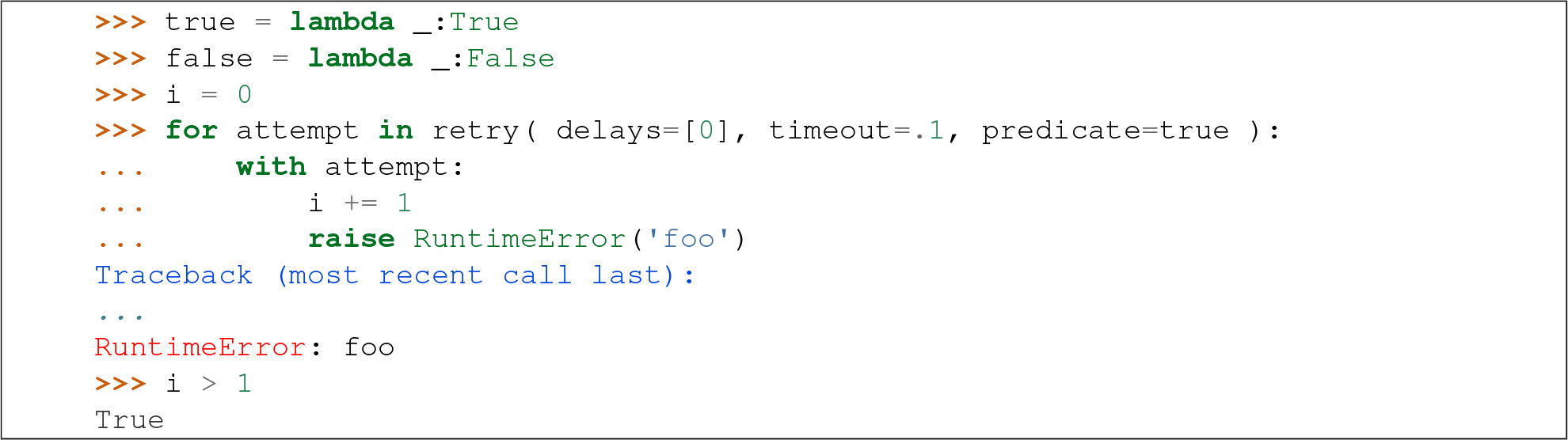

If timeout is 0, do exactly one attempt:

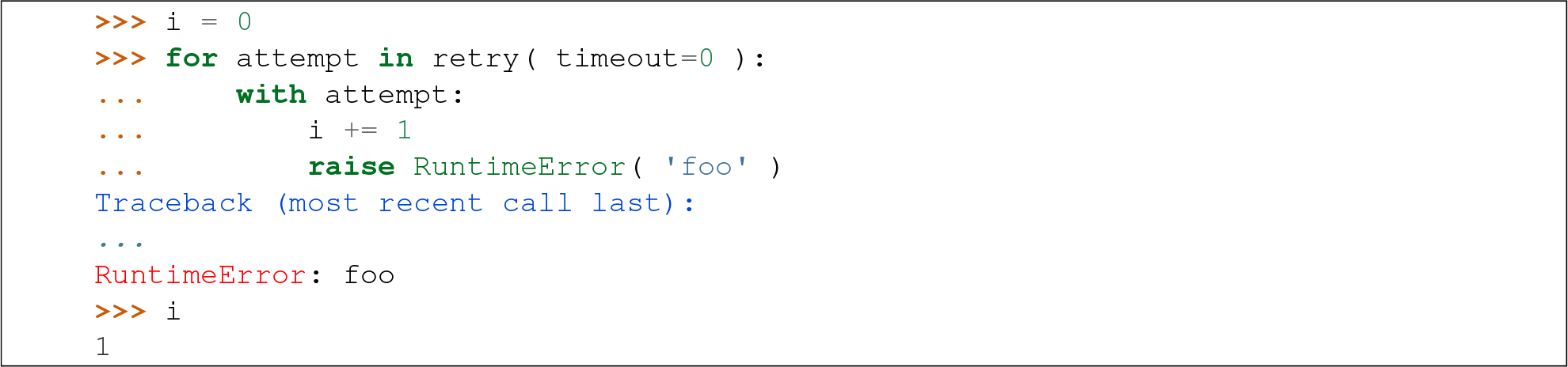

Don’t retry on success:

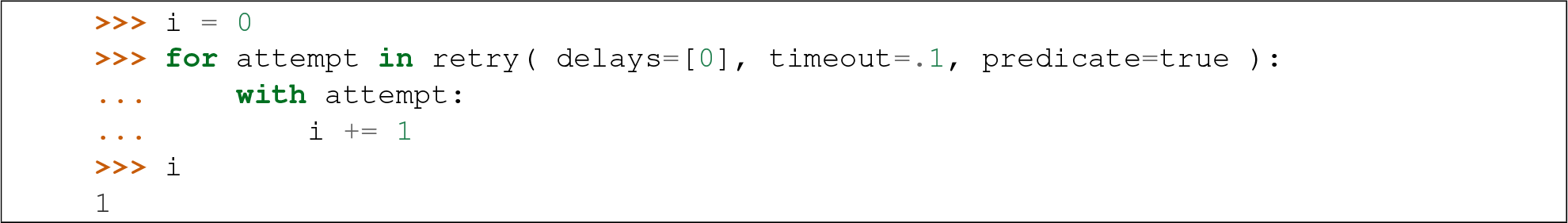

Don’t retry on unless predicate returns True:

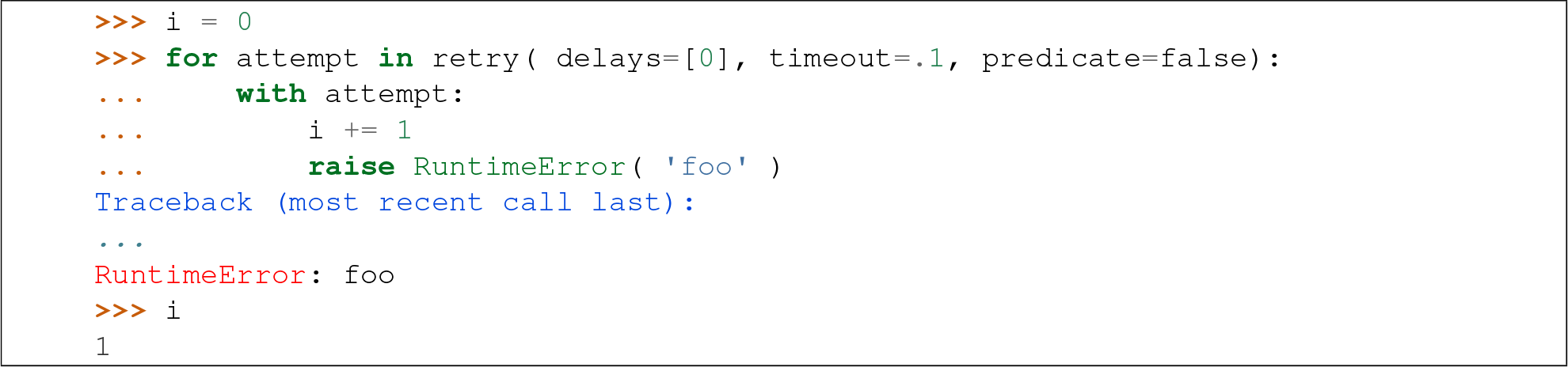

**Module contents**

**toil.lib package**

**Subpackages**

**toiLlib.encryption package**

**Submodules**

**toil.lib.encryption.conftest module**

**Module contents**

**Submodules**

**toil.lib.bioio module**

~~~
toil.lib.bioio.**absSymPath**(*path*)
like os.path.abspath except it doesn’t dereference symlinks
toil.lib.bioio.**addLoggingFileHandler**(*fileName*, *rotatingLogging*=*False*)
toil.lib.bioio.**addLoggingOptions**(*parser*)
toil.lib.bioio.**getBasicOptionParser**(*parser*=*None*)
toil.lib.bioio.**getLogLevelString**()
toil.lib.bioio.**getRandomAlphaNumericString**(*length*=*10*)
Returns a random alpha numeric string of the given length.
toil.lib.bioio.**getTempFile**(*suffix*=’‘, *rootDir*=*None*)
Returns a string representing a temporary file, that must be manually deleted
toil.lib.bioio.**getTotalCpuTime**()
Gives the total cpu time, including the children.
toil.lib.bioio.**getTotalCpuTimeAndMemoryUsage**()
Gives the total cpu time and memory usage of itself and its children.
toil.lib.bioio.**getTotalMemoryUsage**()
Gets the amount of memory used by the process and its children.
toil.lib.bioio.**logFile**(*fileName*, *printFunction*=<*bound method Logger.info of* <*logging.Logger object at 0x104513a50*>>)
Writes out a formatted version of the given log file
toil.lib.bioio.**logStream**(*fileHandle*, *shortName*, *printFunction*=<*bound method Logger.info of* <*logging.Logger object at 0x104513a50*>>)
Writes out a formatted version of the given log stream.
toil.lib.bioio.**makePublicDir**(*dirName*)
Makes a given subdirectory if it doesn’t already exist, making sure it is public.
toil.lib.bioio.**parseBasicOptions**(*parser*)
Setups the standard things from things added by getBasicOptionParser.
toil.lib.bioio.**setLogLevel**(*level*, *logger*=<*logging.RootLogger object*>)
Sets the log level to a given string level (like “INFO”). Operates on the root logger by default, but another logger can be specified instead.
toil.lib.bioio.**setLoggingFromOptions**(*options*)
Sets the logging from a dictionary of name/value options.
toil.lib.bioio.**system**(*command*)
A convenience wrapper around subprocess.check_call that logs the command before passing it on. The command can be either a string or a sequence of strings. If it is a string shell=True will be passed to subprocess. check_call.
~~~

**toil.lib.spark module**

**Module contents**

**toil.provisioners package**

**Subpackages**

**toil.provisioners.aws package**

**Submodules**

**toil.provisioners.aws.provisioner module**

**Module contents**

**Submodules**

**toil.provisioners.abstractProvisioner module**

**toil.provisioners.clusterScaler module**

**Module contents**

**toil.test package**

**Subpackages**

**toil.test.batchSystems package**

**Submodules**

**toil.test.batchSystems.batchSystemTest module**

**Module contents**

**toil.test.cwl package**

**Submodules**

**toil.test.cwl.conftest module**

**toil.test.cwl.cwlTest module**

**Module contents**

**toil.test.jobStores package**

**Submodules**

**toil.test.jobStores.jobStoreTest module**

**Module contents**

**toil.test.mesos package**

**Submodules**

**toil.test.mesos.helloWorld module**

**toil.test.mesos.mesosTest module**

**toil.test.mesos.stress module**

**Module contents**

**toil.test.provisioners package**

**Submodules**

**toil.test.provisioners.clusterScalerTest module**

**Module contents**

**toil.test.sort package**

**Submodules**

**toil.test.sort.lib module**

~~~
toil.test.sort.lib.**copySubRangeOfFile**(*inputFile*, *fileStart*, *fileEnd*, *outputFileHandle*)
Copies the range (in bytes) between fileStart and fileEnd to the given output file handle.
toil.test.sort.lib.**getMidPoint**(*file*, *fileStart*, *fileEnd*)
Finds the point in the file to split. Returns an int i such that fileStart <= i < fileEnd
toil.test.sort.lib.**merge**(*fileHandle1*, *fileHandle2*, *outputFileHandle*)
Merges together two files maintaining sorted order.
toil.test.sort.lib.**sort**(*file*)
Sorts the given file.
~~~

**toil.test.sort.sort module**

**toil.test.sort.sortTest module**

**Module contents**

**toil.test.src package**

**Submodules**

**toil.test.src.helloWorldTest module**

**toil.test.src.importExportFileTest module**

**toil.test.src.jobCacheEjectionTest module**

**toil.test.src.jobCacheTest module**

**toil.test.src.jobEncapsulationTest module**

**toil.test.src.jobFileStoreTest module**

**toil.test.src.jobServiceTest module**

**toil.test.src.jobTest module**

**toil.test.src.jobWrapperTest module**

**toil.test.src.multipartTransferTest module**

**toil.test.src.promisedRequirementTest module**

**toil.testsrc.promisesTest module**

**toil.test.src.realtimeLoggerTest module**

**toil.test.src.regularLogTest module**

**toil.test.src.resourceTest module**

**toil.test.src.retainTempDirTest module**

**toil.test.src.systemTest module**

**toil.test.src.toilContextManagerTest module**

**toil.test.src.userDefinedJobArgTypeTest module**

**Module contents**

**toil.test.utils package**

**Submodules**

**toil.test.utils.utilsTest module**

**Module contents**

Module contents

~~~
toil.test.**experimental**(**test_item**)
Use this to decorate experimental or brittle tests in order to skip them during regular builds.
toil.test.**file_begins_with**(*path*, *prefix*)
toil.test.**make_tests**(*generalMethod*, *targetClass*=*None*, ^**^*kwargs*)
This method dynamically generates test methods using the generalMethod as a template. Each generated function is the result of a unique combination of parameters applied to the generalMethod. Each of the parameters has a corresponding string that will be used to name the method. These generated functions are named in the scheme:
test_[generalMethodName]___[firstParamaterName]_[someValueName]__[secondParamaterName]_…
The arguments following the generalMethodName should be a series of one or more dictionaries of the form {str: type,…} where the key represents the name of the value. The names will be used to represent the permutation of values passed for each parameter in the generalMethod.
**Parameters**
**generalMethod** - A method that will be parametrized with values passed as kwargs. Note that the generalMethod must be a regular method.
**targetClass** - This represents the class to which the generated test methods will be bound. If no targetClass is specified the class of the generalMethod is assumed the target.
**kwargs** - a series of dictionaries defining values, and their respective names where each keyword is the name of a parameter in generalMethod.
~~~

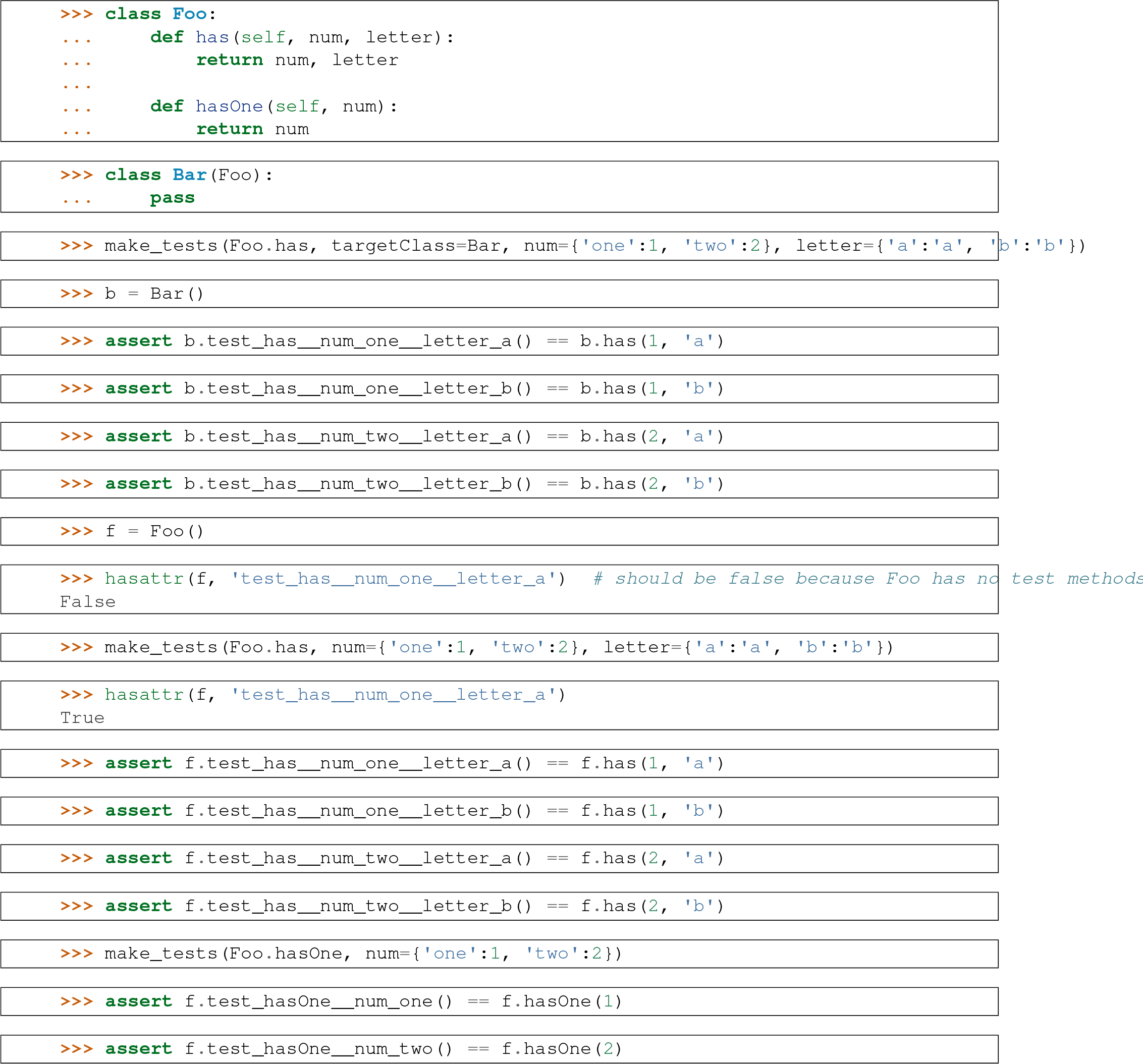

~~~
toil.test.**needs**_**aws**(*test*_*item*)
Use as a decorator before test classes or methods to only run them if AWS usable.
toil.test.**needs**_**azure**(*test*_*item*)
Use as a decorator before test classes or methods to only run them if Azure is usable.
toil.test.**needs**_**cwl**(*test*_*item*)
Use as a decorator before test classes or methods to only run them if CWLTool is installed and configured.
toil.test.**needs**_**encryption**(*test*_*item*)
Use as a decorator before test classes or methods to only run them if PyNaCl is installed and configured.
toil.test.**needs**_**google**(*test*_*item*)
Use as a decorator before test classes or methods to only run them if Google Storage usable.
toil.test.**needs**_**gridengine**(*test*_*item*)
Use as a decorator before test classes or methods to only run them if GridEngine is installed.
toil.test.**needs**_**mesos**(*test*_*item*)
Use as a decorator before test classes or methods to only run them if the Mesos is installed and configured.
toil.test.**needs**_**parasol**(*test*_*item*)
Use as decorator so tests are only run if Parasol is installed.
toil.test.**needs**_**slurm**(*test*_*item*)
Use as a decorator before test classes or methods to only run them if Slurm is installed.
toil.test.**needs**_**spark**(*test*_*item*)
Use as a decorator before test classes or methods to only run them if Spark is usable.
toil.test.**tempFileContaining**(^*^*args*, ^**^*kwds*)
~~~

**toil.utils package**

**Submodules**

**toil.utils.toilClean module**

**toil.utils.toilKill module**

Kills any running jobs trees in a rogue toil.

~~~
toil.utils.toilKill.**main**()
~~~

**toil.utils.toilMain module**

~~~
toil.utils.toilMain.**loadModules**()
toil.utils.toilMain.**main**()
toil.utils.toilMain.**printHelp**(*modules*)
~~~

**toil.utils.toilStats module**

Reports data about the given toil run.

~~~
toil.utils.toilStats.**buildElement**(*element*, *items*, *itemName*)
Create an element for output.
toil.utils.toilStats.**checkOptions**(*options*, *parser*)
Check options, throw parser.error() if something goes wrong
toil.utils.toilStats.**computeColumnWidths**(*job*_*types*, *worker*, *job*, *options*)
Return a ColumnWidths() object with the correct max widths.
toil.utils.toilStats.**createSummary**(*element*, *containingItems*, *containingItemName*, *getFn*)
toil.utils.toilStats.**decorateSubHeader**(*title*, *columnWidths*, *options*)
Add a marker to the correct field if the TITLE is sorted on.
toil.utils.toilStats.**decorateTitle**(*title*, *options*)
Add a marker to TITLE if the TITLE is sorted on.
toil.utils.toilStats.**get**(*tree*, *name*)
Return a float value attribute NAME from TREE.
toil.utils.toilStats.**getStats**(*options*)
**Collect and return the stats and config data.**
toil.utils.toilStats.**initializeOptions**(*parser*)
toil.utils.toilStats.**main**()
Reports stats on the workflow, use withstats option to toil.
toil.utils.toilStats.**padStr**(*s*, *field*=*None*)
Pad the begining of a string with spaces, if necessary.
toil.utils.toilStats.**prettyMemory**(*k*, *field*=*None*, *isBytes*=*False*)
Given input k as kilobytes, return a nicely formatted string.
toil.utils.toilStats.**prettyTime**(*t*, *field*=*None*)
Given input t as seconds, return a nicely formatted string.
toil.utils.toilStats.**printJson**(*elem*)
Return a JSON formatted string
toil.utils.toilStats.**processData**(*config*, *stats*, *options*)
toil.utils.toilStats.**refineData**(*root*, *options*)
walk down from the root and gather up the important bits.
toil.utils.toilStats.**reportData**(*tree*, *options*)
toil.utils.toilStats.**reportMemory**(*k*, *options*, *field*=*None*, *isBytes*=*False*)
*Given k kilobytes, report back the correct format as string.*
toil.utils.toilStats.**reportNumber**(*n*, *options*, *field*=*None*)
Given n an integer, report back the correct format as string.
toil.utils.toilStats.**reportPrettyData**(*root*, *worker*, *job*, *job*_*types*, *options*)
print the important bits out.
toil.utils.toilStats.**reportTime**(*t*, *options*, *field*=*None*)
Given t seconds, report back the correct format as string.
toil.utils.toilStats.**sortJobs**(*jobTypes*, *options*)
Return a jobTypes all sorted.
toil.utils.toilStats.**sprintTag**(*key*, *tag*, *options*, *columnWidths*=*None*)
Generate a pretty-print ready string from a JTTag().
toil.utils.toilStats.**updateColumnWidths**(*tag*, *cw*, *options*)
Update the column width attributes for this tag’s fields.
~~~

**toil.utils.toilStatus module**

**Module contents**

#### 11.1.2 Submodules

#### 11.1.3 toil.common module

~~~
toil.common.**addOptions**(*parser*, *config*=<*toil.common.Config object*>)
Adds toil options to a parser object, either optparse or argparse.
toil.common.**cacheDirName**(*workflowID*)
**Returns** Name of the cache directory.
toil.common.**parseSetEnv**(*l*)
Parses a list of strings of the form “NAME=VALUE” or just “NAME” into a dictionary. Strings of the latter from will result in dictionary entries whose value is None.
**Return type** dict[str,str]
~~~

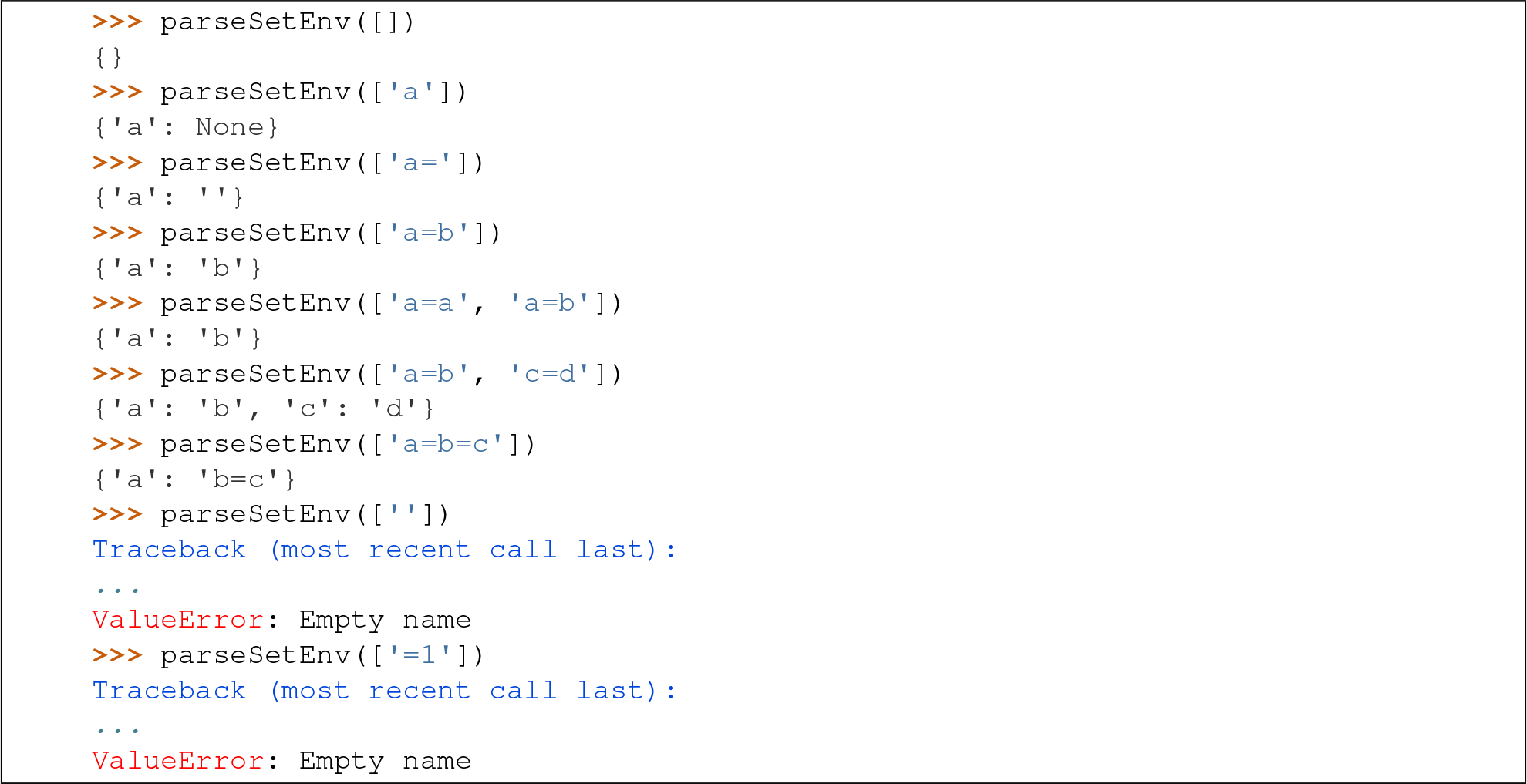

#### 11.1.4 toil.job module

#### 11.1.5 toil.jobWrapper module

#### 11.1.6 toil.leader module

The leader script (of the leader/worker pair) for running jobs.

~~~
toil.leader.**innerLoop**(*jobStore*, *config*, *batchSystem*, *toilState*, *jobBatcher*, *serviceManager*, *statsAndLogging*)
~~~

Parameters

- jobStore (*`toil.jobStores.abstractJobStore.AbstractJobStore`*)
- config (*`toil.common.Config`*)
- batchSystem (*`toil.batchSystems.abstractBatchSystem.AbstractBatchSystem`*)
- toilState (*`ToilState`*)
- jobBatcher (*`JobBatcher`*)
- serviceManager (*`ServiceManager`*)
- statsAndLogging (*`StatsAndLogging`*)

~~~
toil.leader.**mainLoop**(*config*, *batchSystem*, *provisioner*, *jobStore*, *rootJobWrapper*, *jobCache*=*None*)
~~~

This is the main loop from which jobs are issued and processed.

If jobCache is passed, it must be a dict from job ID to pre-existing JobWrapper objects. Jobs will be loaded from the cache (which can be downloaded from the jobStore in a batch).

**Raises** toil.leader.FailedJobsException if at the end of function their remain failed jobs

**Returns** The return value of the root job’s run function.

**Return type** Any

#### 11.1.7

Implements a real-time UDP-based logging system that user scripts can use for debugging.

#### 11.1.8 toil.resource module

#### 11.1.9 toil.toilState module

#### 11.1.10 toil.version module

#### 11.1.11 toil.worker module

~~~
toil.worker.**main**()
toil.worker.**nextOpenDescriptor**()
**Gets the number of the next available file descriptor.**
~~~

#### 11.1.12 Module contents

~~~
toil.**inVirtualEnv**()
toil.**physicalMemory**(^*^*args*)
~~~

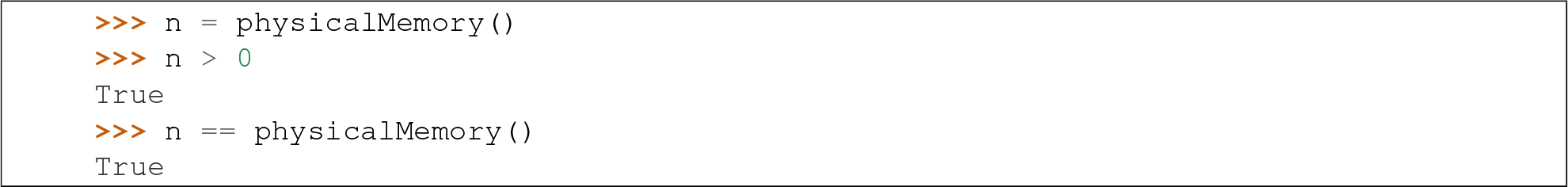

~~~
toil.**resolveEntryPoint**(*entryPoint*)
Returns the path to the given entry point (see setup.py) that *should* work on a worker. The return value may be
an absolute or a relative path.
toil.**toilPackageDirPath**()
Returns the absolute path of the directory that corresponds to the top-level toil package. The return value is guaranteed to end in ‘/toil’.
~~~

## INDICES AND TABLES

- genindex
- modindex
- search

## PYTHON MODULE INDEX

~~~
toil, 62
toil.batchSystems, 50
toil.batchSystems.abstractBatchSystem, 50
toil.batchSystems.gridengine, 50
toil.batchSystems.jobDispatcher, 50
toil.batchSystems.lsf, 50
toil.batchSystems.mesos, 49
toil.batchSystems.mesos.conftest, 49
toil.batchSystems.mesos.test, 49
toil.batchSystems.parasol, 50
toil.batchSystems.parasolTestSupport, 50
toil.batchSystems.singleMachine, 50
toil.batchSystems.slurm, 50
toil.common, 61
toil.cwl, 50
toil.cwl.conftest, 50
toil.jobStores, 53
toil.jobStores.aws, 51
toil.jobStores.aws.utils, 51
toil.jobStores.conftest, 51
toil.jobStores.utils, 51
toil.jobWrapper, 61
toil.leader, 61
toil.lib, 54
toil.lib.bioio, 53
toil.lib.encryption, 53
toil.lib.encryption.conftest, 53
toil.provisioners, 54
toil.provisioners.abstractProvisioner, 54
toil.provisioners.aws, 54
toil.provisioners.clusterScaler, 54
toil.realtimeLogger, 62
toil.test, 57
toil.test.batchSystems, 55
toil.test.cwl, 55
toil.test.cwl.conftest, 55
toil.test.cwl.cwlTest, 55
toil.test.jobStores, 55
toil.test.mesos, 55
toil.test.provisioners, 55
toil.test.provisioners.clusterScalerTest, 55
toil.test.sort, 56
toil.test.sort.lib, 56
toil.test.src, 57
toil.test.src.systemTest, 57
toil.test.utils, 57
toil.toilState, 62
toil.utils, 61
toil.utils.toilKill, 59
toil.utils.toilMain, 59
toil.utils.toilStats, 59
toil.version, 62
toil.worker, 62
~~~

